# Electrophysiological normative responses to emotional, neutral, and cigarette-related images

**DOI:** 10.1101/2022.04.11.487896

**Authors:** Francesco Versace, Nicola Sambuco, Menton M. Deweese, Paul M. Cinciripini

## Abstract

To create reproducible emotional probes, affective scientists rely on sets of standardized pictures that are normed using subjective ratings of valence and emotional arousal. Yet, to investigate psychophysiological emotional responses, it might be more appropriate to select pictures using normative neurophysiological responses rather than normative subjective ratings. Here, we provide electrophysiological normative responses for 323 emotional pictures (215 from the IAPS) covering a wide range of categories (erotica, romantic, appetizing foods, landscapes, people engaged in mundane activities, household objects, disgusting objects, accidents, sad people, violence, mutilations, and cigarette-related contents). Event-related potentials (ERPs) and subjective ratings of pleasure and emotional arousal were collected from 763 individuals (52% females, 41% white) aged between 18 and 65 (mean = 43). For each image, the mean amplitude of the late positive potential (LPP, an electrophysiological index of motivational relevance) and the mean subjective ratings of valence and arousal were calculated. We validated our procedure by showing that the subjective ratings of valence and arousal from this sample were highly correlated to the IAPS’ published norms (Pearson r=.97 for pleasure and r=.82 for emotional arousal). LPP responses and subjective ratings of emotional arousal also were correlated (Pearson r = .61), but some categories that participants reported being significantly more arousing than neutral (i.e., food, landscapes, and unpleasant objects) did not evoke LPPs significantly different from those evoked by neutral pictures. Researchers interested in probing the brain’s affective systems can use these electrophysiological normative responses to create emotional probes that evoke reliable neuroaffective responses.

## 1. INTRODUCTION

Measuring neurobehavioral responses to affectively charged pictures is one of the most frequently used experimental paradigms in affective neuroscience (Bradley & Lang, 2018). Experimental control and reproducibility across laboratories are ensured by the availability of picture sets that include hundreds of standardized pictures. Some of these picture sets cover a wide array of emotional contents (e.g., the International Affective Picture System [IAPS], Lang et al., 2008; the Nencki Affective Picture System [NAPS], Marchewka et al., 2014; the EmoPics, Wessa et al., 2010), while others focus on specific contents such as food (e.g., the open library of affective foods [OLAF], Miccoli et al., 2016); Food-pics, Blechert et al., 2014), drugs (SmoCuDA, Manoliu et al., 2021; Macatee et al., 2021; Methamphetamine and Opioid Cue Database [MOCD], Ekhtiari et al., 2020), trauma-related pictures (Trauma-related Affective Picture Set Muenster [TRAPS-M], Neumeister et al., 2016), or panic-related pictures (Panic-related Picture Set Muenster [PAPS-M], Feldker et al. 2017), to name a few.

The key feature of these sets is that they provide normative ratings of valence and arousal for each picture. Considering emotions as the result of the activation of fundamental motivational circuits that have evolved to protect and sustain life (Lang & Bradley, 2010), valence ratings indicate which motivational system is predominantly active (appetitive or defensive) while arousal ratings indicate the level of motivational activation (Bradley et al., 2001a). Therefore, using the available normative ratings of valence and arousal, researchers can create reproducible sets of pictures to evoke emotional reactions varying in intensity and involving both pleasant and unpleasant affect.

However, several studies have consistently demonstrated that cultural and social desirability biases skew subjective evaluations, and selecting experimental materials considering exclusively self-reported ratings of valence and arousal can complicate the interpretation of physiological findings. For example, studies assessing event-related potentials (ERPs) to emotional and neutral pictures have demonstrated that self-reports do not always predict brain responses (Schupp et al., 2000; Weinberg & Hajcak, 2010). Moreover, while images with sport and adventure contents are rated similarly to erotic images in terms of emotional valence, erotic images prompt significantly stronger physiological responses than sport and adventure contents (Bradley et al., 2001a). In addition, despite the presence of sex differences in ratings of erotic pictures and mutilations, females and males often show similar physiological engagement to these contents when skin conductance response is measured (Bradley et al., 2001b).

While self-reports can provide an important window into an individual’s subjective experience, these reports cannot provide information regarding the extent of neurophysiological emotional engagement. Thus, we propose that neuroaffective responses should be used to integrate the information obtained by self-reports to enhance the stimulus selection process. Toward this goal, we used the Late Positive Potential (LPP) as an objective neurophysiological index of emotional engagement. The LPP, a centroparietal positive ERP component that increases in response to emotional (pleasant or unpleasant) compared to neutral pictures, is a robust physiological measure of motivational relevance that reflects the engagement of attentional resources by emotional stimuli and the activation of motivational systems (Bradley, 2009; Codispoti et al., 2006; Cuthbert et al., 2000; De Cesarei & Codispoti, 2011; Ferrari et al., 2011; Weinberg & Hajcak, 2010). Previous studies have demonstrated that the LPP has high internal and test-retest reliability (Huffmeijer et al., 2014; Moran et al., 2013; Weinberg et al., 2021), and it is robust to manipulations affecting the perceptual characteristics of the images (e.g., presentation time, color, or repeated presentations; Codispoti et al., 2009, 2012). In addition, the LPP has been successfully used to assess individual differences in emotional reactivity in various populations (e.g., Cofresí et al., 2022; Kujawa et al., 2015; Versace et al., 2019).

The primary aim of the current study is to report electrophysiological normative responses for naturalistic pictures spanning pleasant (erotica, romantic, food-related, and landscapes), neutral (people engaged in mundane activities and household objects), unpleasant (mutilation, violence, sadness and accidents), and cigarette-related contents. In a novel picture-based analysis we pooled together data from 763 individuals and 323 pictures from three published studies that used a passive viewing paradigm (Frank et al., 2020; Versace et al., 2012, 2016). Of these images, 215 pictures were selected from the IAPS, 36 were images that had similar contents to those included in the IAPS and 72 were cigarette-related images from existing picture sets (Carter et al., 2006; Gilbert et al., 1999).

We first demonstrate the validity of our results by comparing the normative self-reported ratings of pleasure and arousal for the 215 IAPS pictures (IAPS normative ratings) with the ratings of the same pictures obtained in the current study from a community sample in Houston, Texas (HTX ratings), and then we present the LPP response evoked by these pictures (1) across the entire sample, (2) separately for males and females, and then (3) separately for smokers and non-smokers. By sharing these electrophysiological normative responses (accessible via Supplementary materials), our goal is to provide a neuroaffective measure to integrate the information obtained by self-reports and contribute to improve experimental stimulus selection procedures.

## 2. METHODS

### 2.1 Study Participants

Participants were 763 individuals from the Houston metropolitan area that, between 2012 and 2017, volunteered in studies that enrolled exclusively smokers interested in quitting (Versace et al., 2012), smokers interested in quitting and non-smokers controls (Frank et al., 2020), or enrolled exclusively non-smokers (Versace et al., 2016). Inclusion and exclusion criteria were similar across studies and aimed at ensuring recruitment of healthy individuals free from psychiatric disorders. Pregnant or lactating women were excluded from the studies. All participants signed a consent form before starting the study. Table 1 shows the demographic characteristics of the sample, and supplementary Figure S1 shows the age distribution of the participants.

**Table 1.** Characteristics of the samples included in the current analyses.

| Study | Number of subjects | Mean Age | Age range | % Females | Race |  |  | % Smoker |
| --- | --- | --- | --- | --- | --- | --- | --- | --- |
|  |  |  |  |  | % White | % Black | % Other * |  |
| EEG1 | 242 | 45 | 19-65 | 37 | 65 | 22 | 13 | 100 |
| EEG2 | 340 | 45 | 18-65 | 50 | 26 | 69 | 5 | 70 |
| EEG3 | 181 | 35 | 20-55 | 76 | 33 | 46 | 21 | 0 |
| Total | 763 | 43 | 18-65 | 52 | 41 | 47 | 12 | 63 |

*Note:* EEG1: Versace et al., 2012; EEG2: Frank et al., 2020; EEG3: Versace et al., 2016; \* Non-white other than Black, more than one race, or no response to the question.

### 2.2 Procedures

All procedures were approved by the MD Anderson Institutional Review Board. Procedures were similar across studies: At baseline (i.e., before participants were randomized to any treatment), we (1) collected self-report questionnaires about mood and other variables related to the studies’ specific scientific goals (the questionnaires differed across studies), (2) placed a 129-channel sensor array and continuously record EEG during a passive picture viewing task, (3) and, after removing the EEG sensors, collected subjective ratings of emotional valence and arousal using an electronic version of the Self-Assessment Manikin (Bradley & Lang, 1994). In all the studies (Frank et al., 2020; Versace et al., 2012, 2016), participants rated half of the pictures that they viewed during the EEG session.

### 2.3 Passive Picture Viewing Task

Irrespective of the study, participants viewed comparable slideshows that included approximately the same number of pictures (192 or 160) distributed across the same semantic categories: erotic scenes (pleasant, high arousing contents), romantic scenes, landscapes, food, neutral household objects, people involved in mundane activities, unpleasant objects (including a variety of unpleasant contents such as pollution or accidents), sad and violent scenes (henceforth referred to as violence, since this is the content with the largest number of exemplars), mutilations (unpleasant, high arousing contents), people smoking cigarettes, and cigarette-related objects (e.g., packs of cigarettes, ashtrays). Of the 323 available pictures, most of the emotional and neutral images were selected from the IAPS (N=219); additional scenes were selected from other sources including the internet to supplement categories with a limited number of stimuli. Cigarette-related pictures were selected from two existing stimulus sets (Carter et al., 2006; Gilbert & Rabinovich, 1999). The Supplement includes IAPS catalog numbers for each picture used in the current analysis, as well as coded numbers for the cigarette-related pictures and additional pictures downloaded from the internet (pictures are available upon request).

In each study, participants sat in a comfortable chair placed in front of a computer screen and were instructed to move as little as possible during the picture presentation to limit EEG artifacts. Images appeared on the screen every 3 to 5 seconds and remained visible for 4 seconds. During the picture presentation, no more than two pictures from the same category were presented sequentially. Every 5 minutes, a short pause allowed the participant to relax.

### 2.4 EEG data collection

EEG was recorded using a 129-channel Geodesic Sensor Net, amplified with an AC-coupled high-input impedance amplifier (Geodesic EEG System 200, Electrical Geodesics Inc., Eugene, OR, USA), and referenced to the Cz electrode site. The sampling rate was 250 Hz, and data were filtered online by using 0.1-Hz high-pass and 100-Hz low-pass filters. Scalp impedance of each sensor was kept below 50 KΩ, as suggested by the manufacturer.

### 2.5 SAM rating

Following EEG data collection, the sensor net was removed, and the participants completed the valence and arousal ratings using an electronic version of the Self-Assessment Manikin (Lang, 1980; Bradley & Lang, 1994). Prior to the rating procedure, a trained research assistant read to each participant the rating directions according to the procedure described in the IAPS manual (Lang et al., 2008). Then, two practice trials were presented to familiarize participants with the rating procedure. The research assistant remained in the room to answer any questions during the practice trials. If the participant asked how they should rate a particular picture, the research assistant reiterated that there were no right or wrong answers, and that it was up to the participant to rate the picture according to the feelings that the picture prompted, keeping in mind the anchor adjectives at the two extremes of the SAM valence and arousal scales. After the short practice, participants rated half of the pictures that they viewed in each category during the EEG assessment. Pictures were presented for 3 seconds in pseudo random order (no more than two pictures from the same picture category). Then, the valence scale appeared and remained visible until the participant clicked on one of the available ratings boxes, followed by the arousal scale. After one second, the next picture appeared, and the rating procedure was repeated.

### 2.6 EEG Data Reduction

Offline data reduction followed a standard pipeline that included low-pass filtering (30Hz), interpolation of broken channels, re-referencing to the average reference, and eye blink correction (using a spatial filtering method as implemented in BESA (v5.1.8.10; MEGIS Software GmbH, Gräfelfing, Germany). Data were then exported to Brain Vision Analyzer (Brain Products GmbH), where the EEG data were segmented into 900-millisecond epochs starting 100 milliseconds before picture onset. Baseline correction was applied by using the data from the 100-millisecond segment preceding picture onset. Artifacts were identified in each segment and each channel using the following criteria: EEG amplitude above 100 or below −100 μV, absolute voltage difference between any two data points within the segment larger than 100 μV, voltage difference between two contiguous data points above 25 μV or variation of less than 0.5 μV for more than 100 ms. Channels contaminated by artifacts in more than 40% of the segments were interpolated using spherical splines. All stimulus triggers were recoded using identifiers unique to the pictures that the participants saw during the session. Then, we pooled the voltage from 10 central and parietal sensors (see Results section for additional details) and, using the same criteria outlined above, artifacts affecting this pooled channel were identified and eliminated from the subsequent analyses. This procedure yielded one segment for every picture that the participant saw.

These segments were used to compute the average ERPs for each image, and based on previous studies (Frank et al., 2020; Minnix et al., 2013; Versace et al., 2012, 2016, 2019) the LPP was calculated for each picture by averaging the voltage recorded between 400 and 800 ms post picture onset. These LPP values were used as dependent variables in the statistical analyses. For each picture, LPPs were averaged across subjects in different ways: (1) across the whole sample, (2) separately for males and females and (3) separately for non-smokers and smokers.

Since stimuli varied slightly across studies or were not always presented an equal number of times, the average LPP value for each picture varied in the number of participants that contributed to the mean. The image with the least number of participants that contributed to the LPP mean resulted from the average of 56 subjects, while the image with the largest amount of contributing data resulted from the average of 543 subjects. The average number of participants contributing to the LPP mean for a picture was 312 subjects.

### 2.7 SAM Data Reduction

For each participant, we exported the ratings of valence and arousal associated with each picture and averaged separately the ratings of valence and arousal attributed to each picture. Each picture was rated by an average of 160 participants (range: 47-323)^1^. For each image we computed average ratings of emotional valence and arousal across the whole sample, separately for males and females, and separately for smokers and non-smokers.

### 2.8 Data analyses

First, to ensure the validity of our procedures, we compared the subjective ratings of emotional valence and arousal for pleasant, neutral, and unpleasant contents that we collected in our studies with the normative ratings reported in the IAPS. We correlated the IAPS normative ratings of valence and arousal with the corresponding values obtained from our sample, and then we assessed the pattern of differences across categories (MU, VI, OU, NO, NP, FD, LA, RO, ER) in separate ANOVAs for the IAPS normative ratings and HTX ratings. We assessed differences between the two ratings sets with one-way ANOVAs, separately by valence and arousal. Lastly, we used separate ANOVAs to assess the effects of category in males and females, and between-sex differences for each picture category.

We used an identical approach to evaluate differences in LPP amplitude as a function of picture content: We followed up the ANOVA assessing the main effect of picture content with separate ANOVAs for males and females, and then we ran a between-sex comparison.

The primary analyses reported in the current study were performed using exclusively IAPS images. Then, we ran a second set of analyses including all images. The number of exemplars included in each analysis is reported in Table S1.

## 3. RESULTS

The Supplement includes mean valence and arousal ratings and mean LPP amplitudes for each picture included in the study.

### 3.1 Valence and arousal ratings for pleasant and neutral pictures

These analyses included only images from the IAPS and were primarily designed as a validation procedure to assess the similarity between the IAPS normative ratings and the newly compiled HTX ratings.

As illustrated in Figure 1A (left), we observed a high positive correlation between the valence ratings that the HTX participants attributed to the IAPS pictures and the normative ratings reported in the IAPS (r=.97, p<.001). We observed a significant main effect of category for both the IAPS normative ratings (F_8,206_ = 282.06, p<001) and HTX ratings (F_8,206_ = 272.70, p<001). For both sets of ratings, pleasant contents had the highest valence ratings, followed by neutral contents and then unpleasant contents. Pairwise comparisons are reported in Table 2, while Figure 1A (right) illustrates differences in valence ratings between the IAPS normative ratings and HTX ratings. A similar response pattern for valence ratings was observed in males and females (Figure S2) and, in line with many previous findings (e.g., Bradley et al., 2001b), males rated erotic images as more pleasant than females (IAPS normative: F_1,42_ = 44.26, p < .001; HTX ratings: F_1,42_ = 139.91, p < .001) and mutilations were rated as more unpleasant by females than males (IAPS normative: F_1,46_ = 35.06, p < .001; HTX ratings: F_1,46_ = 17.50, p < .001).

**Figure 1.**
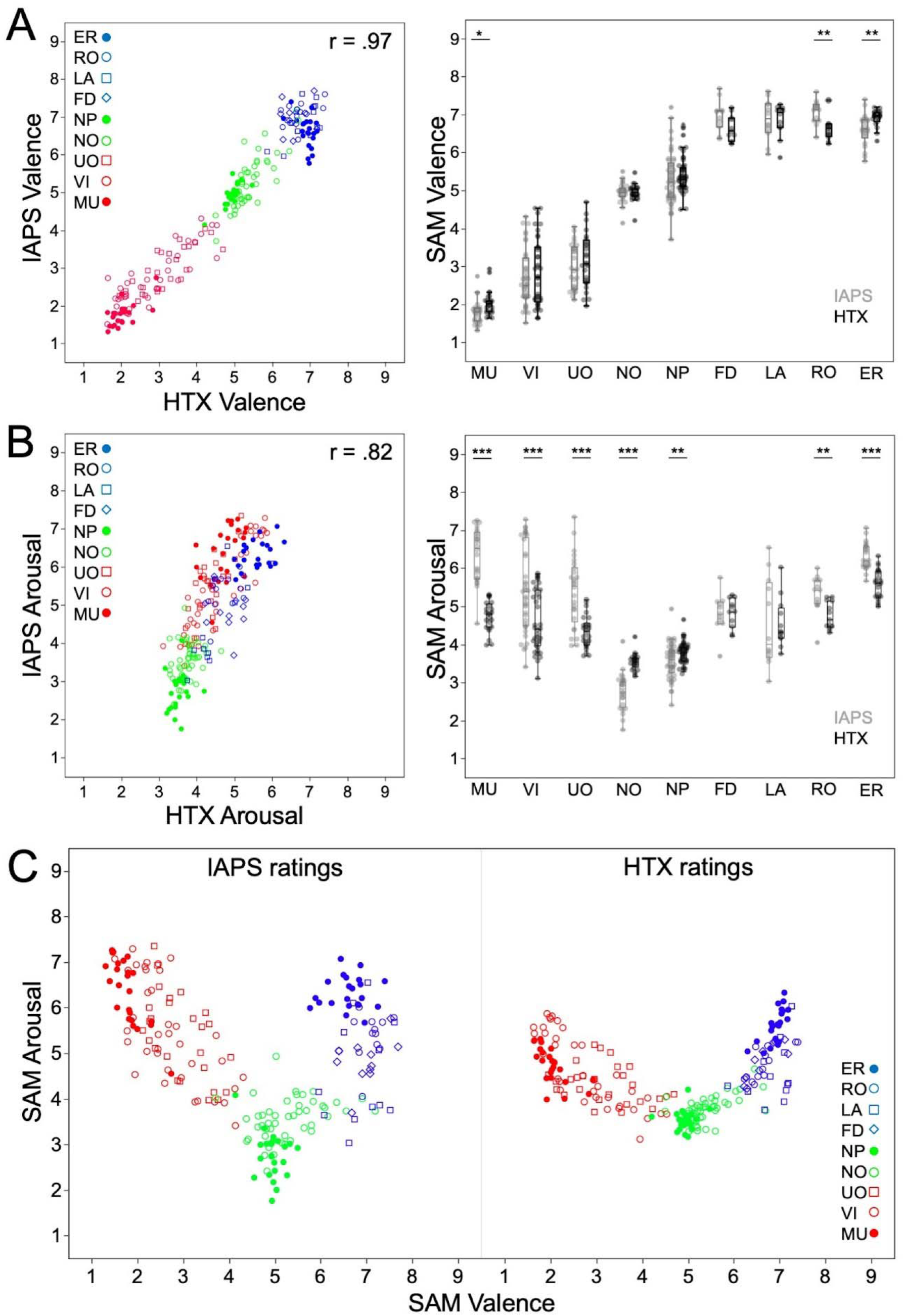
Comparison of SAM ratings between the IAPS normative ratings and HTX ratings for valence (A), arousal (B), and affective space (C). In A and B, the left panel illustrates the ratings correlations between IAPS normative ratings and HTX ratings, while the right panel illustrates differences based on categorical contents. Note. MU: Mutilations; VI: Violence; UO: Unpleasant Objects; NO: Neutral Objects; NP: Neutral People; FD: Food; LA: Landscapes; RO: Romance; ER: Erotica. Results of between-group comparison significant at ***p<.001, **p<.01, *p<.05.

**Table 2.** Tukey HSD pairwise comparisons (p<.05) for SAM rating of valence and arousal (separately reported for IAPS normative ratings and HTX ratings) and LPP*. **Note:** Categories which share at least one letter do not significantly differ (e.g., in the LPP column, Erotica and Mutilations do not differ from each other, and both differ from all other categories)*.

| Category | SAM ratings |  |  |  |  |  | LPP |
| --- | --- | --- | --- | --- | --- | --- | --- |
|  | Valence |  |  | Arousal |  |  |  |
|  | IAPS |  | HTX | IAPS |  | HTX |  |
| Erotica | A |  | A | A |  | A | A |
| Romance | A |  | A | B | C | B | B |
| Landscapes | A |  | A |  | C | B | D |
| Food | A |  | A | B | C | B | C D |
| Neutral People | B |  | B |  | D | C | D |
| Neutral Objects | B |  | C |  | E | C | D |
| Unpleasant Objects |  | C | D | B |  | B | C |
| Violence |  | C | D | B |  | B | B C |
| Mutilations |  | D | E | A |  | B | A |

We also observed a high positive correlation between the emotional arousal ratings that the HTX participants attributed to the IAPS pictures and the normative ratings reported in the IAPS (r=.82, p<.001; Figure 1B, left). A significant main effect of category was observed for both the IAPS normative ratings (F_8,206_ = 68.17, p<001) and HTX ratings (F_8,206_ = 42.19, p<001), where erotica and mutilations had the highest arousal ratings compared to other contents (pairwise comparisons are reported in Table 2). Between-sample comparisons revealed that many pleasant and unpleasant categories were rated significantly lower in arousal from HTX ratings compared to the IAPS normative ratings (Figure 1B, right). In line with valence ratings, males rated erotic pictures as more arousing than females (IAPS normative: F_1,42_ = 42.26, p < .001; HTX ratings: F_1,42_ = 49.62, p < .001), while females rated mutilations as more arousing than males (IAPS normative: F_1,46_ = 35.06, p < .001; HTX ratings: F_1,46_ = 18.93, p < .001).

Despite the differences in arousal ratings observed in the HTX sample relative to the IAPS normative ratings, the affective space resulting from integrating valence and arousal was similar in both the IAPS and the HTX. Figure 1C shows that in both the IAPS and the HTX sample, pictures rated as highly pleasant or highly unpleasant are associated with higher arousal ratings, resulting in a significant quadratic relationship between pleasure and arousal (IAPS normative ratings, quadratic trend: R^2^ = .56, p<.001; HTX ratings, quadratic trend: R^2^ = .66, p<.001).

### 3.2 Electrophysiological normative responses for pleasant and neutral pictures

These analyses included images from the IAPS and images added by the authors, for a total of 251 pictures (see Table S1).

#### 3.2.1 Grand-average ERPs

Figure 2 (left) illustrates the average ERPs, with the average computed over participants for each picture in a first step and then for pleasant, neutral, and unpleasant contents as a second step.

**Figure 2.**
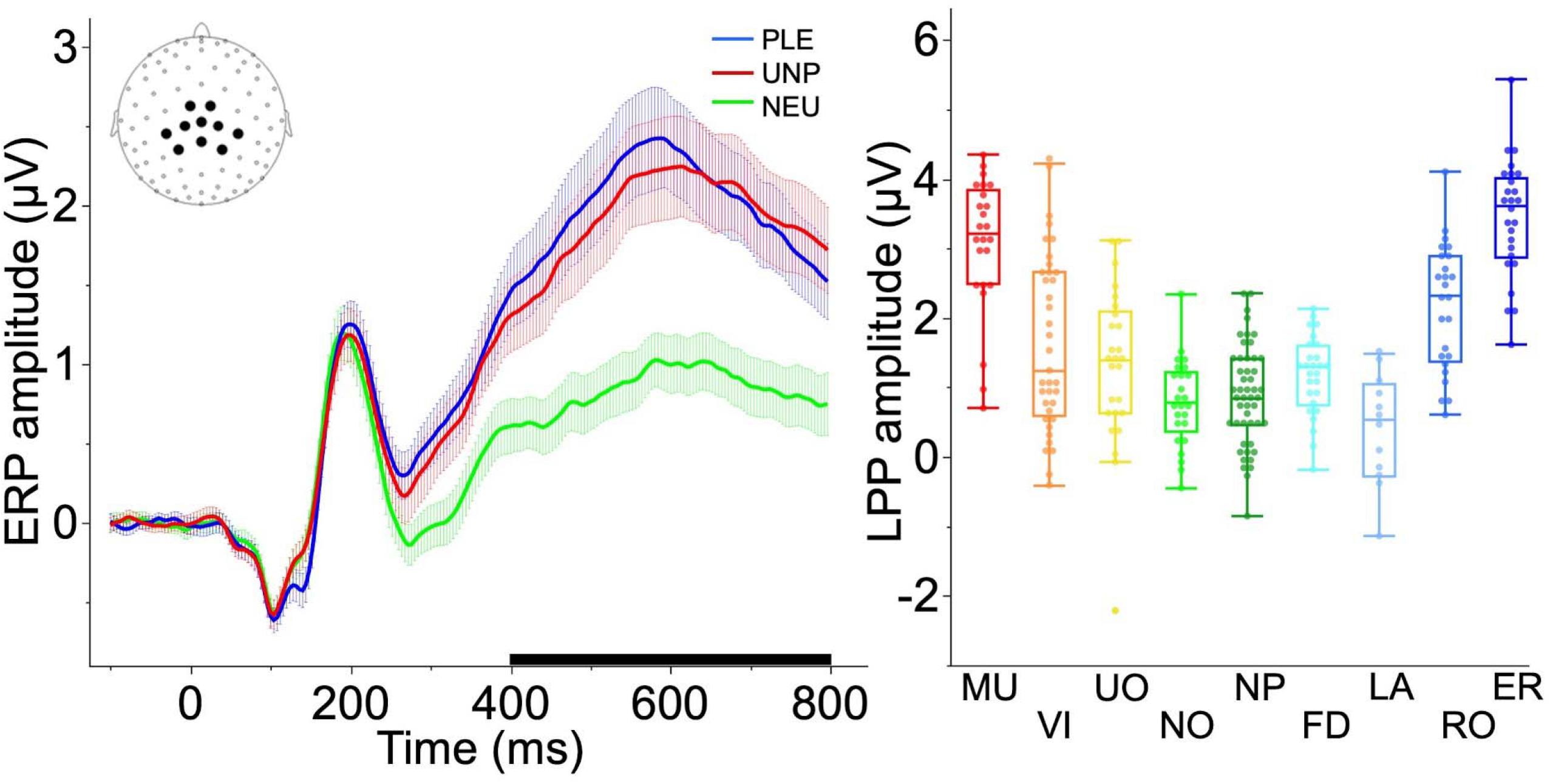
Left. Grand-averaged ERP waveforms (the shaded area represents a 95% confidence interval) for each image across participants in the central sensors (illustrated in the inset) show that, on average, both pleasant and unpleasant contents increase the amplitude of the late positive potential (LPP) compared to neutral contents; The black box on the X axis indicates the time interval (400-800ms) used to compute the Late Positive Potential (LPP). Right. LPP amplitude elicited by pictures of different contents. Note. MU: Mutilations; VI: Violence; UO: Unpleasant Objects; NO: Neutral Objects; NP: Neutral People; FD: Food; LA: Landscapes; RO: Romance; ER: Erotica.

#### 3.2.2 LPP amplitude

In these analyses, first we assessed differences in LPP amplitude between emotional categories across the whole sample and then separately for males and females. The amplitude of the LPP (400-800 ms post-stimulus onset) was positively correlated with arousal ratings, r = 0.61, p<.001. As expected, LPP amplitude changed as a function of emotional content (main effect of Category: F_8,242_ = 32.48, p<001) and, as illustrated in Figure 2 (right), larger LPP amplitudes were observed for erotic pictures and mutilations, followed by romance and violence, and then food, landscapes, unpleasant objects, neutral people, and neutral objects (pairwise comparisons are reported in Table 2). An identical LPP pattern was observed for both males and females (Figure 3, left), where the largest LPP amplitude was observed for erotic pictures and mutilations. The LPP amplitude for erotic pictures and mutilations was also significantly larger than the LPP amplitude for all other stimulus contents (pairwise comparisons are reported in Table S1) and did not differ from one another. A small difference between males and females was found for LPP amplitude evoked by erotic pictures (F_1,50_ = 4.12, p = .048), where males showed slightly larger LPP amplitudes than females.

**Figure 3.**
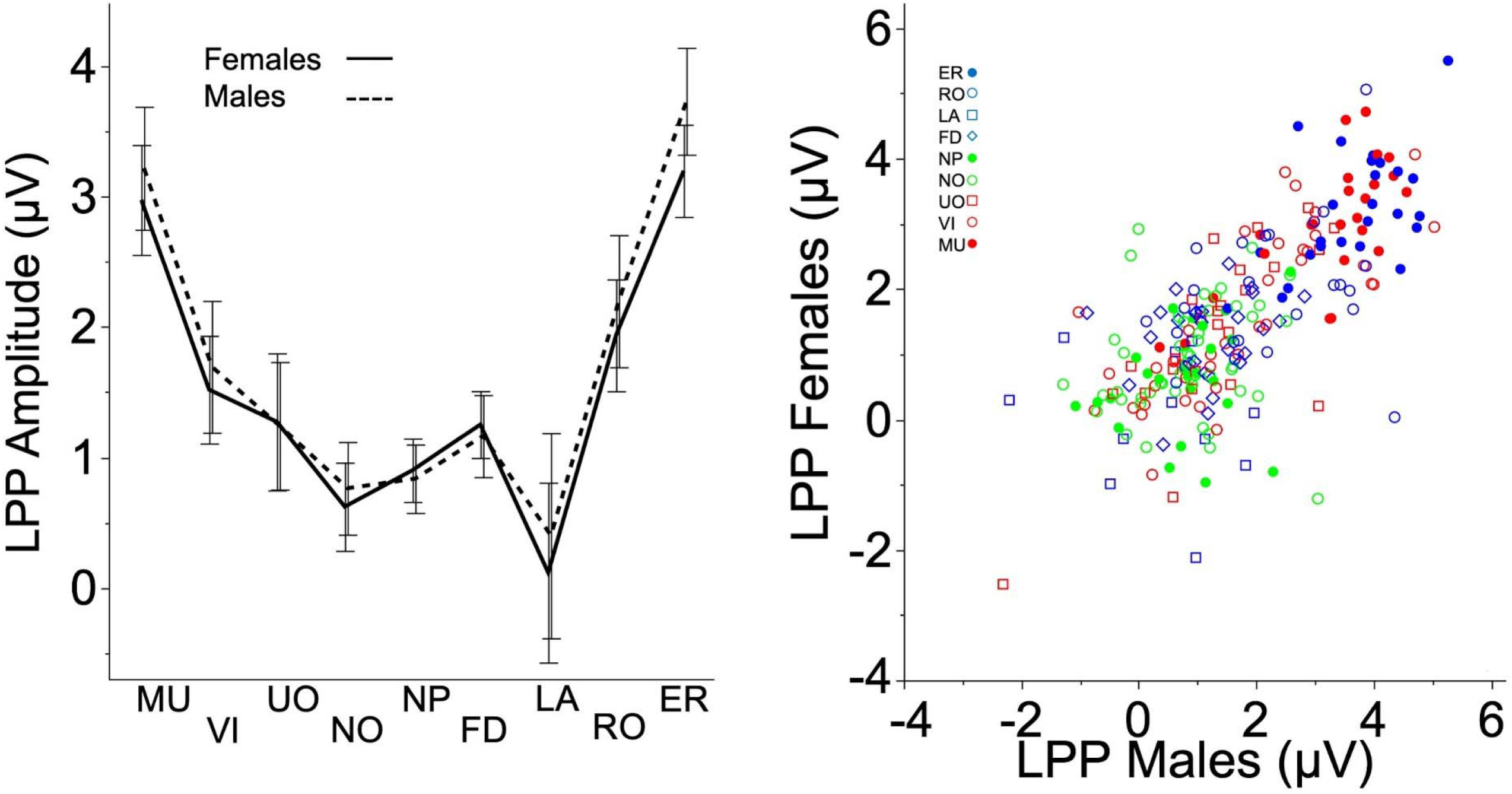
Mutilations and erotica prompt the largest LPPs (400–800 ms) compared with other contents across the entire sample (left) and in both males and females (right). The bars represent 95% CI. Note. MU: Mutilations; VI: Violence; UO: Unpleasant Objects; NO: Neutral Objects; NP: Neutral People; FD: Food; LA: Landscapes; RO: Romance; ER: Erotica.

We observed a moderate correlation between the LPP amplitude elicited by each picture from the HTX sample for males and females (R^2^ = .52; Figure 3 right), suggesting that the contribution of each exemplar into the average categorical effect reported earlier differed between sex. Hence, LPP values are separately reported in the supplementary material for males and females.

### 3.3 Cigarette-related pictures

*These analyses aimed at exploring the extent to which self-reported ratings of valence and arousal, and electrophysiological responses to the images differed between smokers and non-smokers.* Both smokers and non-smokers showed the classic ‘boomerang’ shape for the affective space (Figure 4, left) with a clear distinction for cigarette pictures (black) that stood out compared to other emotional and neutral contents (faded colors). A main effect of content was observed in both smokers and non-smokers for both valence (smokers: F_9,172_ = 253.88, p<.001; non-smokers: F_9,172_ = 347.91, p<.001) and arousal (smokers: F_9,172_ = 59.86, p<.001; non-smokers: F_9,172_ = 83.45, p<.001)), indicating that subjective emotional experience varied based on pictures contents of the pictures. Non-smokers rated cigarette pictures as more unpleasant than neutral contents and lowest in arousal compared to every other category, while smokers rated cigarette pictures similarly to neutral scenes in terms of valence but reported arousal levels similar to those reported when viewing romance and unpleasant objects (pairwise comparisons for smokers and non-smokers are reported in Table S3). Importantly, a direct comparison between smokers and non-smokers revealed that smokers rated cigarette-related pictures as more pleasant (p<.001) and more arousing (p<.001) than non-smokers (Figure 4, right).

**Figure 4.**
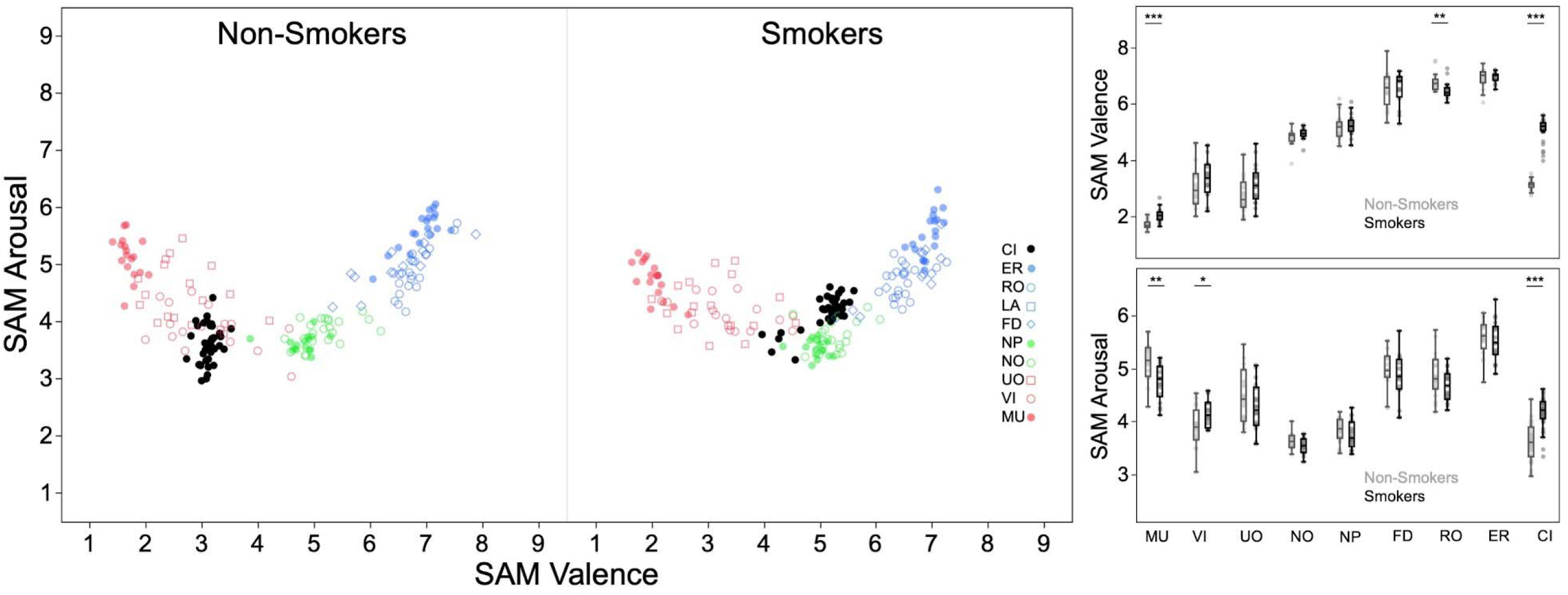
On the left, the affective space for smokers and non-smokers illustrates the relationship between valence and arousal for emotional (faded colors) and cigarette pictures (black). On the right, valence (top) and arousal (bottom) ratings across contents for smokers (light gray) and non-smokers (black). The bars represent 95% CI. Note. MU: Mutilations; VI: Violence; UO: Unpleasant Objects; NO: Neutral Objects; NP: Neutral People; FD: Food; RO: Romance; ER: Erotica; CI: Cigarettes.

Stimulus content modulated the LPP amplitude similarly for both non-smokers (F_8,173_ = 26.95, p<.001) and smokers (F_8,173_ = 26.20, p<.001): pleasant, unpleasant, and cigarette pictures prompted larger LPPs than neutral ones (Figure 5, left). For both groups erotica and mutilations prompted the largest LPPs and neutral contents the lowest (Figure 5, right; pairwise comparisons between categories, separately for non-smokers and smokers, are reported in Table S3). We did not observe any between-group differences in LPP amplitude for any category, including cigarette-related pictures (p = .088; all other categories: p_s_ > .31).

**Figure 5.**
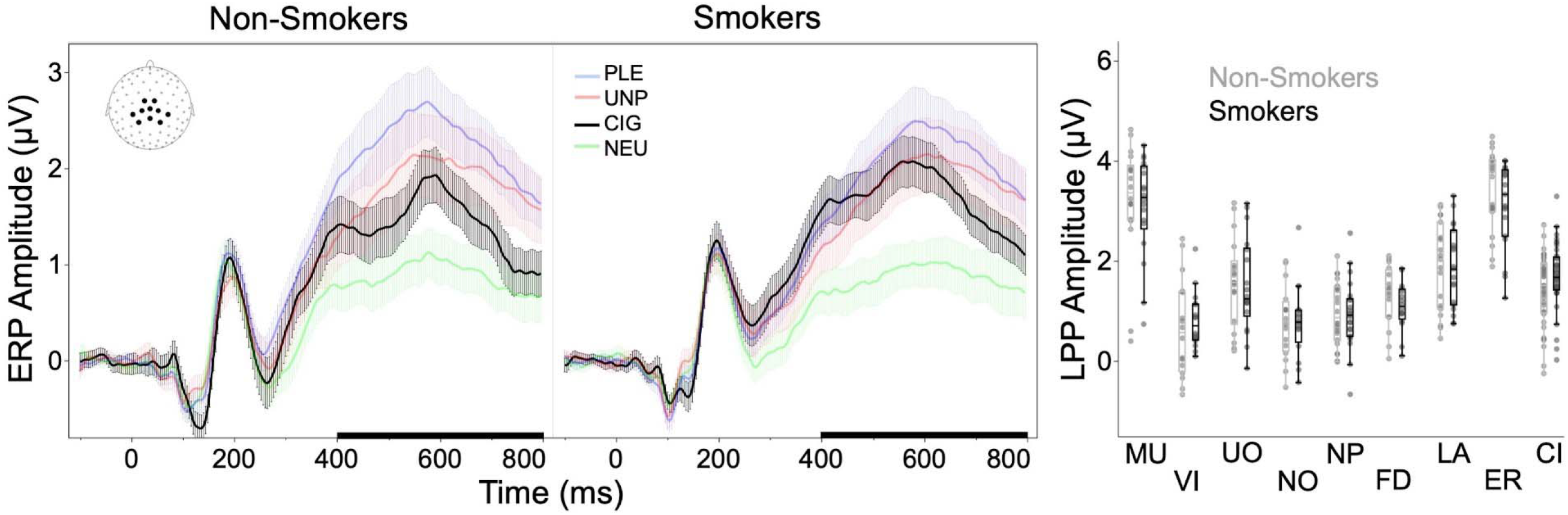
Left. Grand-averaged ERP waveforms (he shaded area represents a 95% confidence interval) for each image across participants in the central sensors (illustrated in the inset) show that, on average, pleasant (PLE), unpleasant (UNP), and cigarette-related contents (CIG) increase the amplitude of the late positive potential (LPP) compared to neutral contents (NEU); The black box on the X axis indicates the time interval (400-800ms) used to compute the Late Positive Potential (LPP). Right. LPP amplitude elicited by pictures of different contents for non-smokers and smokers. Note. MU: Mutilations; VI: Violence; UO: Unpleasant Objects; NO: Neutral Objects; NP: Neutral People; FD: Food; RO: Romance; ER: Erotica; CI: Cigarette.

## 4. DISCUSSION

The primary aim of the current picture-based analysis was to report electrophysiological normative responses for naturalistic images across a wide variety of pleasant, neutral, unpleasant, and cigarette-related contents. By averaging LPP responses (a robust measure of motivational relevance) across subjects, instead of across images, we were able to compute the mean LPP response associated with each image. These values are available in the supplementary material and could be used to integrate information derived from self-reports of valence and arousal to improve the selection of experimental materials to probe both the appetitive and defensive motivational systems.

### 4.1 Replicability of self-reports of valence and emotional arousal

In a first set of analyses, the normative data of valence and arousal from the IAPS picture set were used as a ‘benchmark’ for the HTX dataset. Valence and arousal ratings from the HTX sample scattered across the affective space in a pattern similar to that of the IAPS normative ratings: neutral stimuli grouped at the center of the ‘boomerang’ shape and arousal progressively increased as ratings of pleasantness or unpleasantness increased.

While valence ratings in the HTX sample were similar to the IAPS normative ratings, the arousal ratings from the HTX sample covered a more restricted range of values. Demographic differences across participants may have contributed to this inconsistency. While we enrolled a diverse community sample ranging in age from 18 to 65 years, the IAPS norms were obtained from undergraduate students, the standard sample of most research studies on emotion (Henrich et al., 2010). Anecdotally, however, we noticed that participants were more confused by the arousal scale than by the valence scale and most questions directed to the research assistants during the practice trials were about the interpretation of the arousal scale. We think that confusion about how to interpret the arousal scale might have led some participants to give implausible ratings to some images (See Supplementary Discussion for details).

Despite the restricted ratings range for arousal, the overall pattern of valence and arousal ratings in the HTX sample was remarkably similar to the pattern found in the IAPS normative rating and replicates previous findings (Bradley et al., 2001a). In fact, valence ratings in the HTX analysis demonstrated that all pleasant contents were rated as more pleasant than neutral contents. Unpleasant objects and pictures of violence were rated with similar valence ratings, followed by mutilation pictures that were rated as the least pleasant of the dataset. Similar levels of pleasantness were reported when viewing erotica, romance, pleasant objects, and food. On average, pictures depicting erotic contents were rated as the most arousing, while all other emotional contents (romance, pleasant objects, food, unpleasant objects, violence, mutilations) were rated similarly in terms of induced arousal; neutral pictures (people and objects) were rated lowest in arousal. Males and females reported very similar ratings of valence and arousal across all categories. Replicating previous studies (Bradley et al., 2001b), males rated erotic scenes as more pleasant and more arousing than females, while females rated mutilations as more unpleasant and more arousing than males. Another important replication is that when smokers and non-smokers were compared directly, smokers rated cigarette-related pictures as more pleasant and more arousing than smokers (Deweese et al., 2018). Closely replicating established patterns of self-reported ratings of valence and emotional arousal validated our experimental approach and allowed us to proceed with our analyses of the LPP.

### 4.2 Late Positive Potential (LPP)

Across the entire sample, the centroparietal LPP responses evoked by pleasant and unpleasant images were higher than those evoked by neutral images. Specifically, the highest LPPs were elicited by pictures of erotica and mutilations, followed by romance, food, unpleasant objects, and violence, with the lowest LPP elicited by landscapes, neutral objects, and neutral people. This pattern replicated previous studies demonstrating that the largest LPP amplitudes are reliably evoked by erotica and mutilation pictures (De Cesarei & Codispoti, 2011; Schupp et al., 2000; Weinberg & Hajcak, 2010) and extends previous research showing that these contents prompt the largest skin conductance changes (e.g., Bradley et al., 2001a), the largest attenuation of startle probe P3 amplitude (Bradley et al., 2006), enhanced scanning (Bradley, Costa, & Lang, 2015), the largest pupil dilation (Bradley et al., 2017), largest attentive capture (Padmala et al., 2018), and enhanced blood-oxygen-level-dependent (BOLD) activity (Sabatinelli et al., 2005; Sambuco et al., 2020; Versace et al., 2011). When we analyzed separately the LPP responses from males and females, we observed the same pattern of brain responses we observed across the entire sample: erotica and mutilations prompted the largest LPPs followed by romantic, violence, unpleasant objects, food, neutral people, neutral objects, and landscapes.

Since the data pooled for the HTX analyses came from studies that were not originally designed to provide a normative evaluative and LPP responses, the number of subjects that contributed to each picture in the dataset is not perfectly balanced. Hence, the mean value calculated for pictures viewed by fewer subjects might be somewhat less reliable than the mean value calculated for images viewed by a larger sample. That being said, the picture viewed by the fewest number of subjects was presented to 47 participants, and on average, 160 participants contributed to each picture’s value.

### 4.3 Self-reports and LPP: Different and complementary information

Although the LPP and self-report ratings are both indices of emotional arousal, these measures provided partially divergent information about emotional engagement following picture presentation. First, when considering arousal self-reports, erotica were rated as most arousing and neutral pictures as least arousing, but all other emotional contents were rated similarly. That is, mutilation pictures received similar arousal ratings to pictures of food or pleasant and unpleasant objects. Yet, the LPP responses demonstrated that erotica and mutilations were the most effective contents in prompting affective engagement, while images of food and pleasant or unpleasant objects prompted LPPs more similar to those of neutral contents. Second, self-reported ratings of emotional arousal and LPP responses diverged when we considered responses by sex. Males and females rated erotica and mutilations differently in terms of both valence and arousal but showed comparable LPP responses for both contents. Third, smokers and non-smokers rated smoking-related contents differently on both valence and arousal, yet these contents prompted similar LPP responses in both groups. Altogether, these findings demonstrate that patterns of emotional engagement based on self-reports and LPP are not identical and selecting stimuli using self-reports norms alone to investigate neuroaffective engagement may bias the interpretation of the physiological findings (Franken et al., 2008; Ito et al., 1998; Schupp et al., 2000; Weinberg & Hajcak, 2010).

This proposal is obviously not new, and largely stems from influential work suggesting that evaluative dimensions of affect, such as valence and arousal, are only one possible system to investigate affective reactions. Lang (1988, 2010) proposed that the investigation of emotional responses can be pursued across three separate systems: (1) the language of emotion (expressive and evaluative; SAM ratings of valence and arousal in the present study); (2) physiological changes (somatic, autonomic, and electrophysiological; LPP in the present study); and (3) behavior (e.g., approach and avoidance, “freezing”). Hence, selecting stimuli solely based on the evaluative dimension to investigate physiological correlates of emotional processes might be problematic for several reasons. First, subjective experiences of emotion and biologically derived measures of affect are known to be poorly correlated (Taschereau-Dumouchel et al., 2022). Second, conscious evaluations are fully formed a few seconds after pictures are presented, at the end of a cascade of processes that also include appraisal. Neuroaffective measures such as the LPP, however, reliably peak within a short interval of encountering an emotional stimulus (for the LPP, between 300 to 900 ms post picture onset), providing information regarding the extent of attentional and motivational engagement on a millisecond basis. Finally, expressive and evaluative measures require individuals to willingly share their responses. Neuroaffective measures are particularly useful in this case because they do not require a behavioral response to document processing of visually presented information. Hence, these measures can circumvent the limitations of self-report assessments while still reflecting subtle individual differences in affective stimulus processing.

While advanced methods to select pictures for psychophysiological experiments based on valence and arousal ratings can be developed (Wilson et al., 2021), so far these methods rely exclusively on self-reports. Altogether, the present findings and results from other psychophysiological studies (e.g., Bradley et al., 2017) demonstrated that only some picture contents that are rated as emotionally arousing reliably prompt psychophysiological responses. As recently described by Bradley et al. (2017) “*After more than 25 years of research using emotional pictures, it is now clear that evaluative (arousal) ratings of natural scenes are not synonymous with the degree of engagement of defensive or appetitive activation measured in the brain or body*” (pag. 1432). Hence, we propose that evaluative and physiological information should be integrated to select the most appropriate set of pictures to use in psychophysiological assessments.

Given current trends in personalized approaches to treatment and assessment, the results of this study may also help select materials that are better suited for subject-by-subject analyses (Schupp & Kirmse, 2021, 2022). In a series of recent studies evaluating the potential to use neural correlates of affective stimulus evaluation at the individual level, Schupp and Kirmse (2021, 2022) observed significantly larger LPP amplitudes to high- compared to low-arousing emotional stimuli spanning three behavioral systems: predator fear (threatening animals, non-threatening animals), disease avoidance (mutilated and injured bodies, neutral people), and sexual reproduction (erotic couples, romance). LPP amplitude was reliably larger for sexual reproduction than predator fear in 100% of participants, whereas there was slightly more variability among the unpleasant contents. Furthermore, they observed a limited number of non-significant LPP effects that were limited to a single behavioral domain (predator fear) in a small subset of participants, leading the authors to conclude that these effects may be stimulus-specific. Data from the individual subject level may shed light on the mismatch in LPP and self-reported ratings of emotional arousal we observed in the present study and may push forward the need to rely on electrophysiological normative responses to reliably probe appetitive and defensive motivational systems.

### 4.4 Summary

The results of this study demonstrate the replicability of self-reported ratings of valence and emotional arousal and provide electrophysiological normative responses for naturalistic pictures spanning a wide range of contents. In highlighting a divergence in the self-reported stimulus ratings and LPP data, we suggest that using normative electrophysiological responses may be a better suited approach to stimulus selection compared to utilizing only normative ratings of valence and arousal. This approach circumvents the limitations of self-report assessments that rely on subjective and/or behavioral responses, and capitalizes on an objective, neural indicator of the processing of emotionally arousing stimuli. This approach may also be useful moving forward to tailor stimulus selection to populations whose behavior is sensitive to specific environmental cues.

## Acknowledgements

We thank Ms. Maria Kypriotakis for her help in organizing the database.

## Data availability statement

The normative electrophysiological responses of the Late Positive Potential (LPP) reported in the present manuscript are openly accessible in the supplementary material as csv files (1) across the entire sample, (2) separate for males and females, and (3) separately for smoking status (smokers vs. non-smokers). Valence and arousal ratings for each image are also provided.

## SUPPLEMENTARY MATERIALS

**Figure S1.**
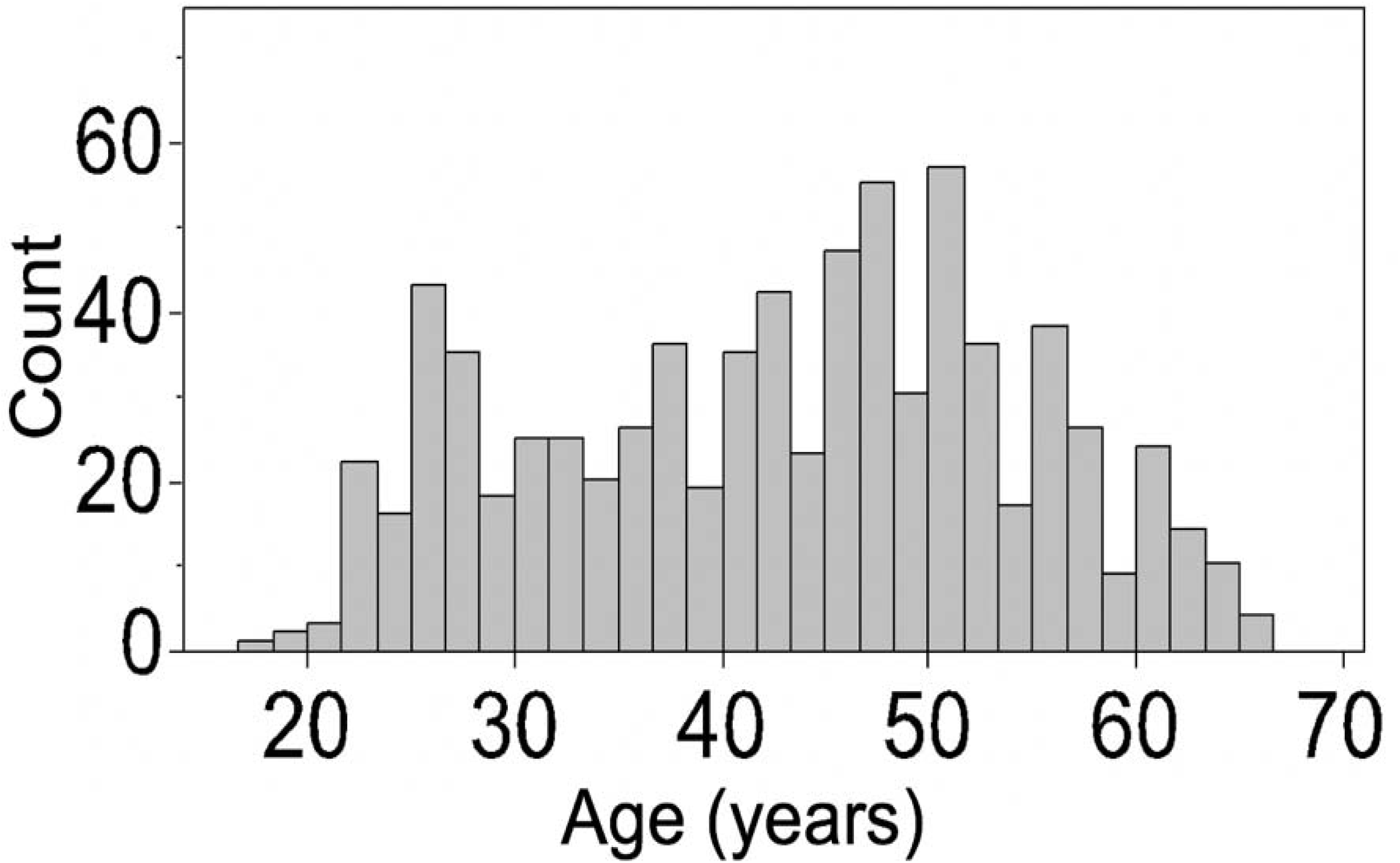
Age distribution (count) of the participants in the three different studies considered in the current analyses.

**Figure S2.**
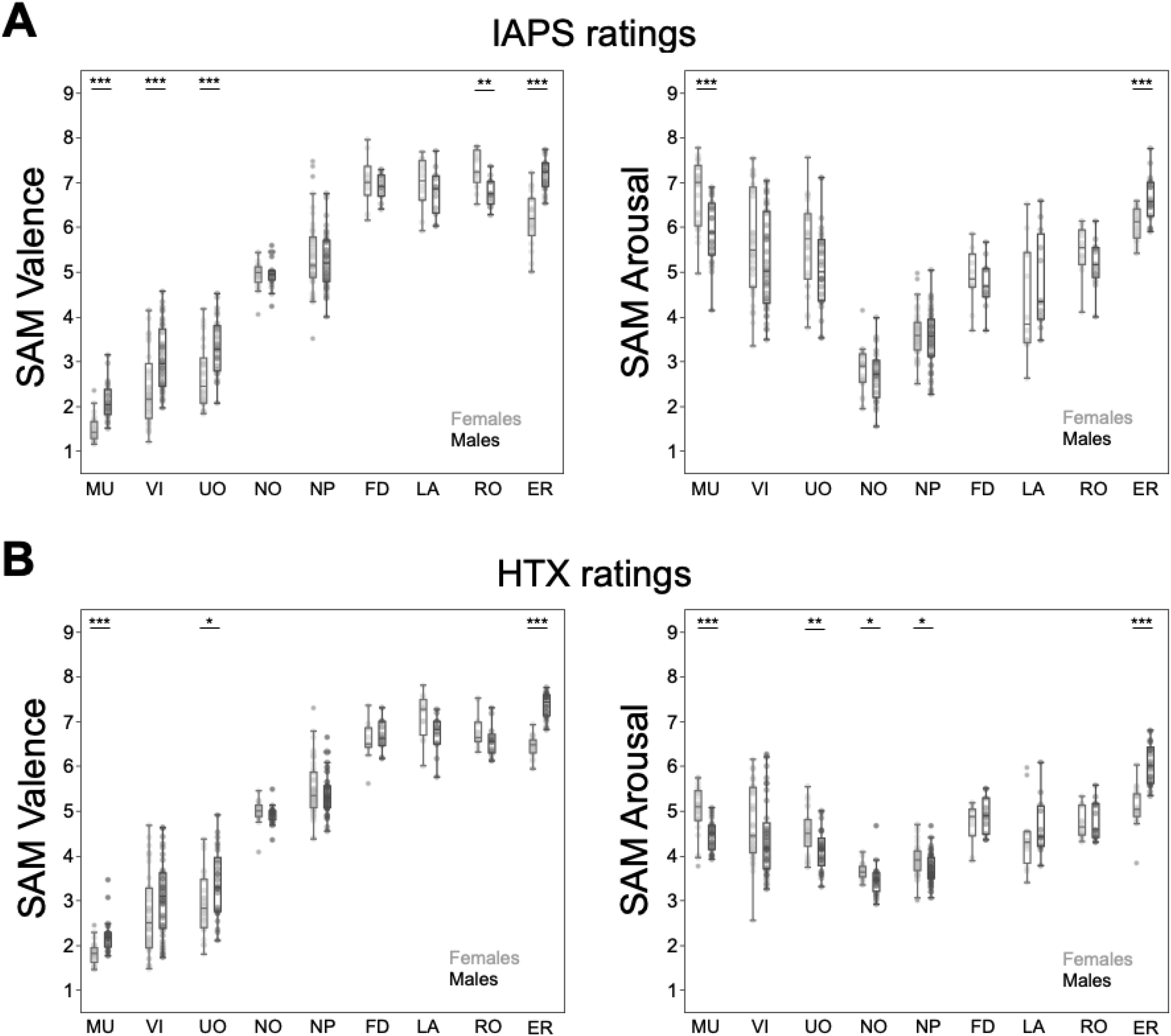
Valence and arousal ratings across the different categories, separately reported for males and females, in both IAPS ratings and HTX ratings. *Note.* MU: Mutilations; VI: Violence; UO: Unpleasant Objects; NO: Neutral Objects; NP: Neutral People; FD: Food; LA: Landscapes; RO: Romance; ER: Erotica. Results of between-group comparison significant at ***p<.001, **p<.01, *p<.05.

**Table S1.** Number of pictures used for each category in the primary analyses reported in the manuscript, assessing patterns of SAM ratings (valence and arousal) and LPP for the whole-sample analysis, the analysis assessing gender differences (males/females), and the analysis on smoking status (smokers/non-smokers).

| Category | Number of Exemplars |  |
| --- | --- | --- |
|  | Whole-sample and males/females | Smokers/Non-Smokers |
| Cigarette | - | 38 |
| Erotica | 26 | 19 |
| Romance | 25 | 19 |
| Landscapes | 12 | - |
| Food | 27 | 16 |
| Neural People | 49 | 22 |
| Neutral Objects | 25 | 16 |
| Unpleasant Objects | 26 | 18 |
| Violence | 37 | 16 |
| Mutilations | 24 | 18 |
| **Total** | **251** | **182** |

**Table S2.**
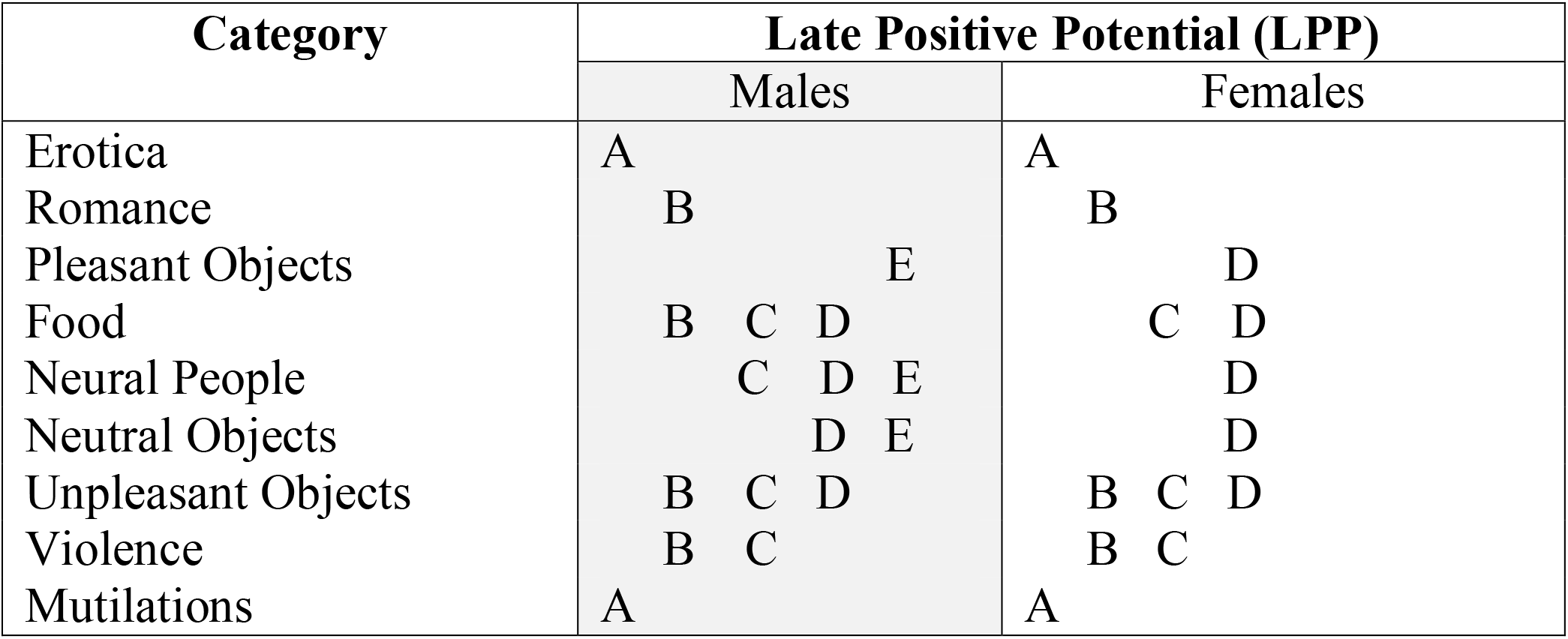
Tukey HSD pairwise comparisons (p<.05) for the LPP, separately reported for males and females. *Note: Categories which share at least one letter do not significantly differ*.

| Category | Late Positive Potential (LPP) |  |
| --- | --- | --- |
|  | Males | Females |
| Erotica | A | A |
| Romance | B | B |
| Pleasant Objects | E | D |
| Food | B C D | C D |
| Neural People | C D E | D |
| Neutral Objects | D E | D |
| Unpleasant Objects | B C D | B C D |
| Violence | B C | B C |
| Mutilations | A | A |

**Table S3.** Tukey HSD pairwise comparisons (p<.05) for SAM rating of valence and arousal separately reported for smokers and non-smokers. *Note: Categories which share at least one letter do not significantly differ*.

| Category | SAM ratings |  |  |  |
| --- | --- | --- | --- | --- |
|  | Valence |  | Arousal |  |
|  | Non-Smokers | Smokers | Non-Smokers | Smokers |
| Cigarette | C | C | D | C |
| Erotica | A | A | A | A |
| Romance | A | B | B | B |
| Food | A | A B | B | B |
| Neural People | B | C | D | D |
| Neutral Objects | B | C | D | D |
| Unpleasant Objects | C D | D | C | C |
| Violence | D | D | D | C |
| Mutilations | E | E | B | B |

**Table S4.** Tukey HSD pairwise comparisons (p<.05) for LPP amplitude in non-smokers and smokers. *Note: Categories which share at least one letter do not significantly differ*.

| Category | LPP |  |  |  |  |
| --- | --- | --- | --- | --- | --- |
|  | Non-Smokers |  |  | Smokers |  |
| Cigarette | B | C |  | B |  |
| Erotica | A |  |  | A |  |
| Romance | B |  |  | B |  |
| Food | B | C | D | B | C |
| Neural People |  | C | D |  | C |
| Neutral Objects |  | C | D |  | C |
| Unpleasant Objects | B | C |  | B | C |
| Violence |  |  | D |  | C |
| Mutilations | A |  |  | A |  |

## Supplementary Discussion

As mentioned in the primary text, we observed a compressed range of arousal ratings in the HTX sample compared to the IAPS normative ratings (see Figure 1C): While mean IAPS ratings of arousal ranged from 2 to 7, in the HTX sample they ranged from 3 to 6. Many participants in our studies were significantly older than the participants commonly enrolled in psychology experiments (including the experiments in which IAPS pictures were originally rated). Hence, it is possible that older individuals rate high arousing images as less arousing than what younger individuals do. It is also possible that an increase in daily consumption of digital media may have dampened the raters’ perceived “shock value” of the most arousing images. With individuals spending upwards of 6 hours per day consuming digital media and nearly doubling time spent online in the past 15 years, it is conceivable that more frequent exposure to highly arousing content may reduce present day self-reported ratings of emotional arousal. In the laboratory, Codispoti et al. (2006) showed that arousal ratings decreased following repeated picture presentation, but that the most arousing picture contents (erotica, mutilations) remained more arousing than neutral. Yet, while these factors might explain the lower arousal ratings attributed to erotica and mutilations, they do not explain the higher arousal ratings attributed to neutral images.

In fact, we are inclined to attribute the differences in self-reported arousal observed between our sample and the IAPS to difficulties in interpreting the meaning of the arousal scale. In the laboratory, we noticed that participants were more confused by the arousal scale than by the valence scale and that most questions directed to the research assistants during the practice trials were about the arousal scale. Visual inspection of the individual responses to the arousal scale highlighted the presence of two peculiar response patterns that were shared by several participants. It is important to consider the layout of the electronic version of the SAM: valence and arousal scales are presented sequentially, with the rating scales moving from left (most pleasant, most arousing) to right (most unpleasant, least arousing). In the first response pattern that we observed, some individuals selected on the Likert scales the same numerical value for both valence and arousal. In the second response pattern, individuals used the full range of values on the valence scale, but consistently reported the same midline value on the arousal scale for all the pictures. The first response pattern, then, reduces the arousal ratings of the most unpleasant images and inflates that of the neutral ones, whereas the second response pattern reduces the arousal rating of both pleasant and unpleasant images and inflates the arousal ratings of neutral images. When averaged with the other ratings to obtain the mean arousal values of individual pictures, these peculiar response patterns reduce the range of arousal ratings (see Figure 1C). Even though we cannot exclude that some individuals might find very unpleasant images also not arousing and others might find neutral images mildly arousing, the frequency of these two response patterns led us to conclude that some individuals had difficulties understanding the meaning of the arousal scale and/or decided not to comply with the instructions.

Despite the potential confusion in a subset of the sample, the LPP results (which are not under volitional control), are not “compressed” (see Figure 2). This provides further justification of the need to integrate an objective neuroaffective measure of emotional engagement with the self-report data when selecting stimuli in experiments designed to engage appetitive and defensive motivational systems.

### Individual Ratings

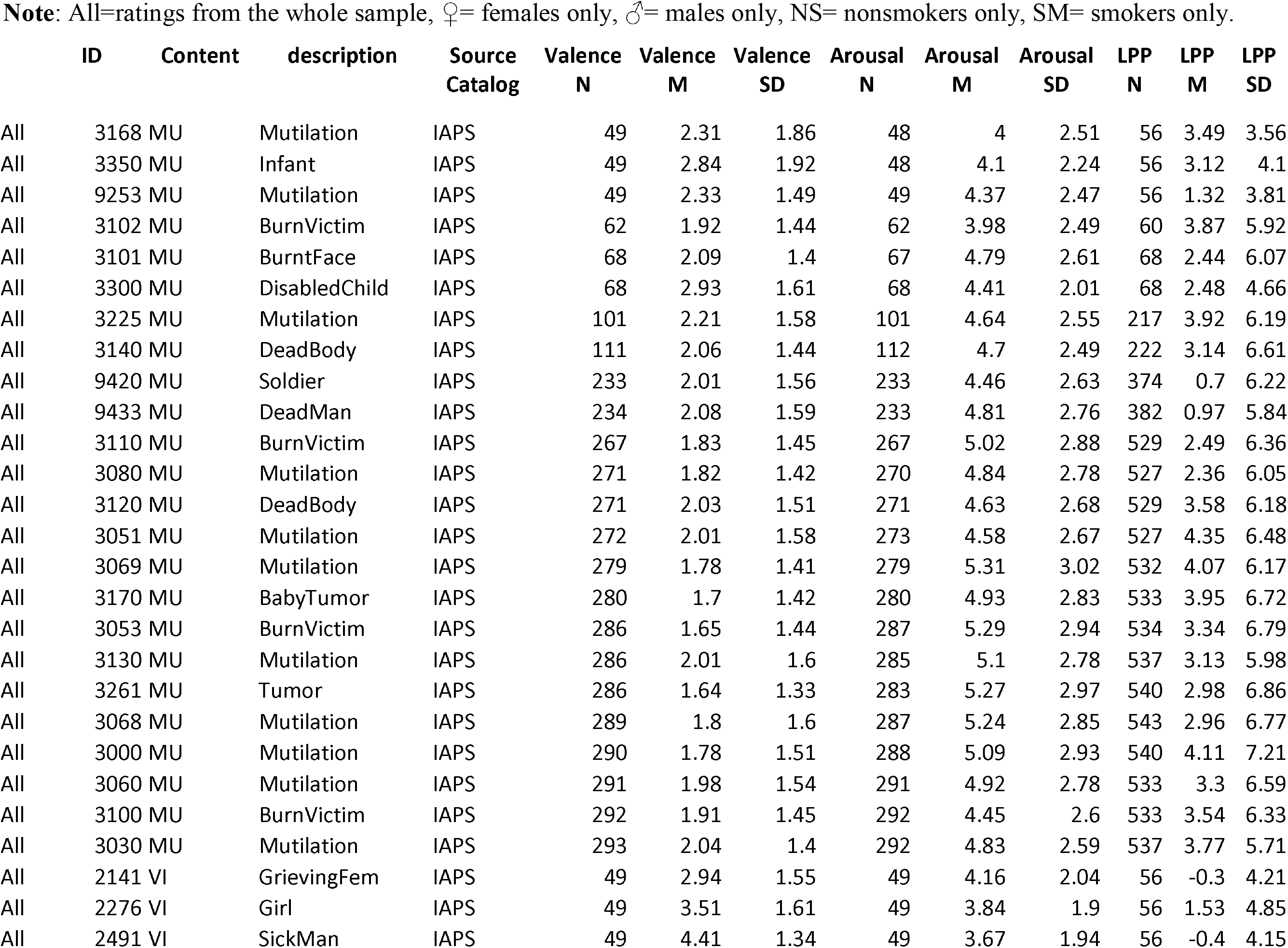
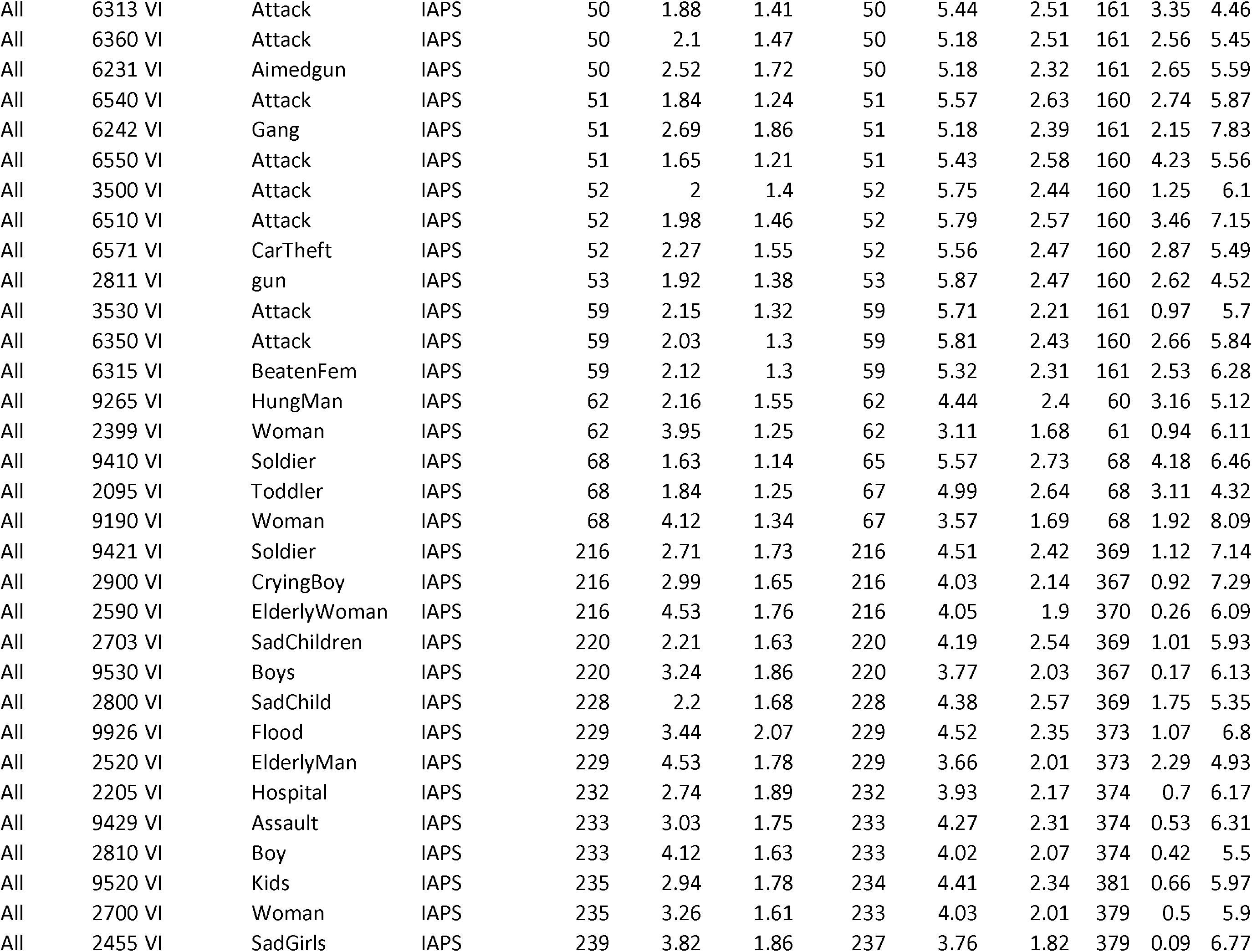
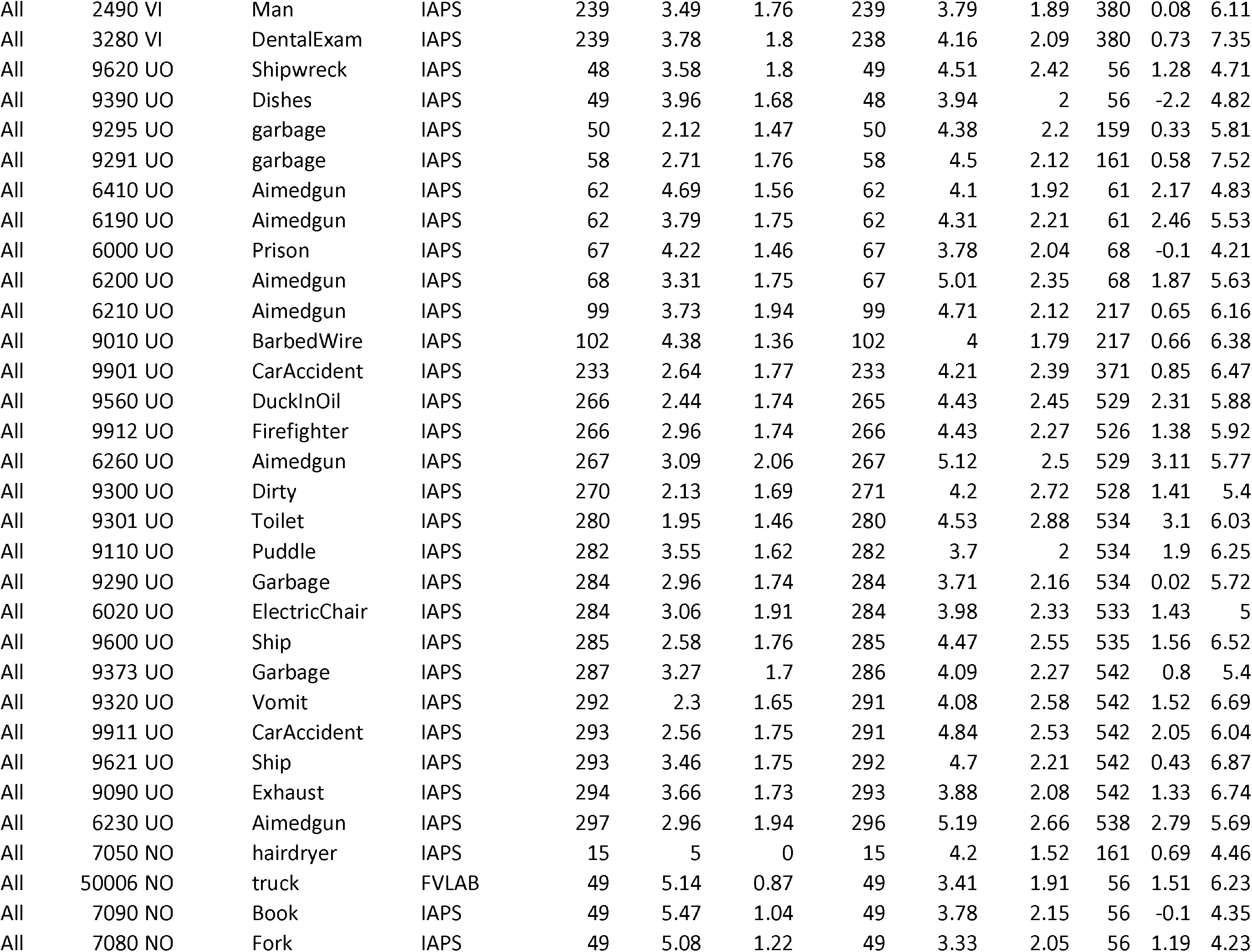
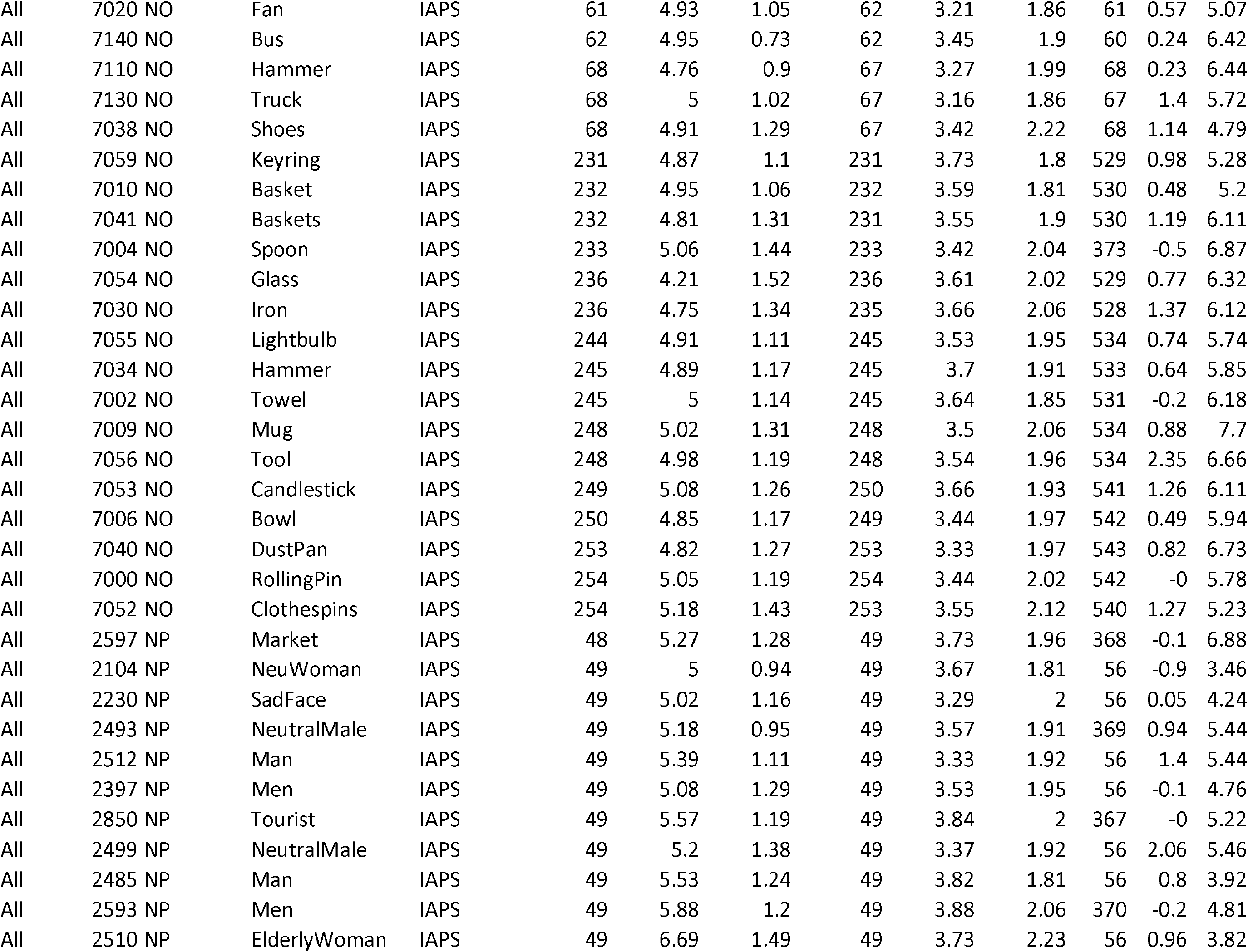
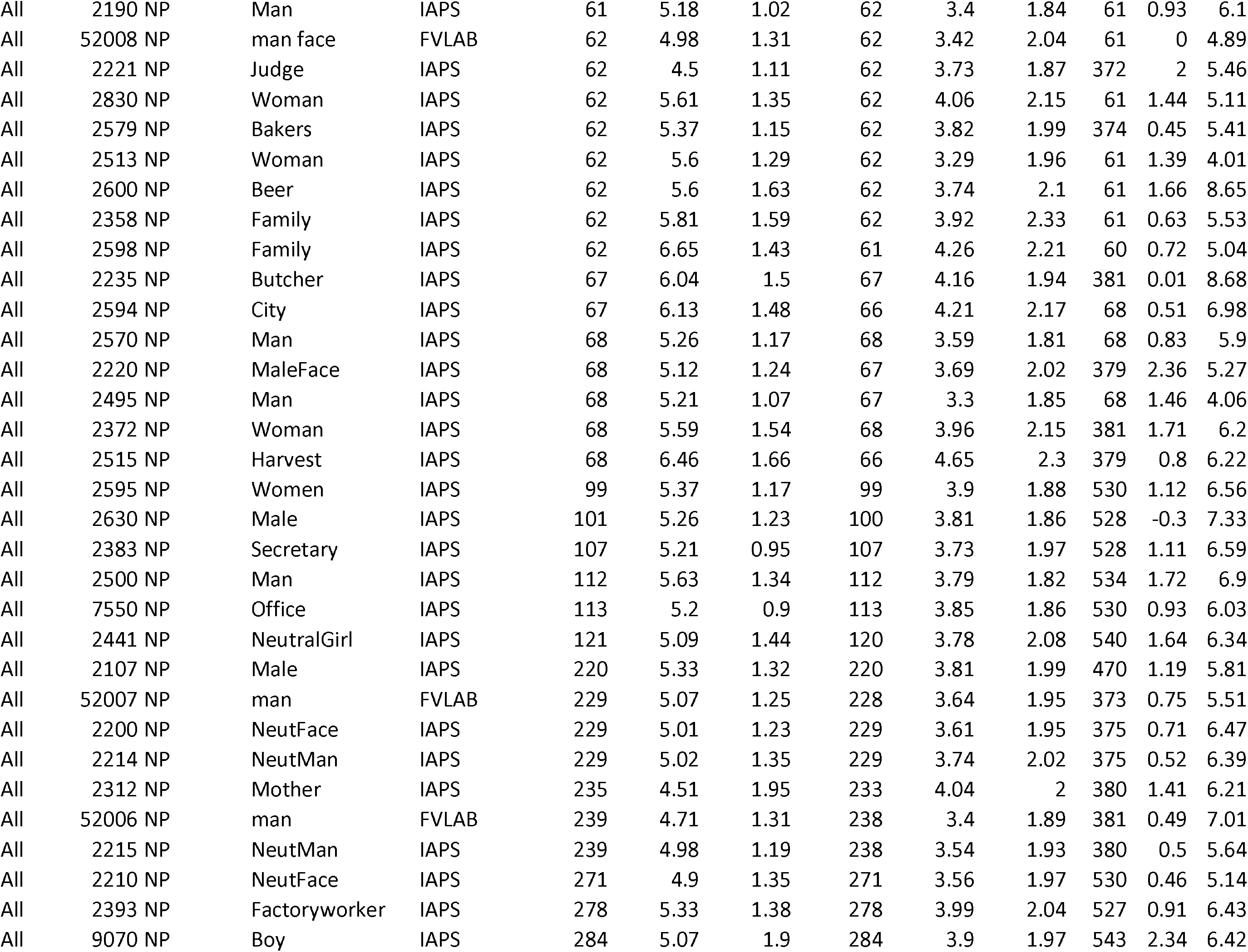
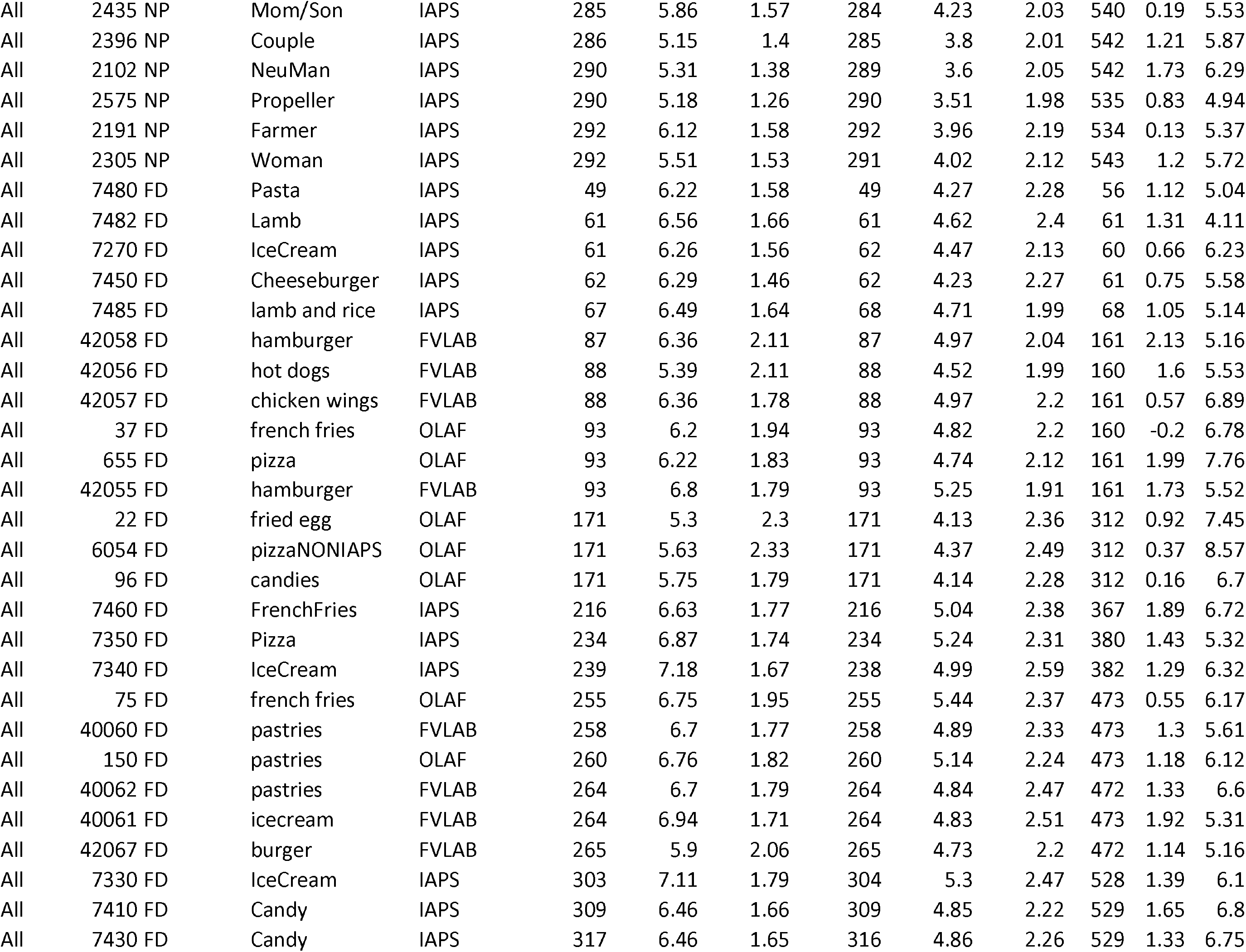
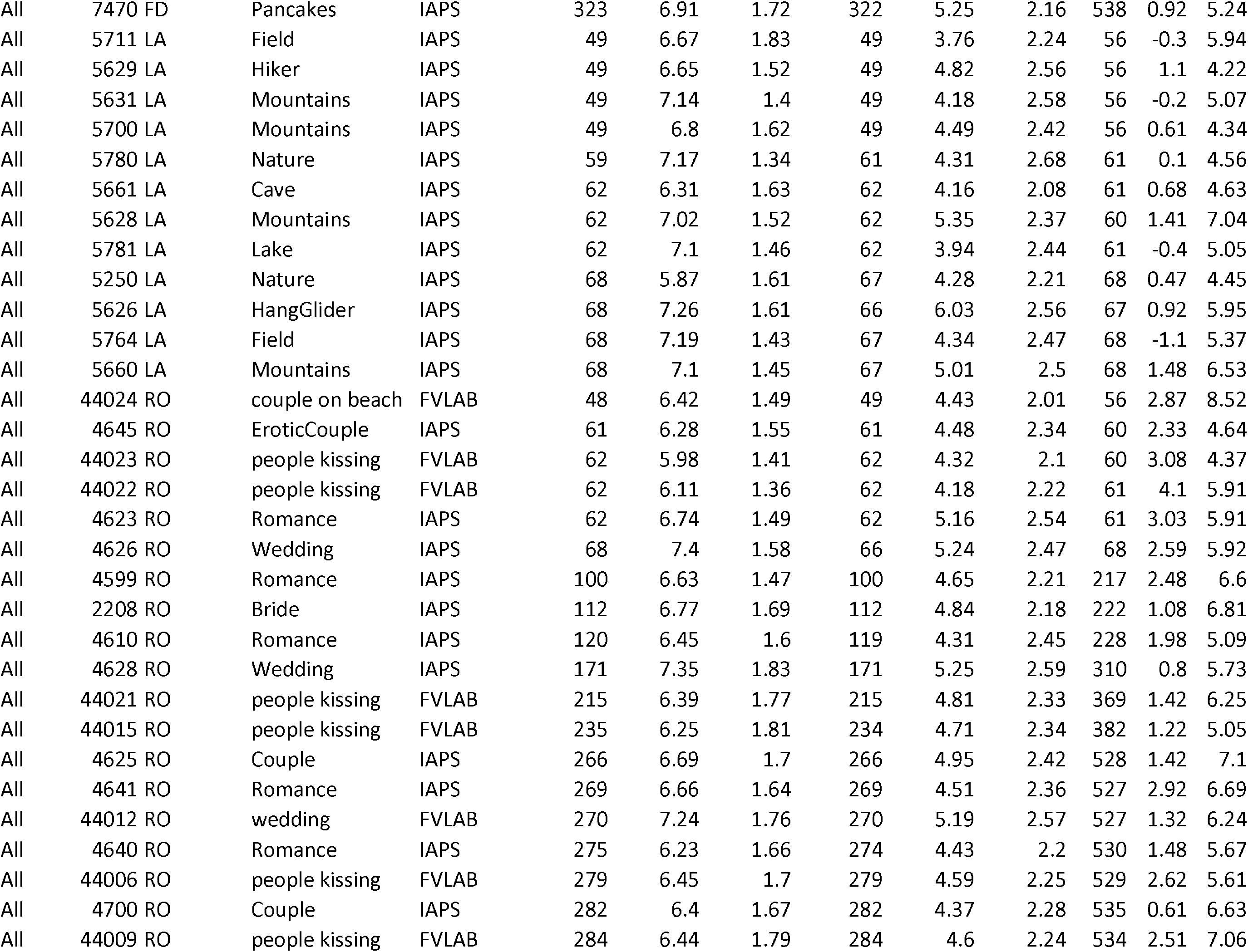
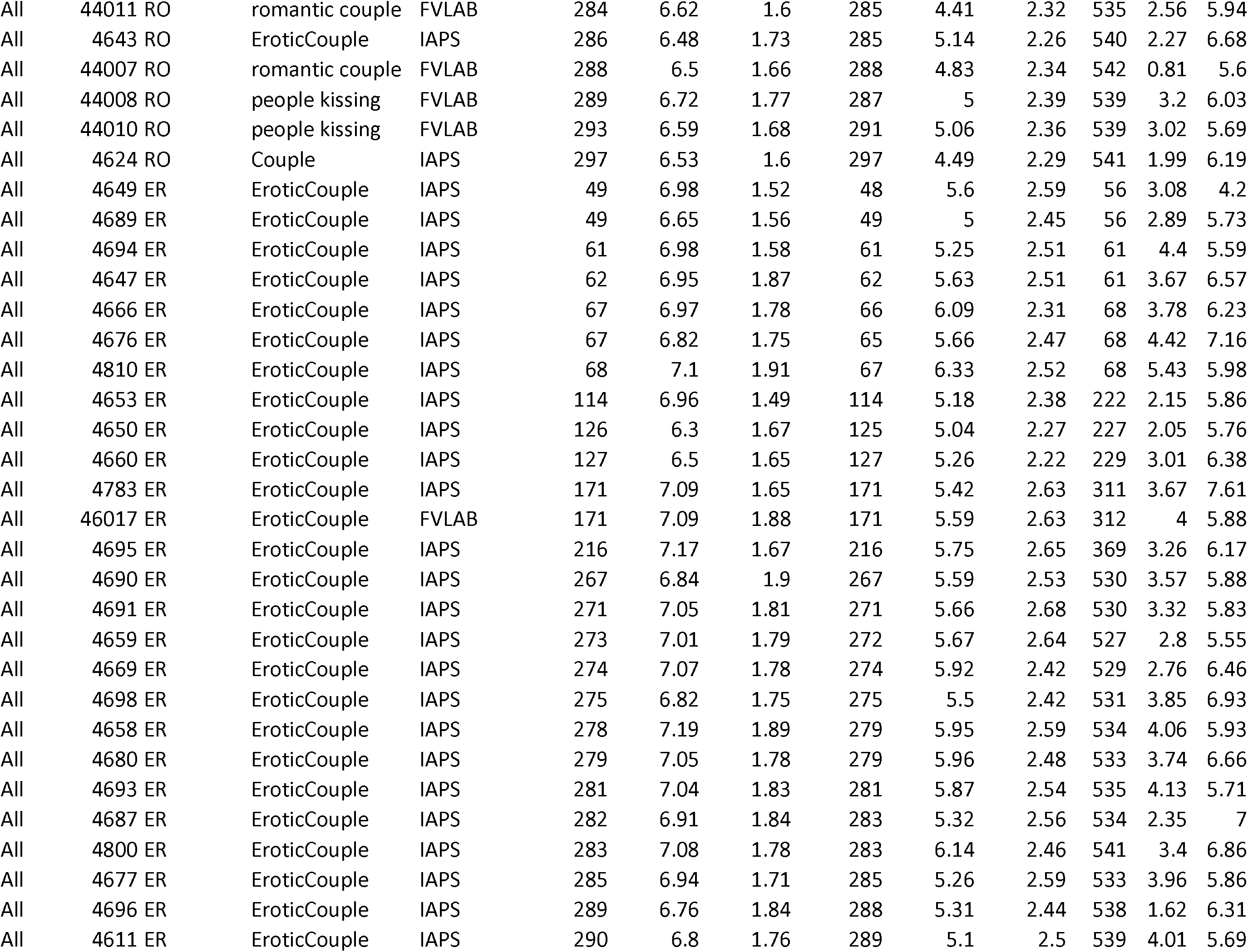
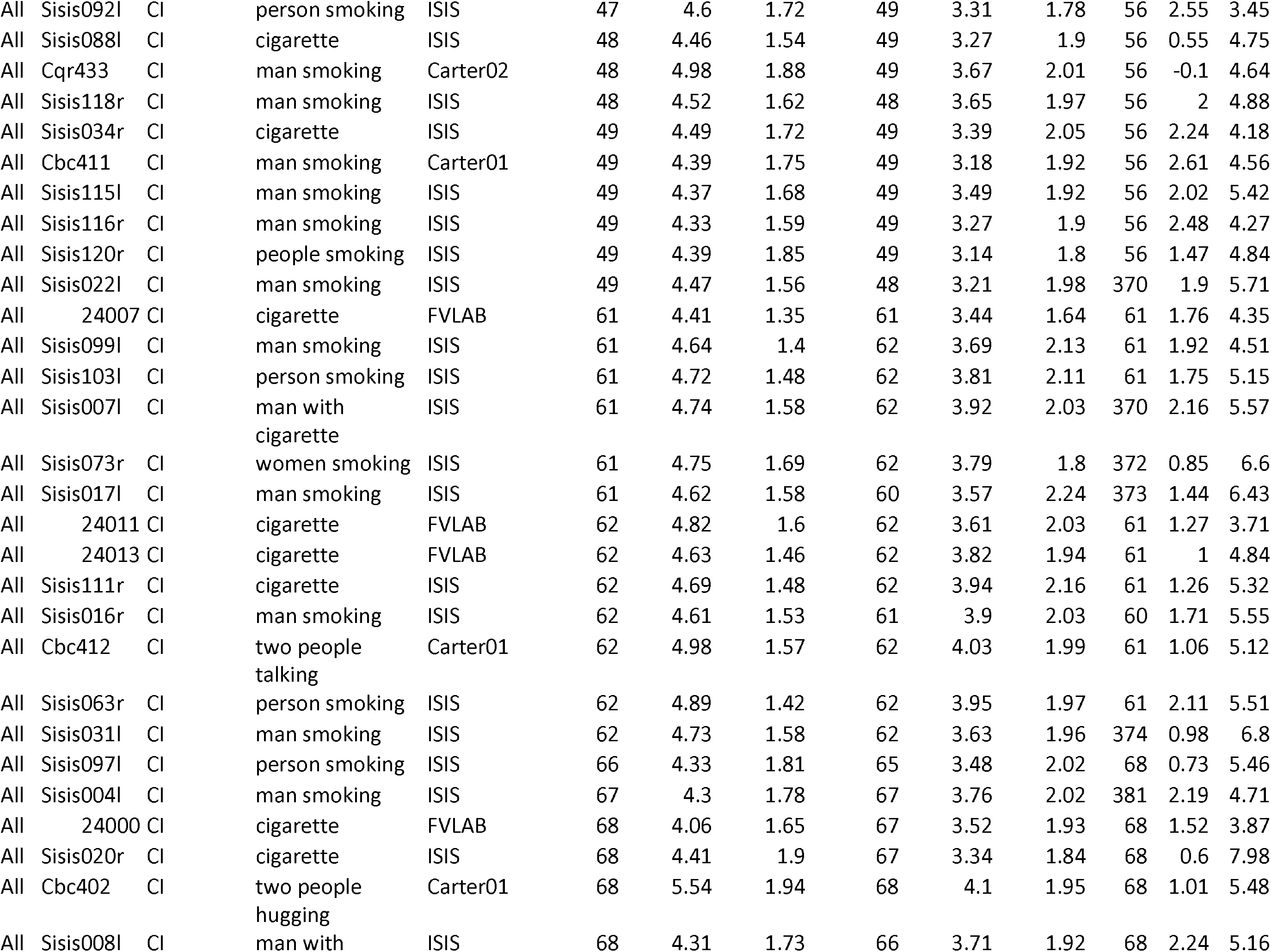
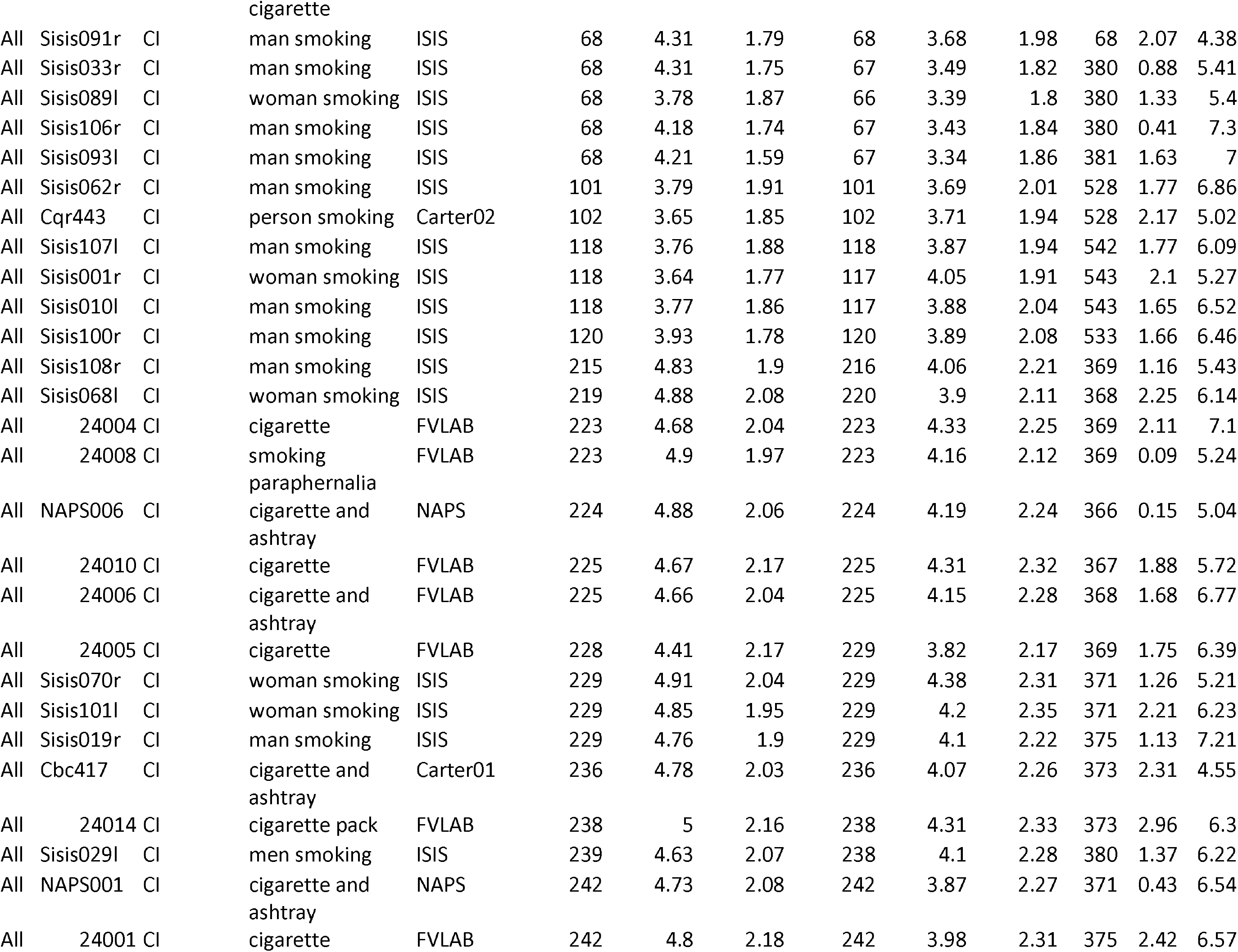
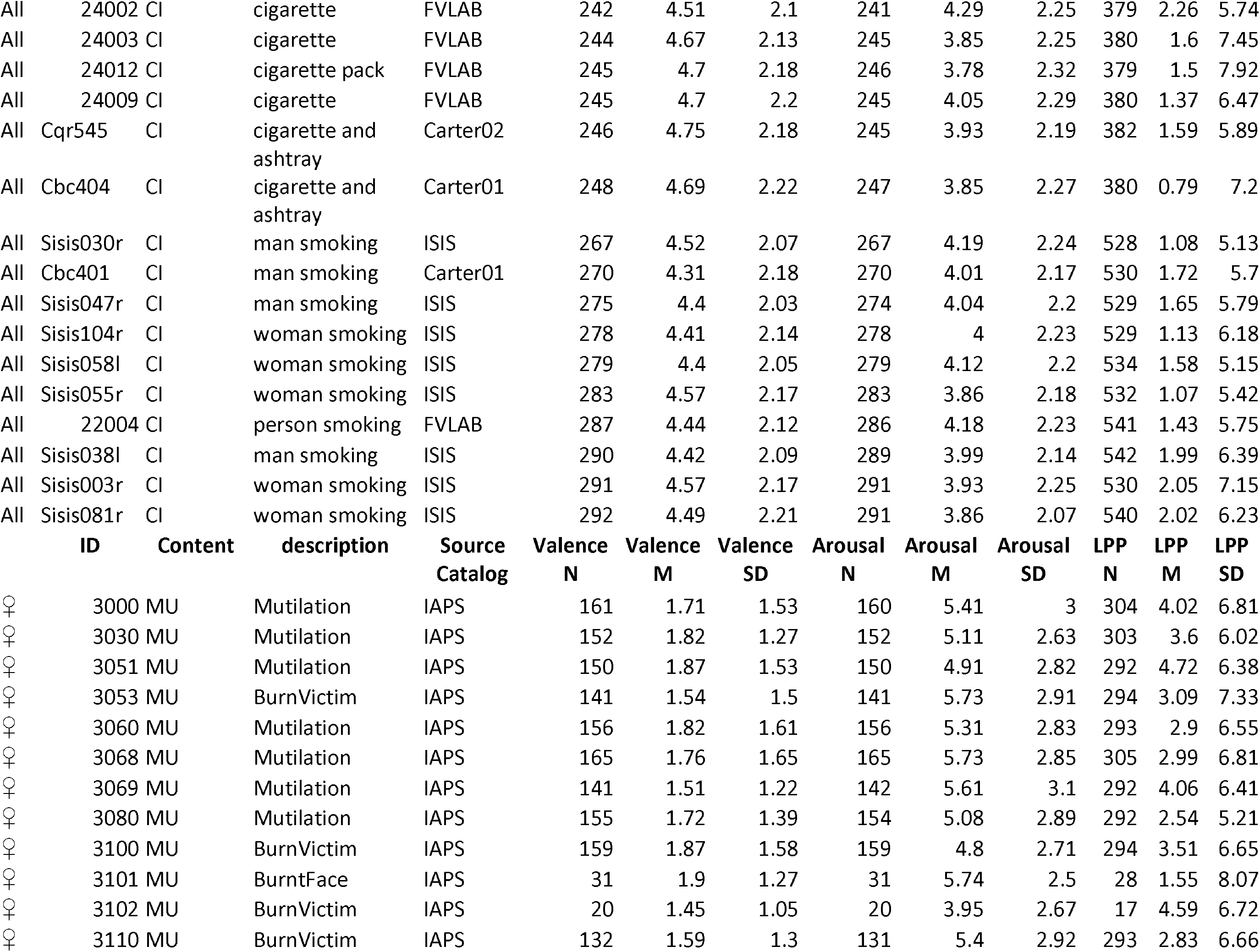
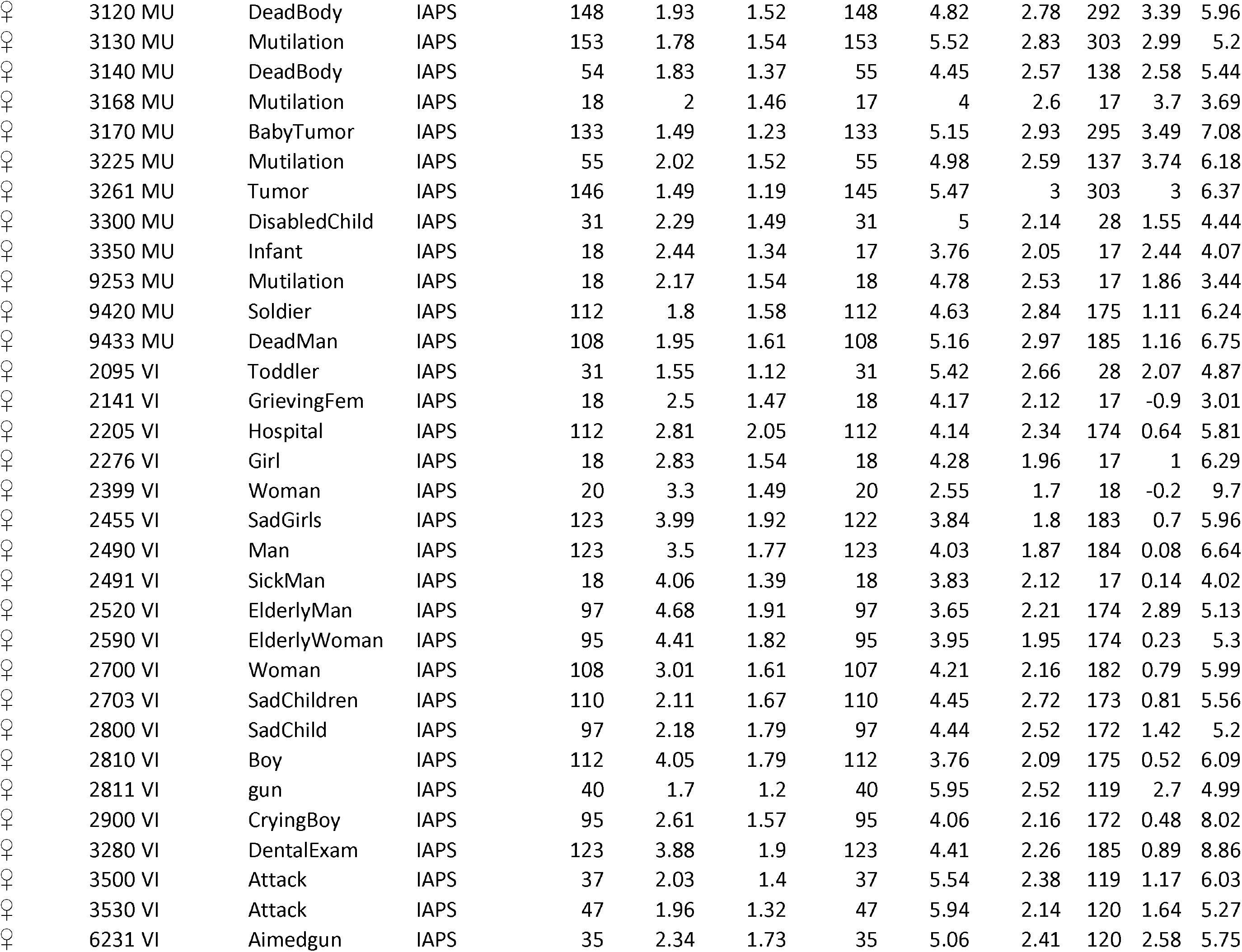
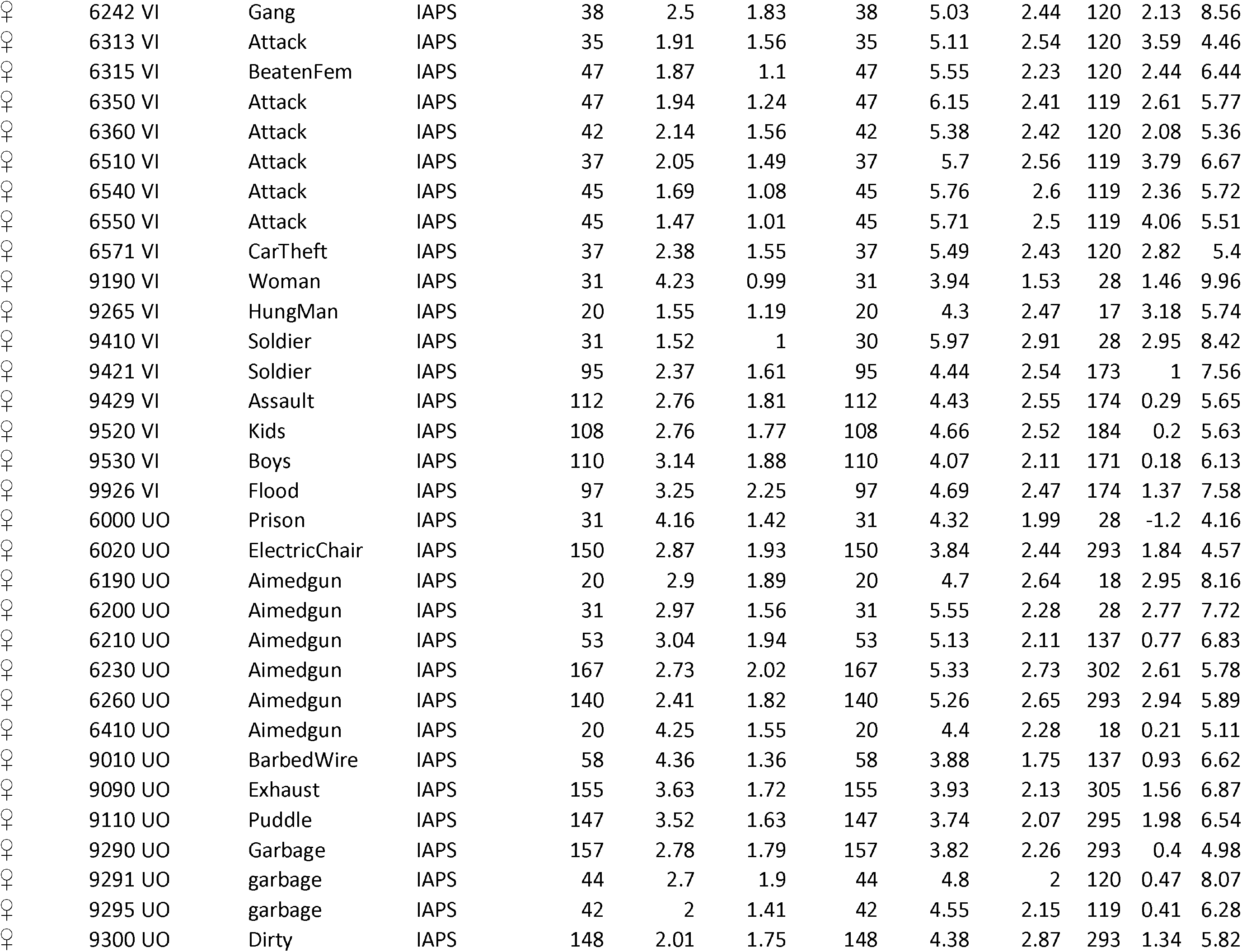
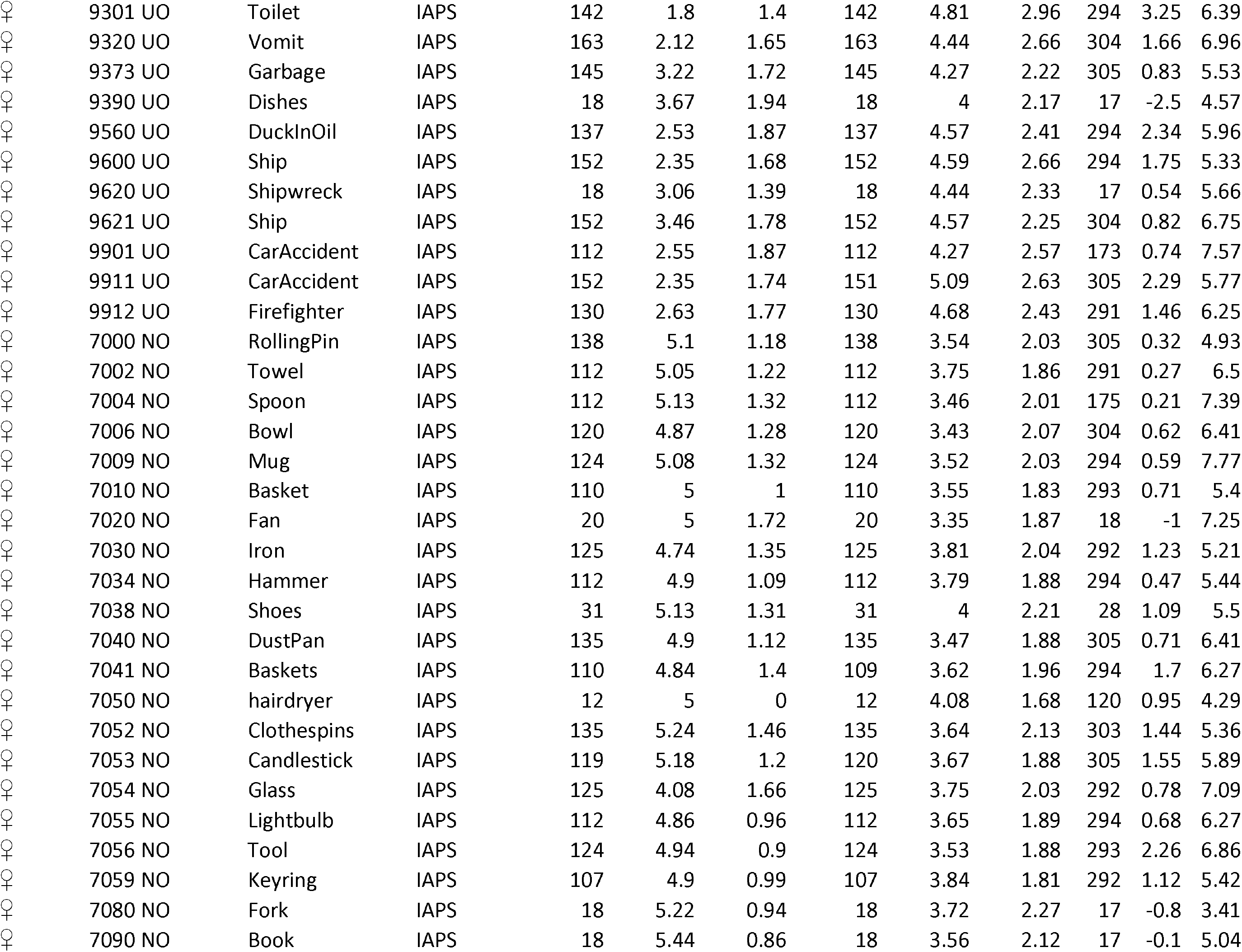
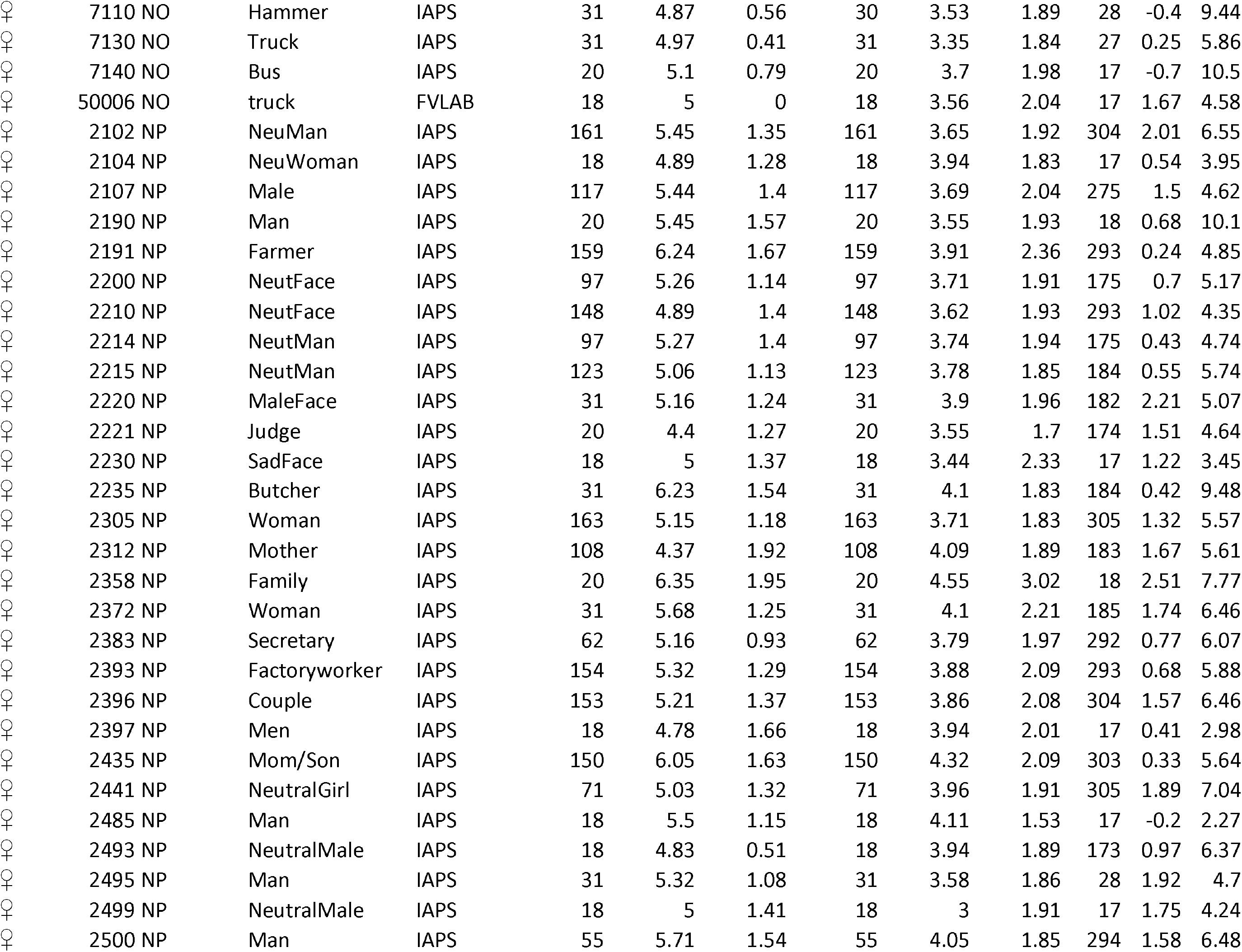
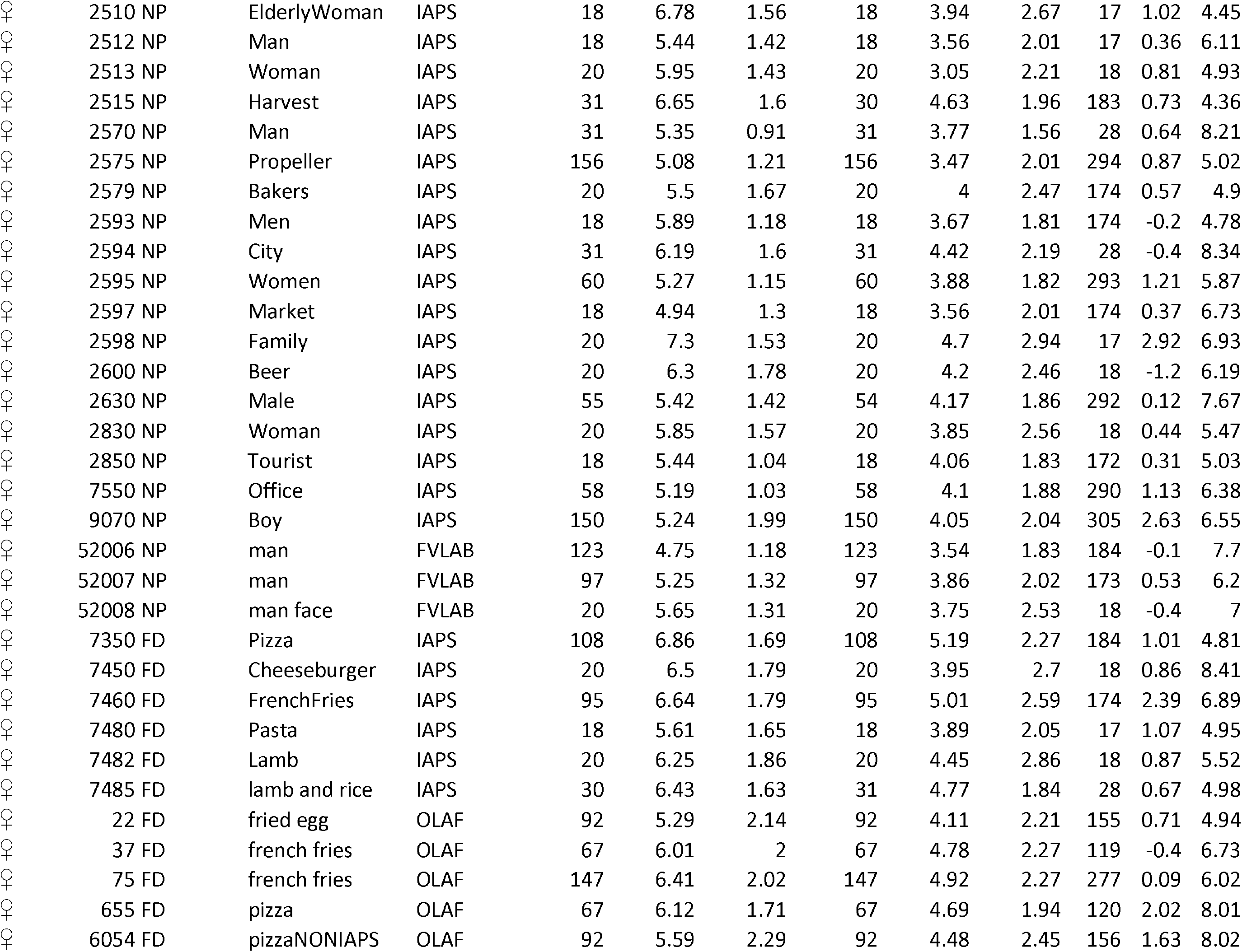
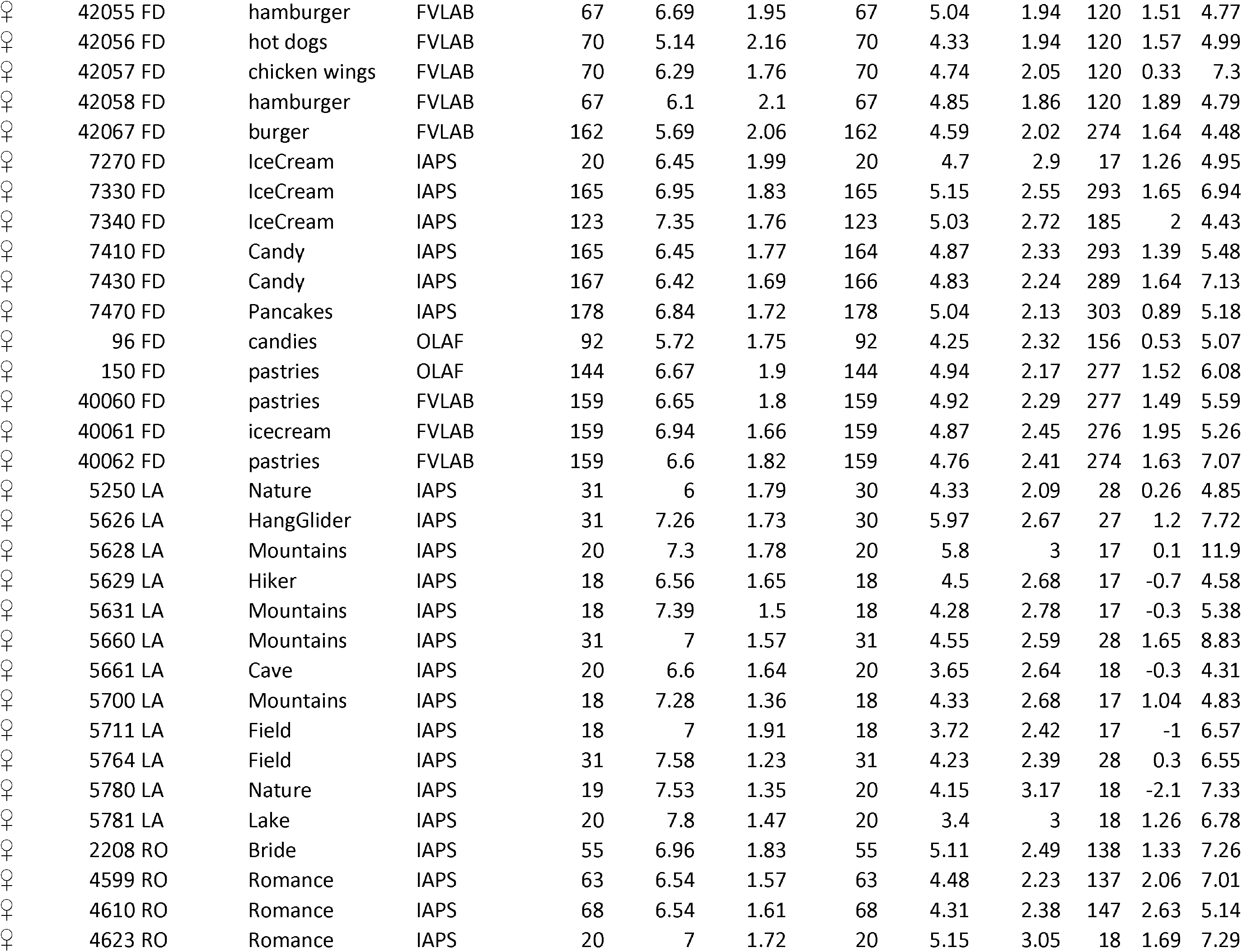
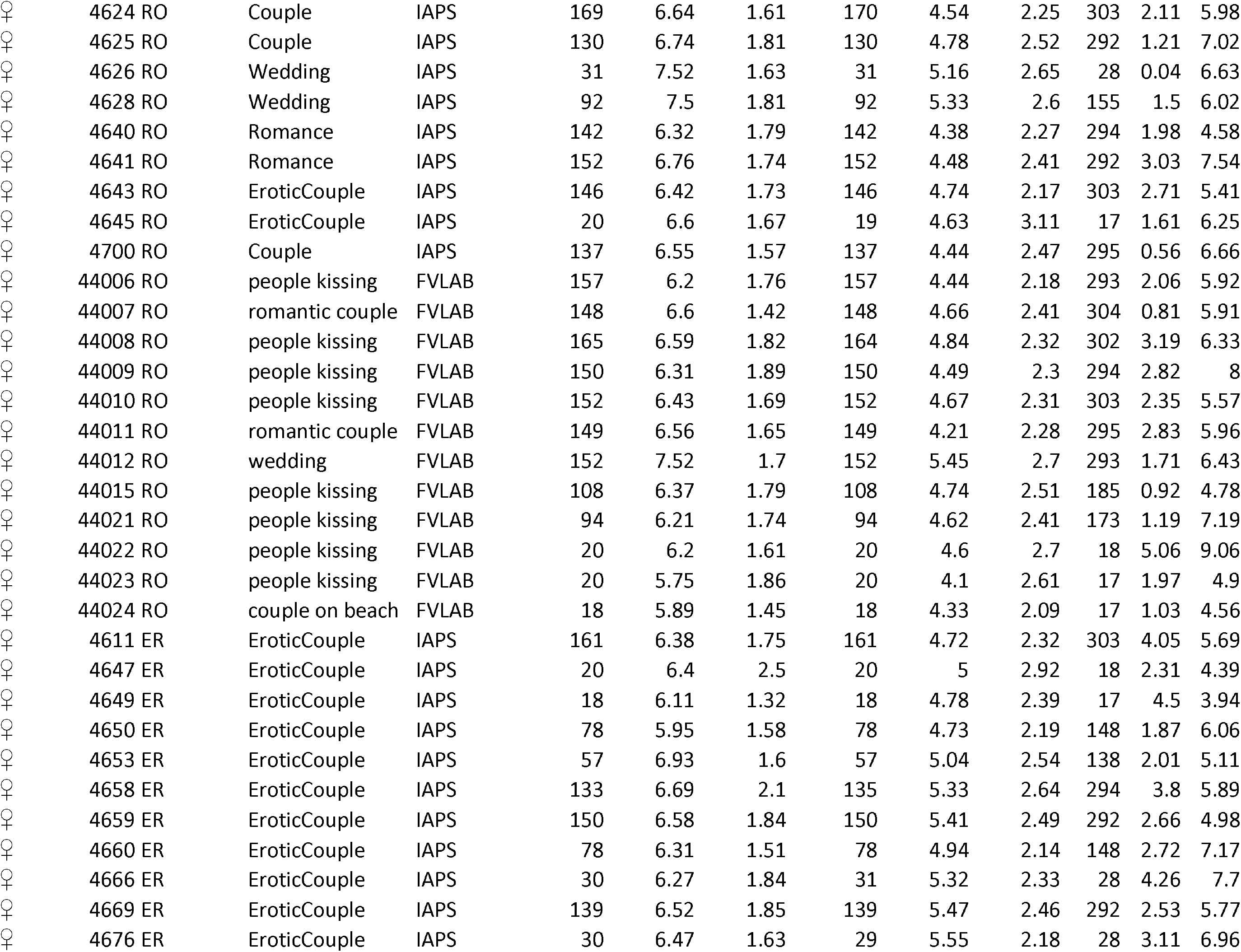
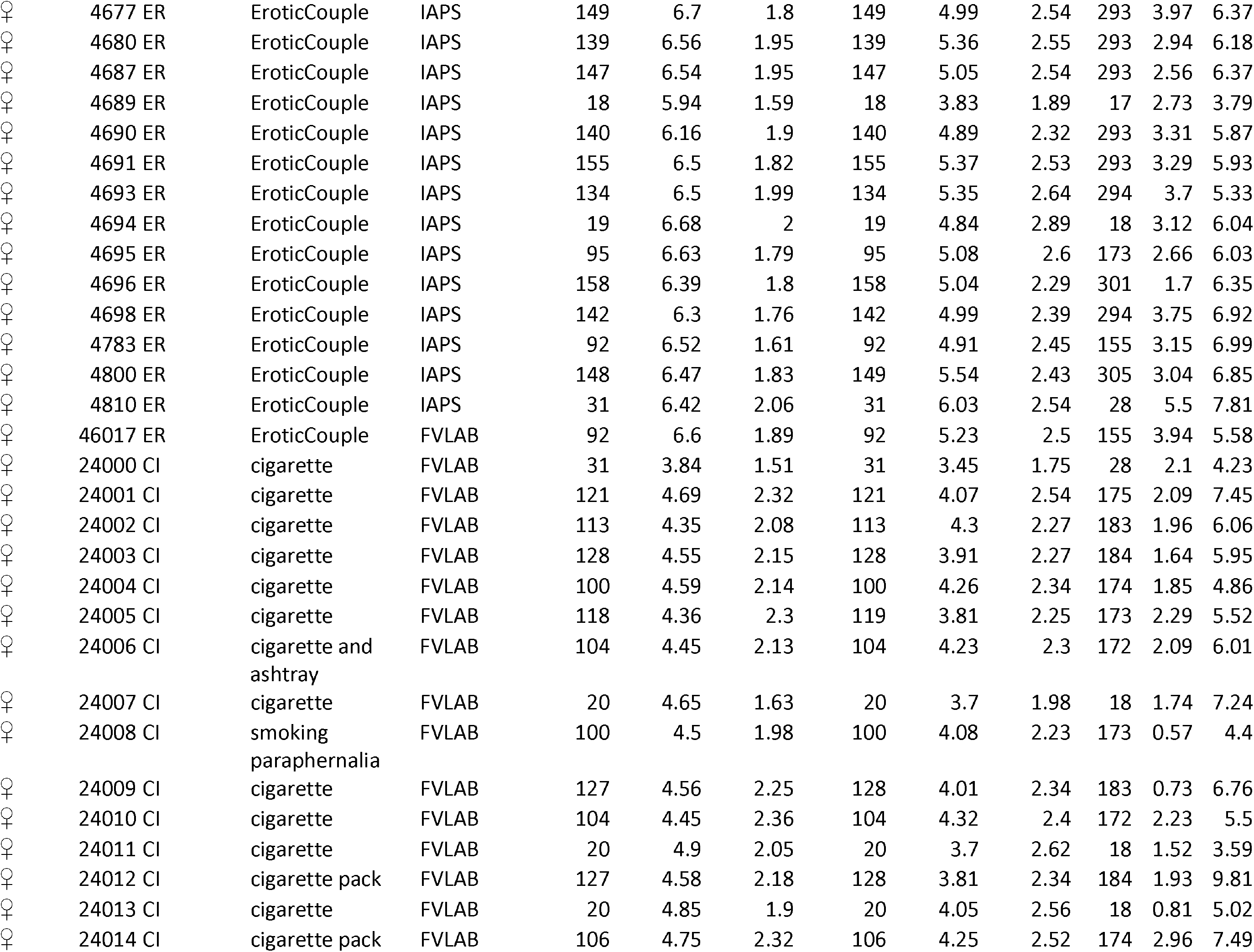
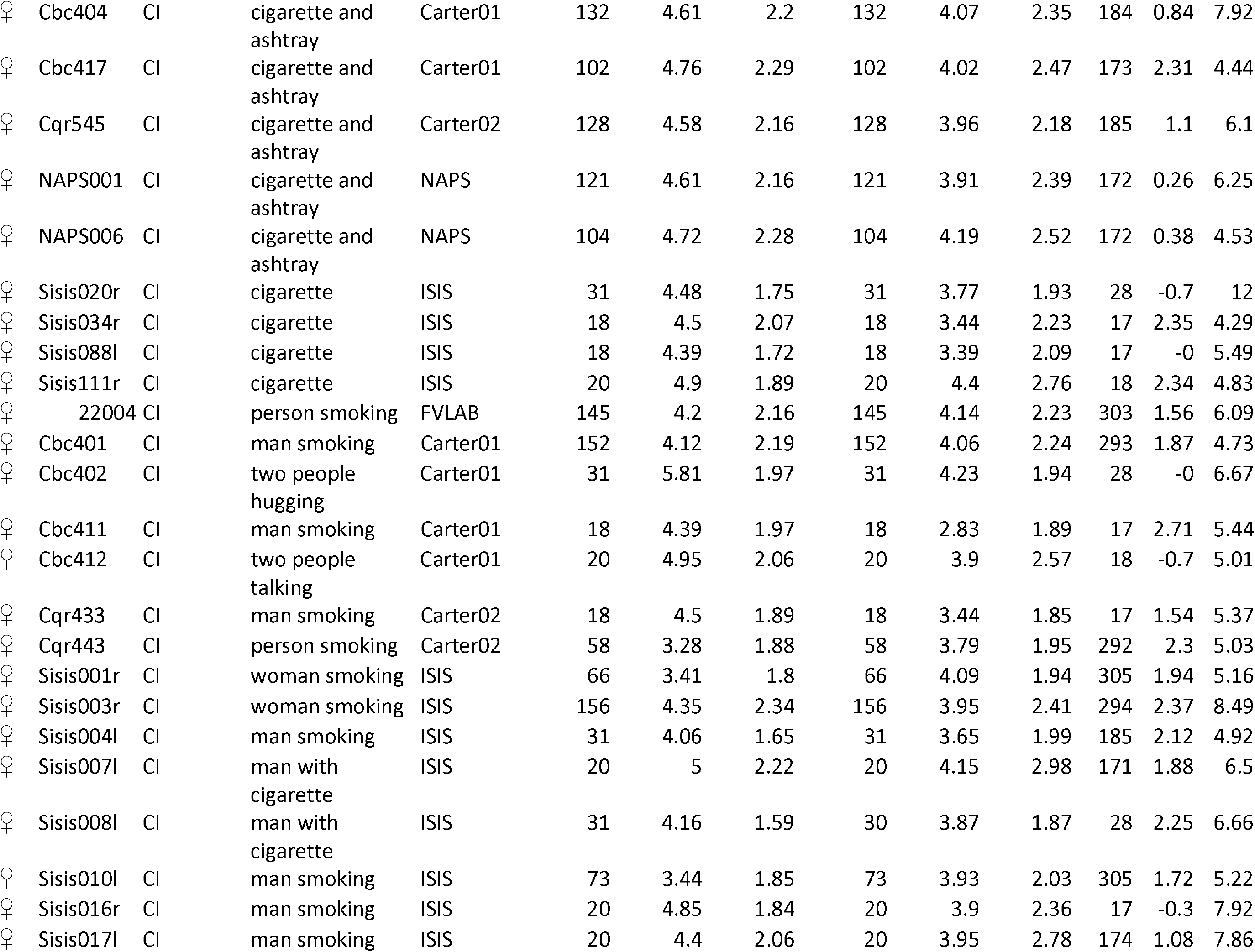
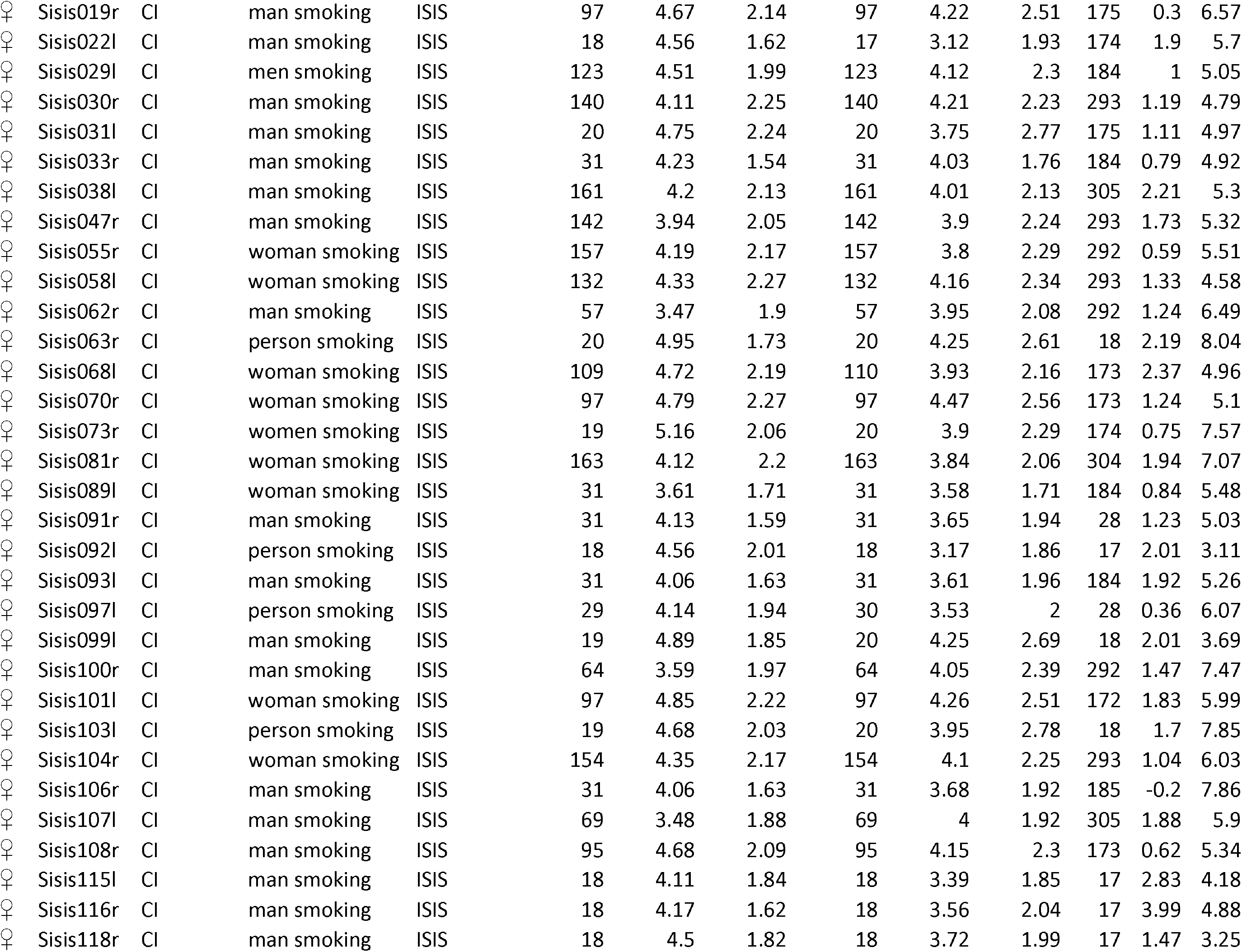
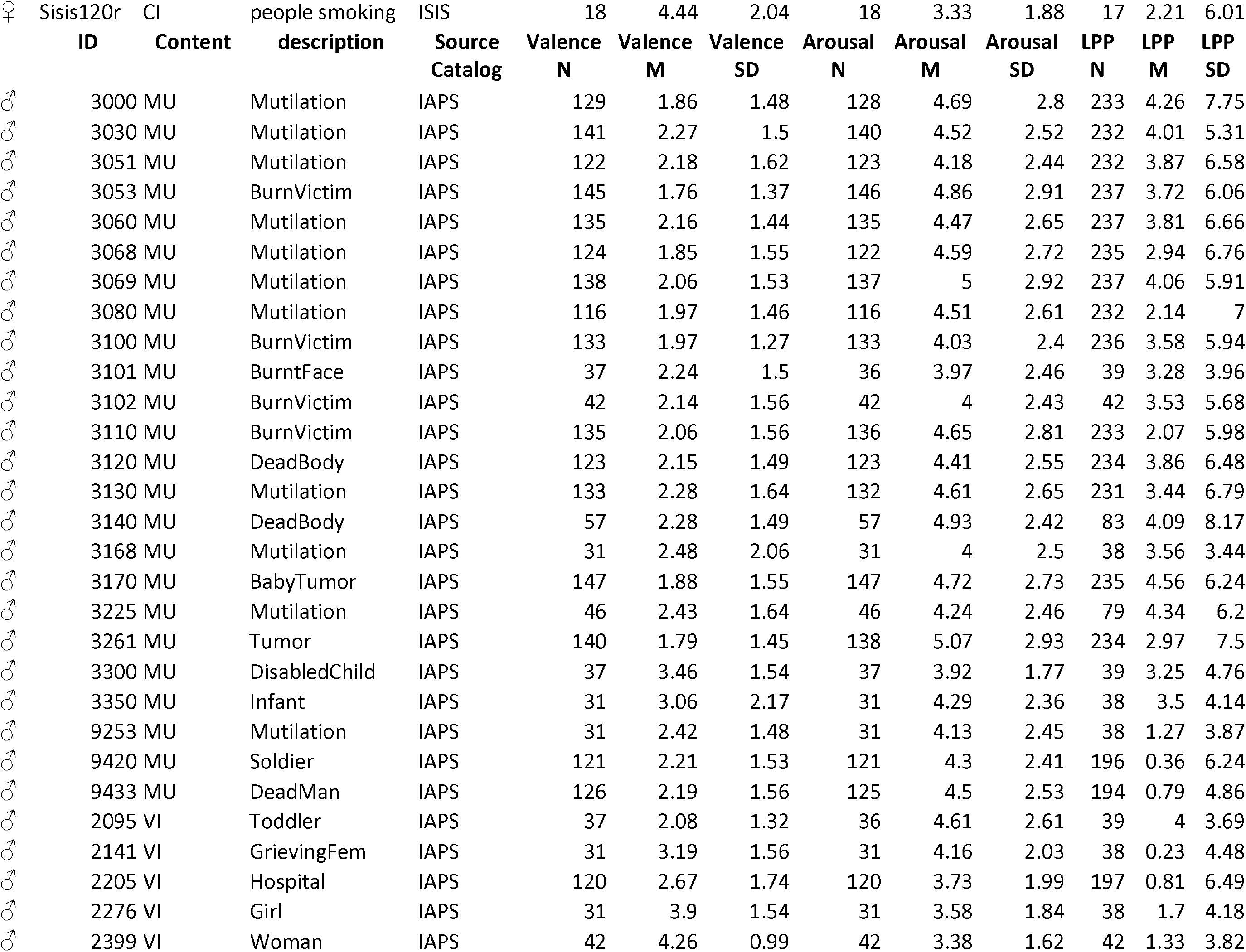
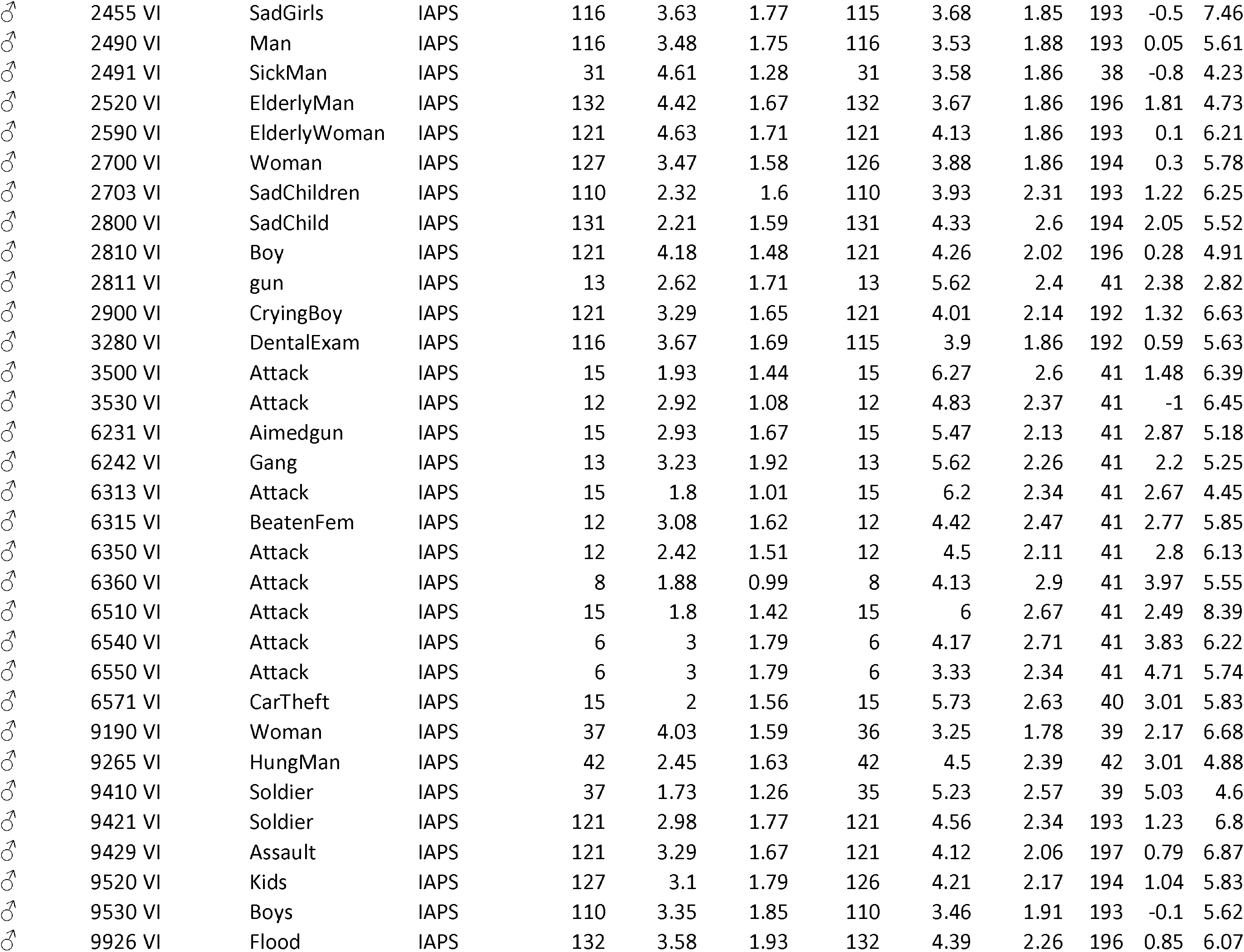
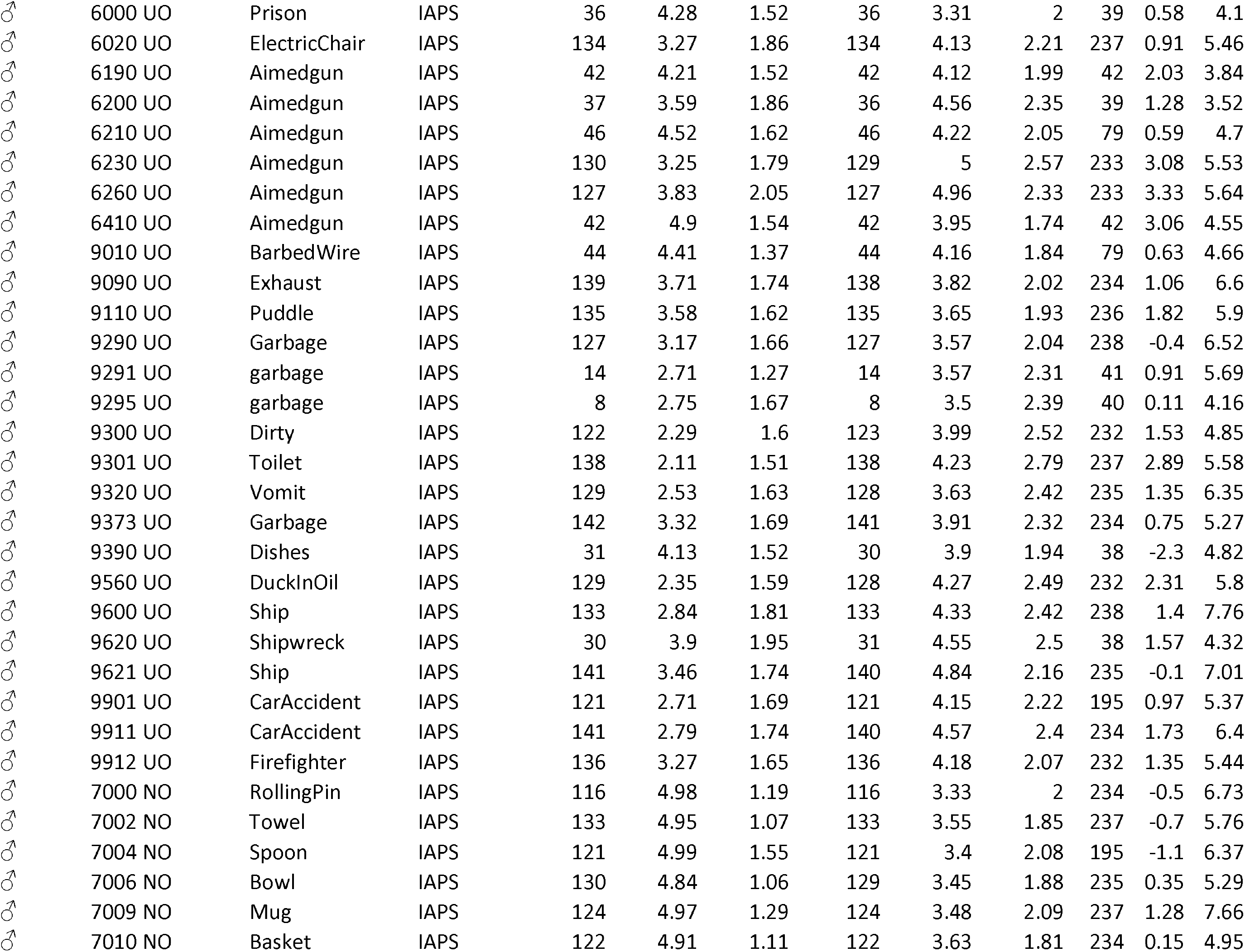
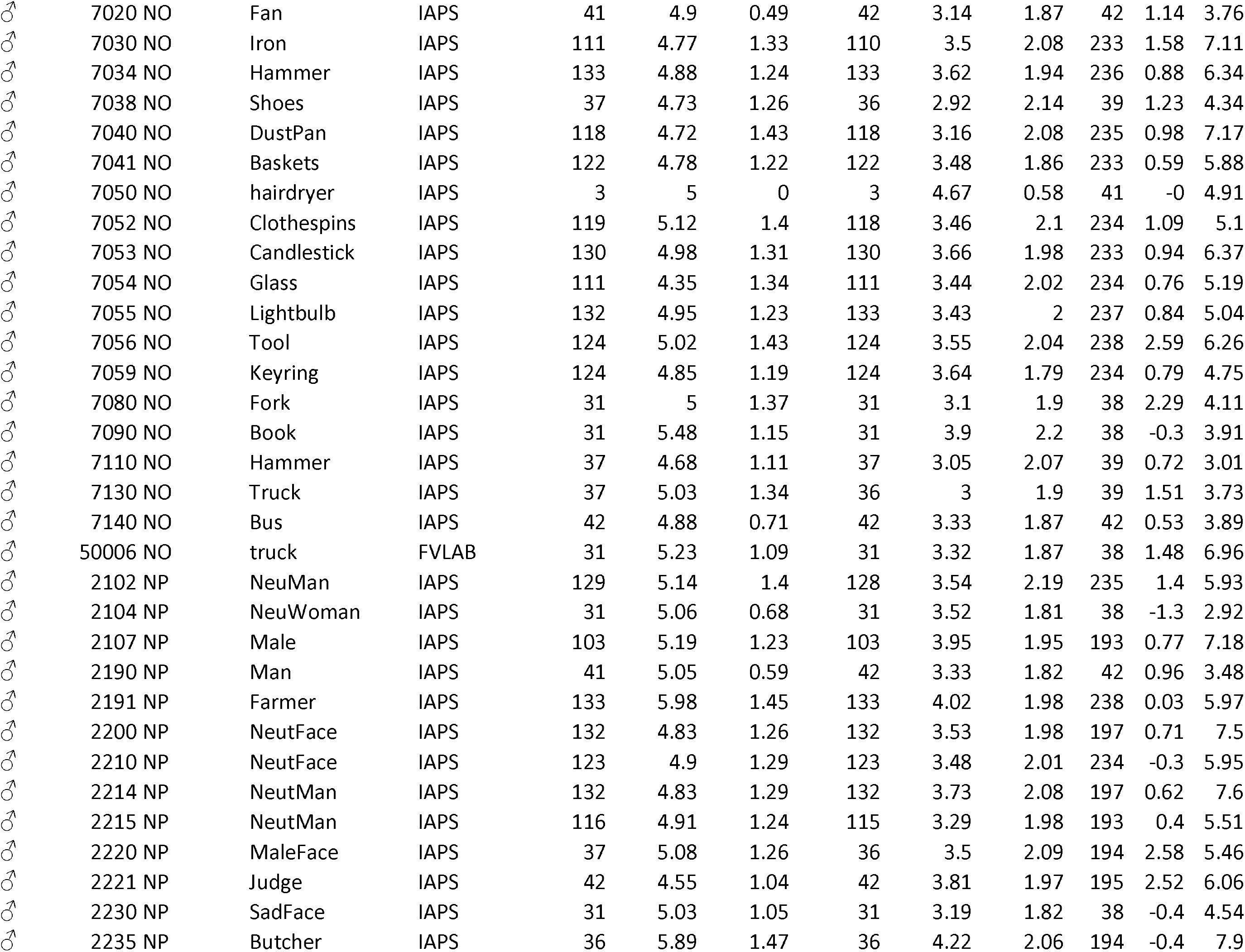
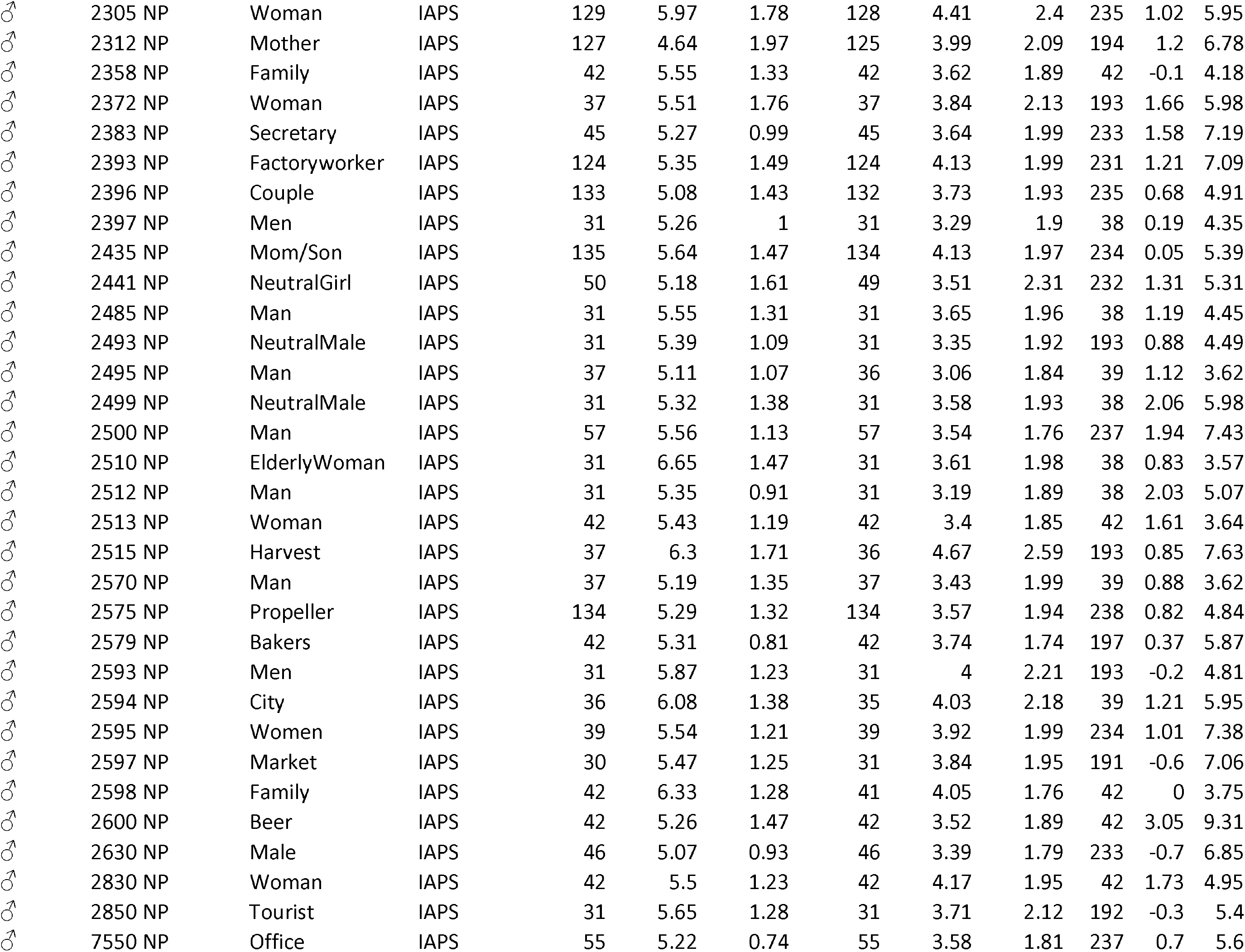

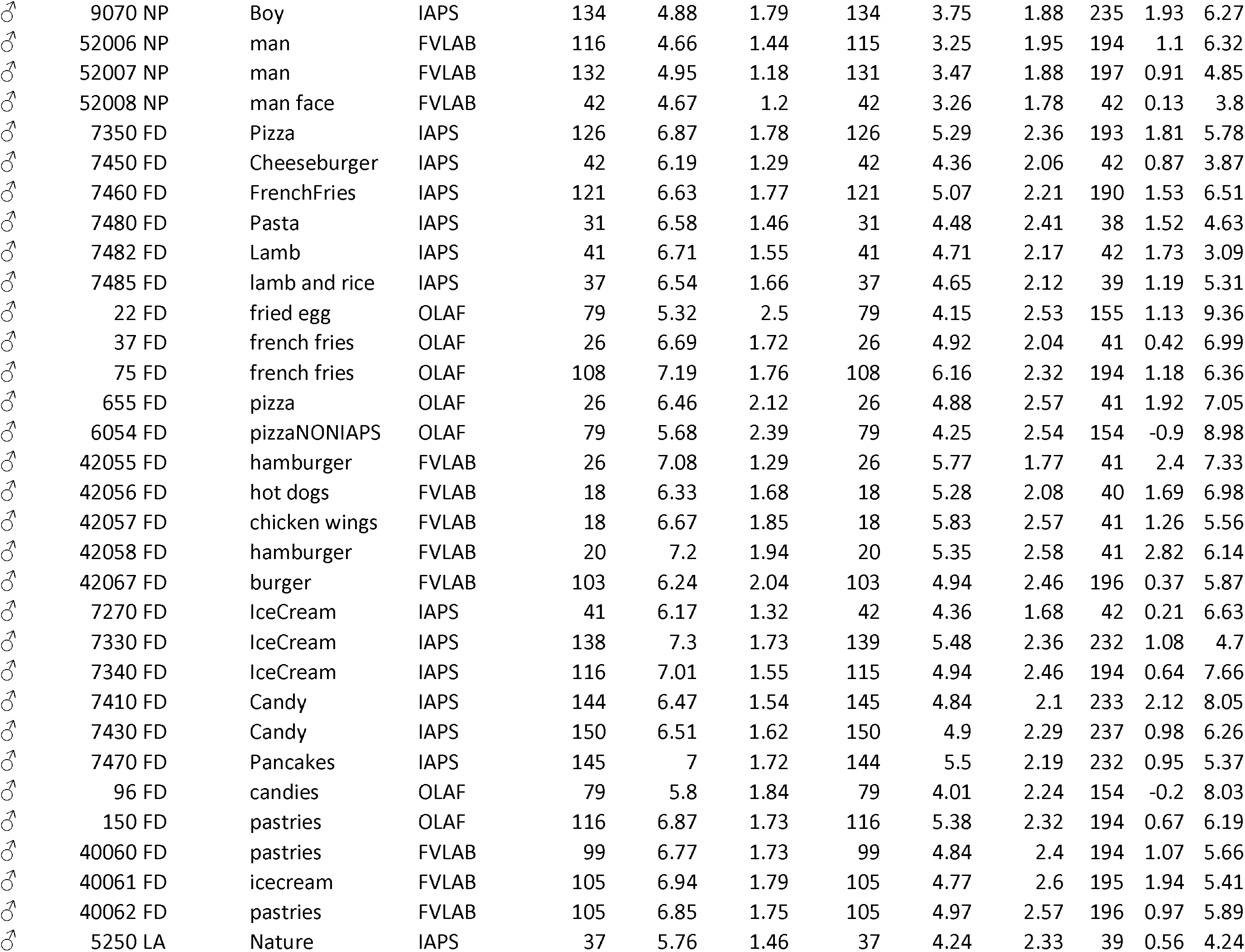
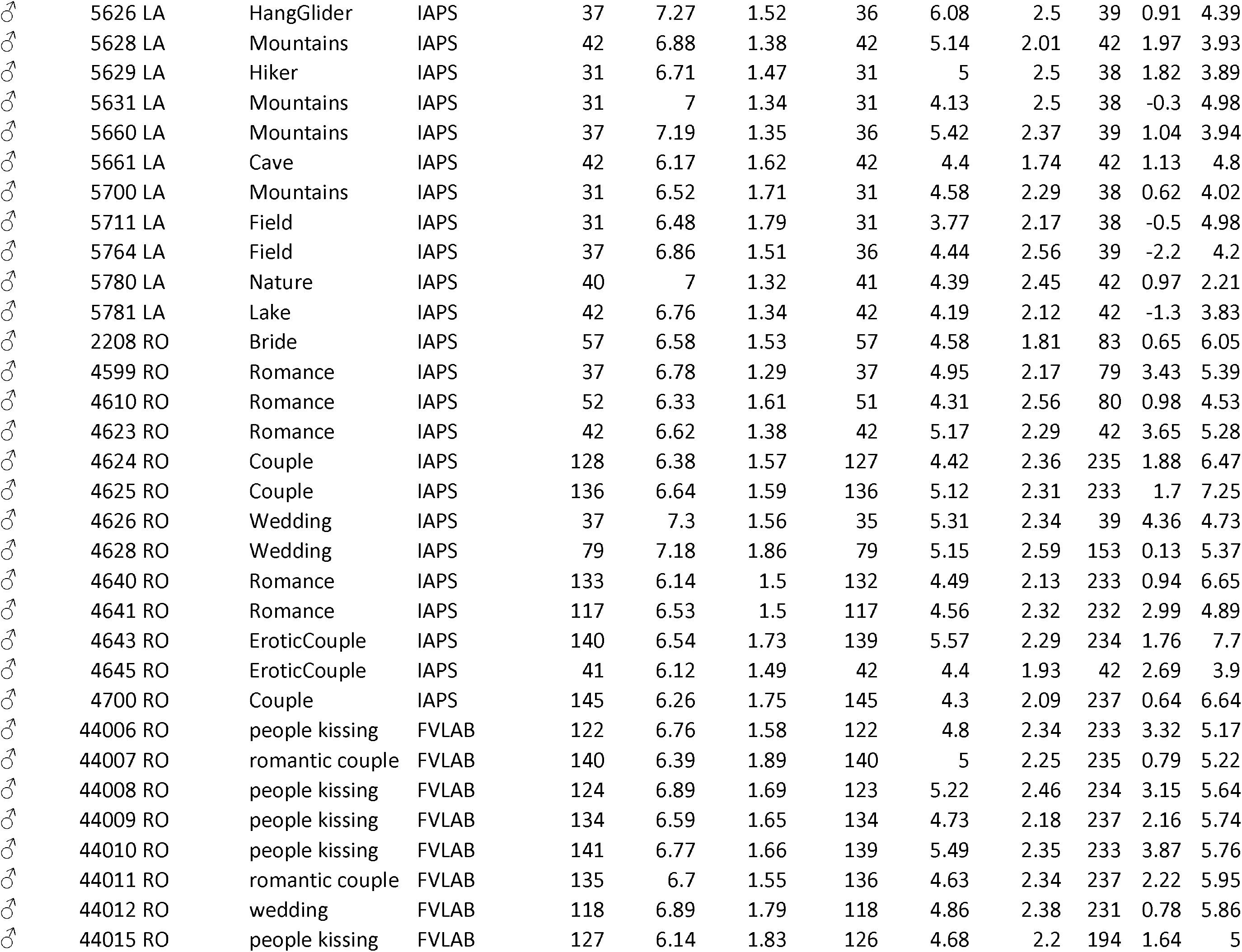
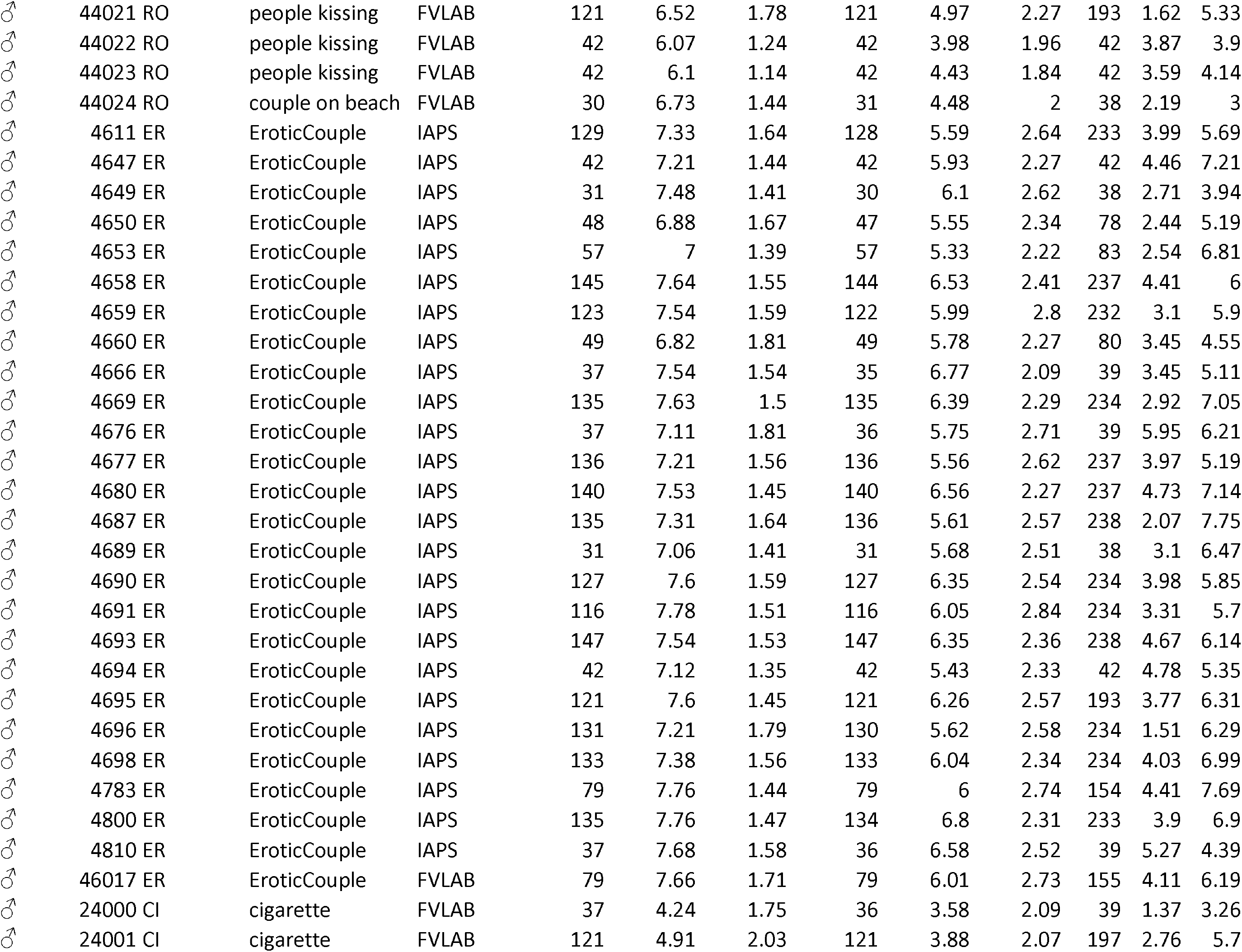
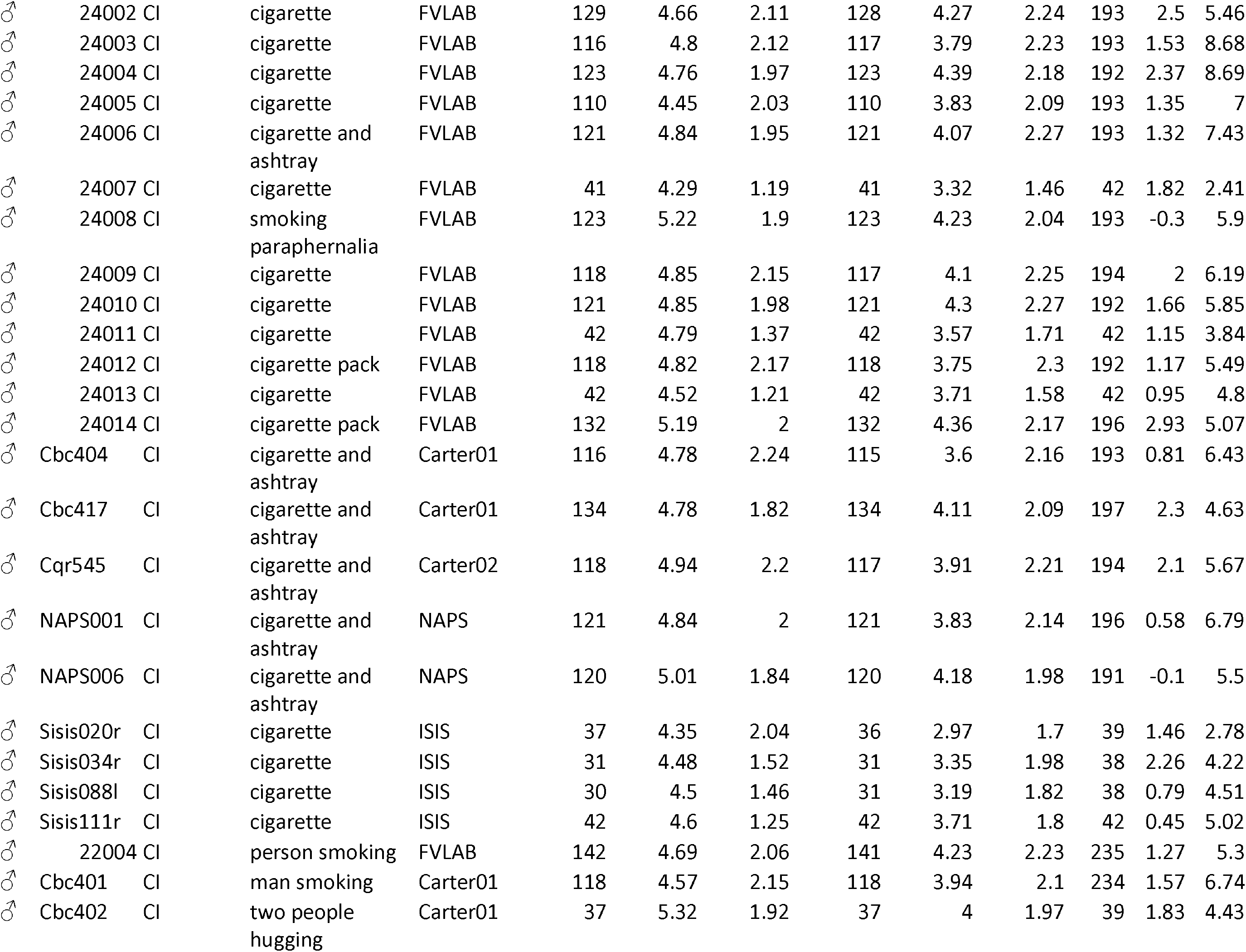
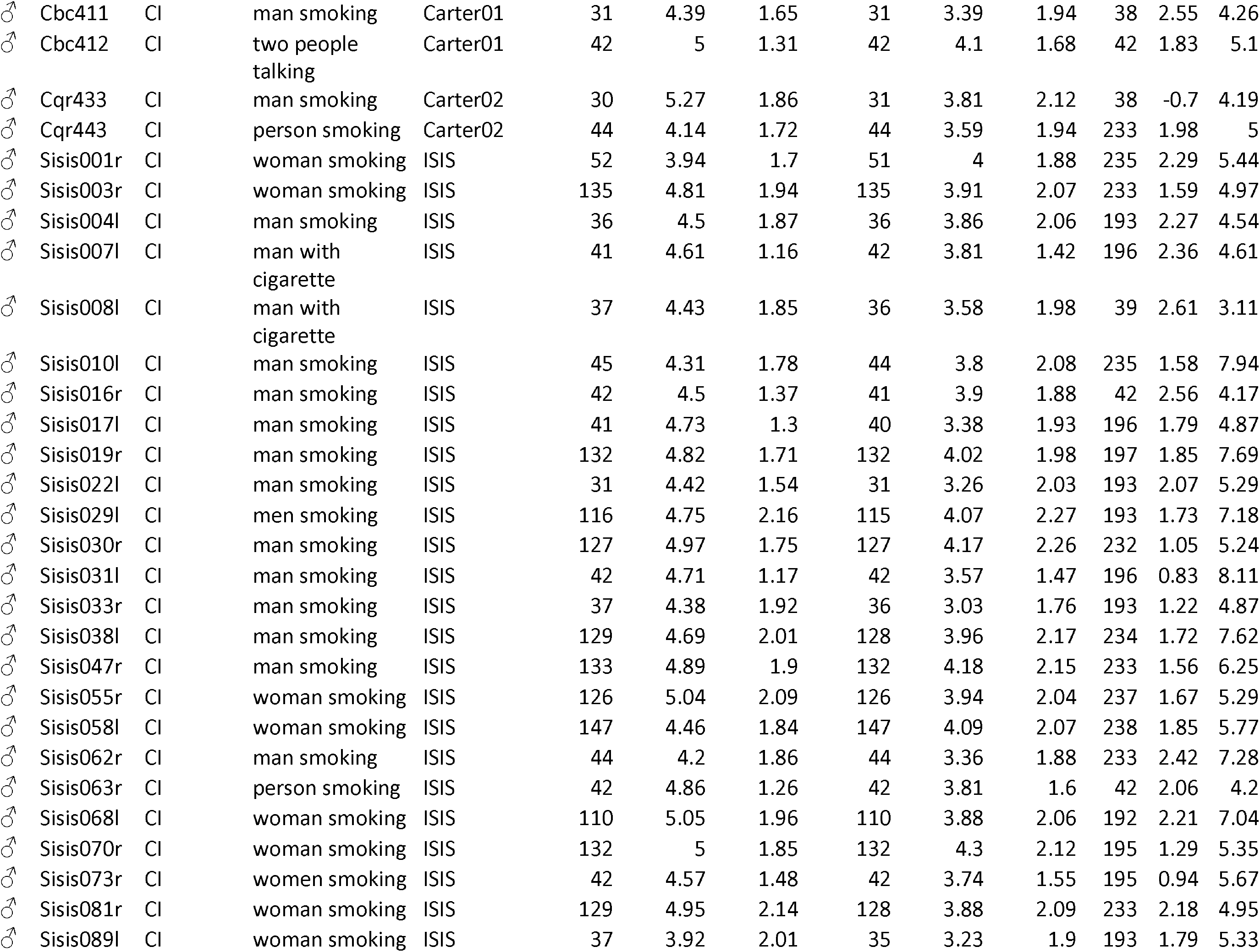
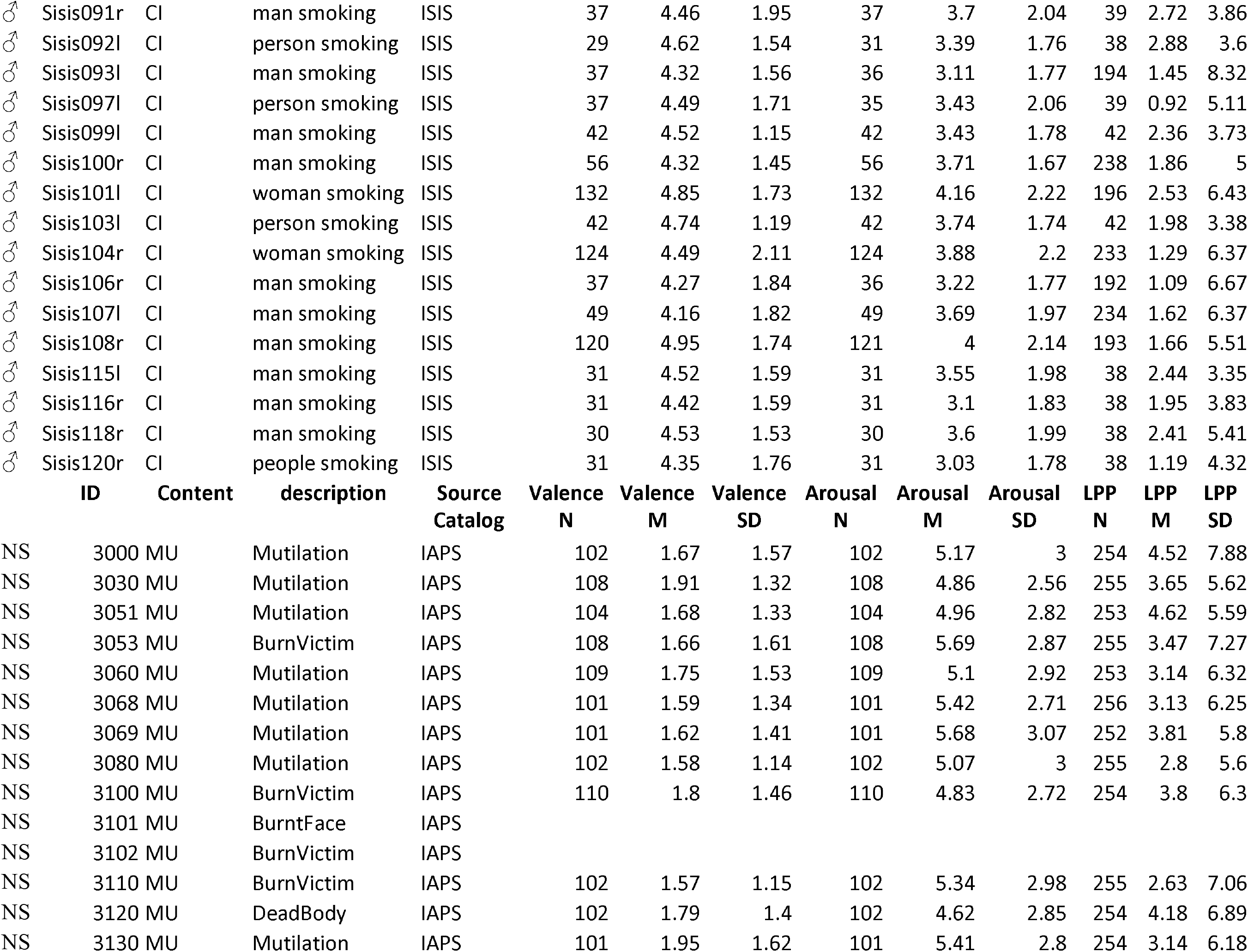
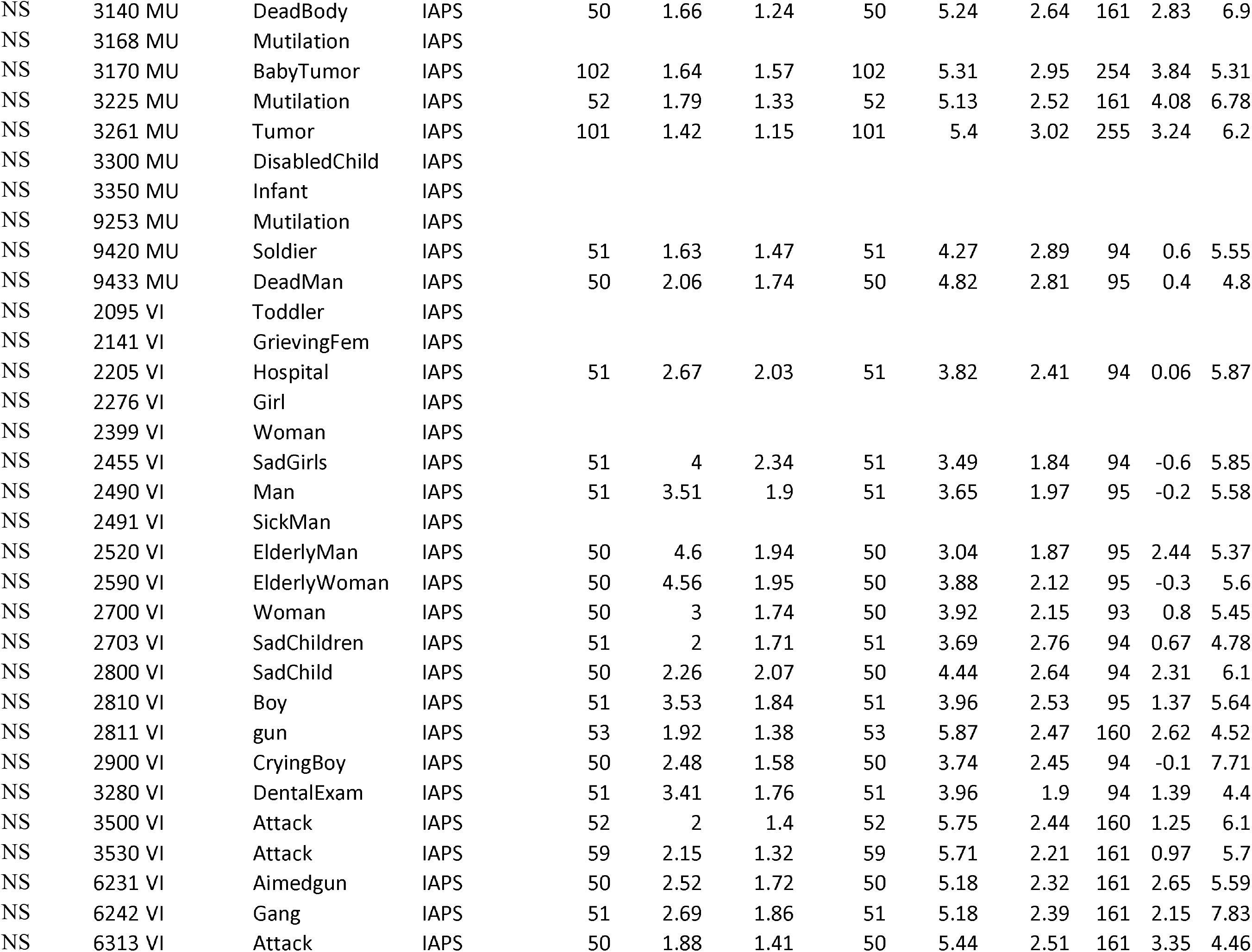
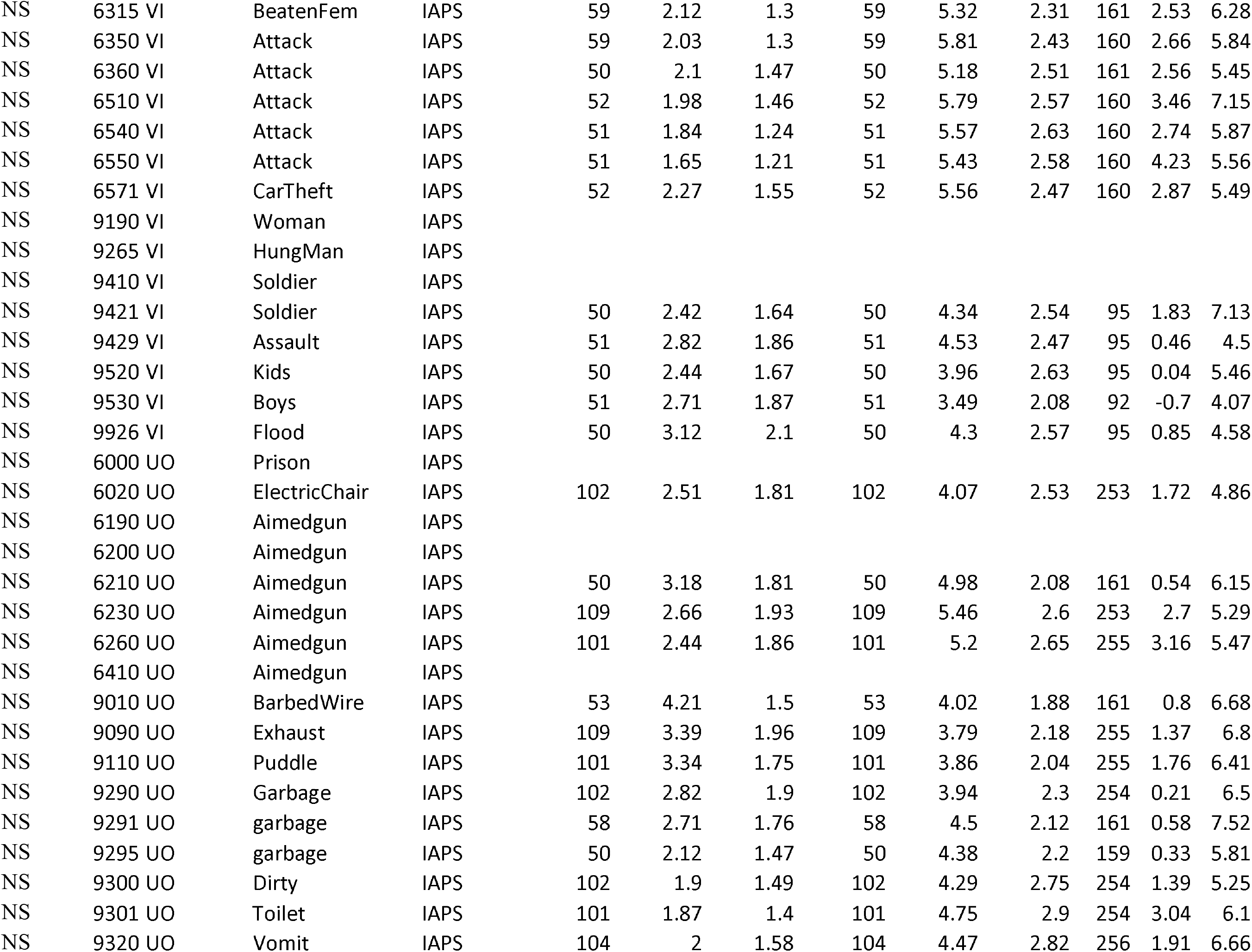
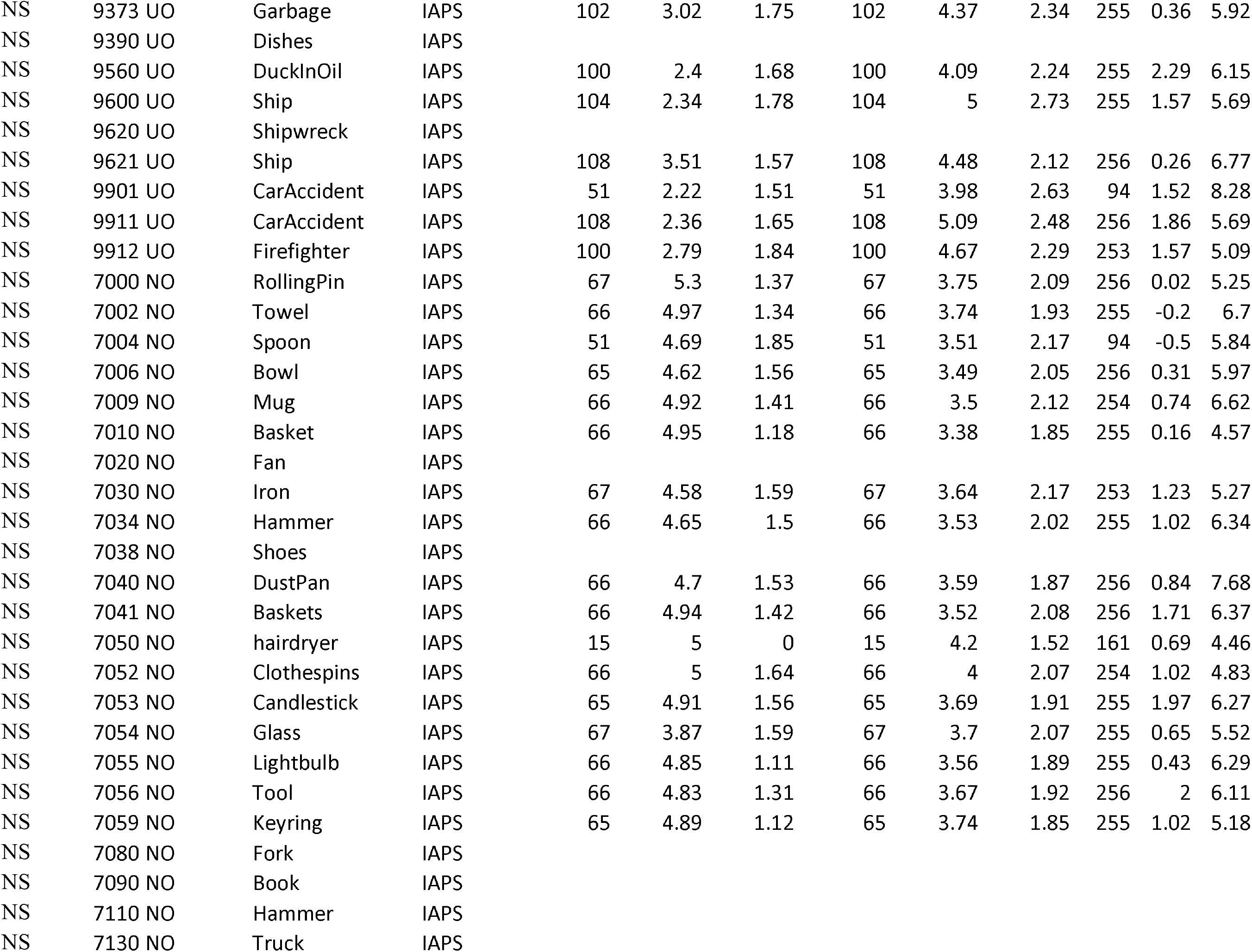
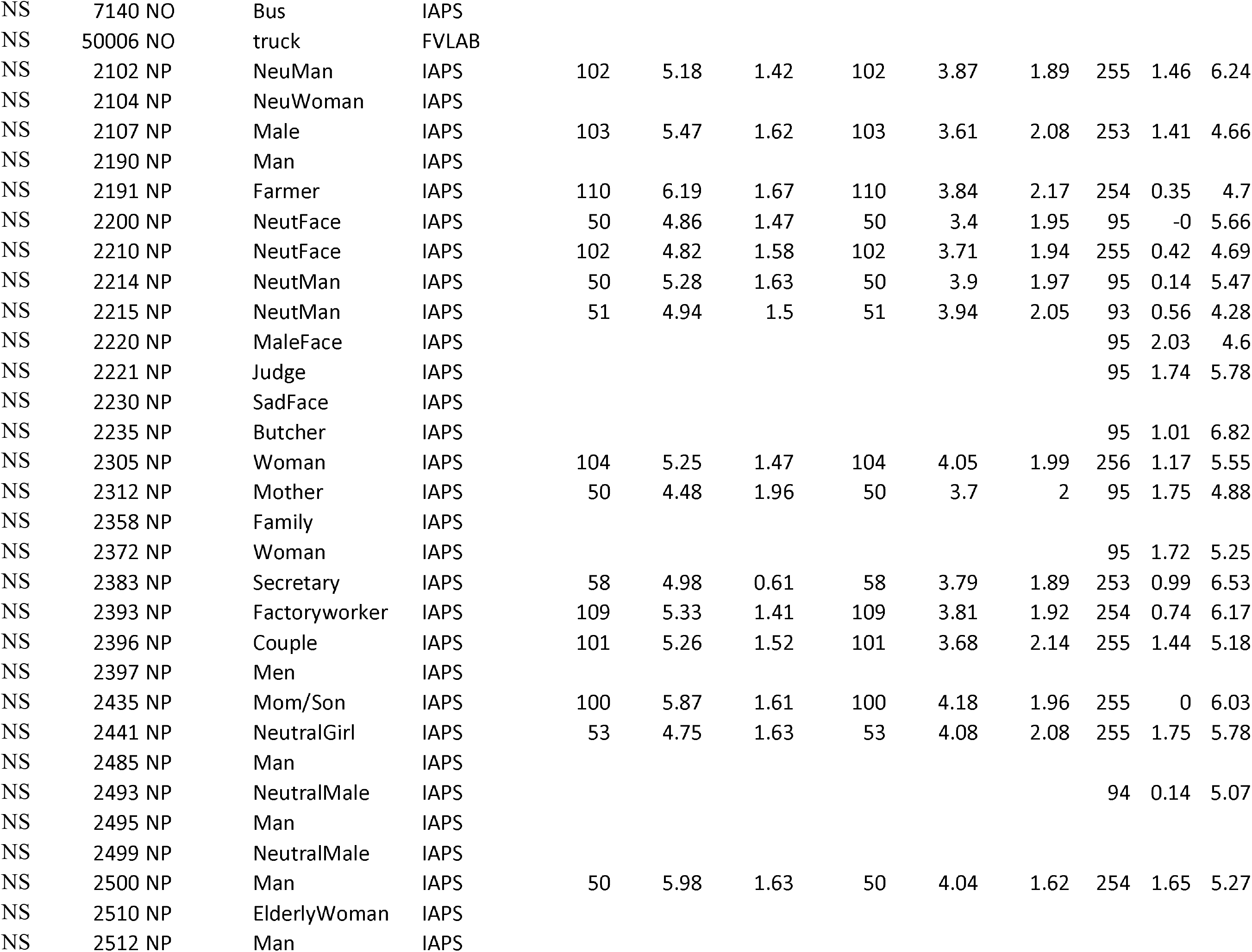
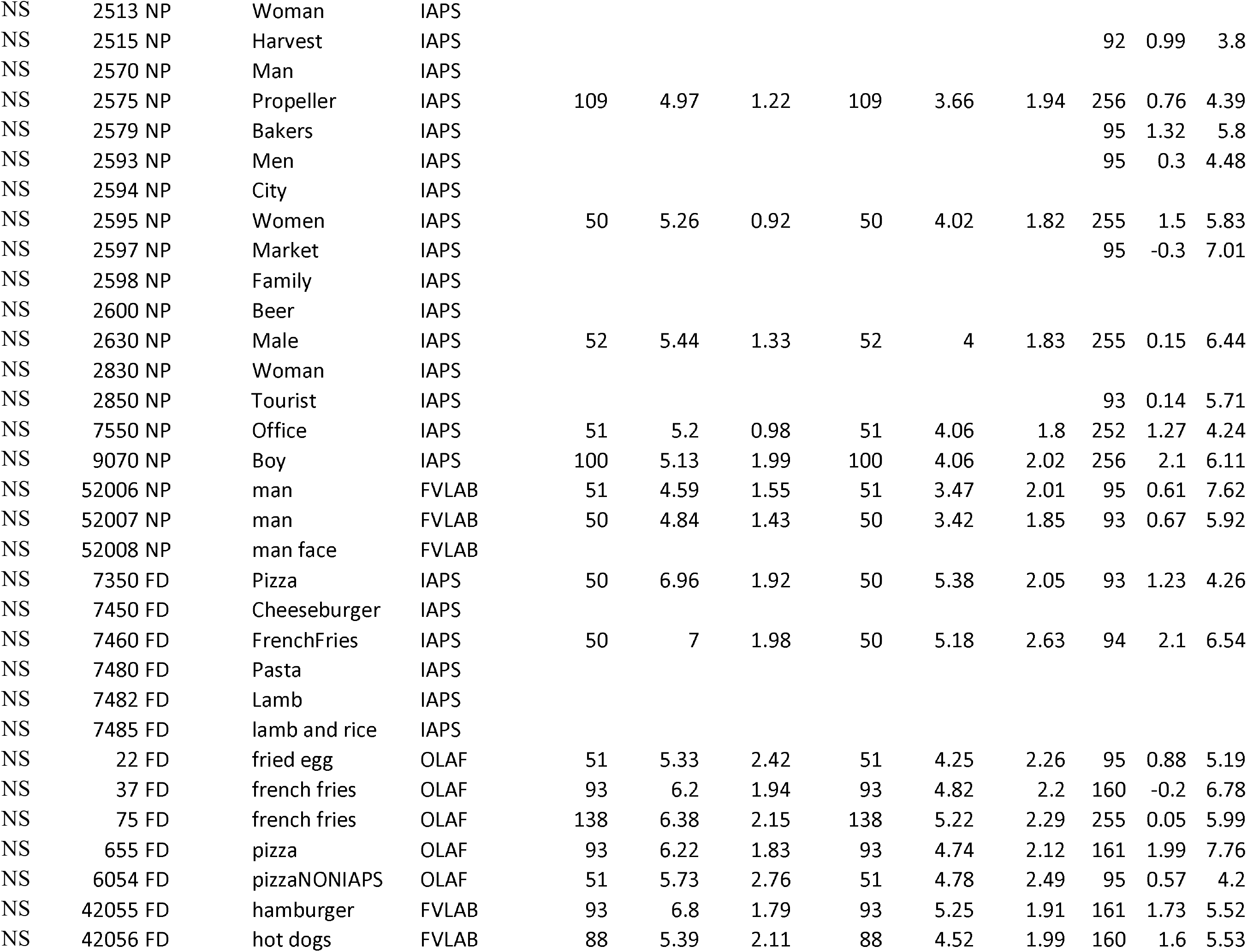
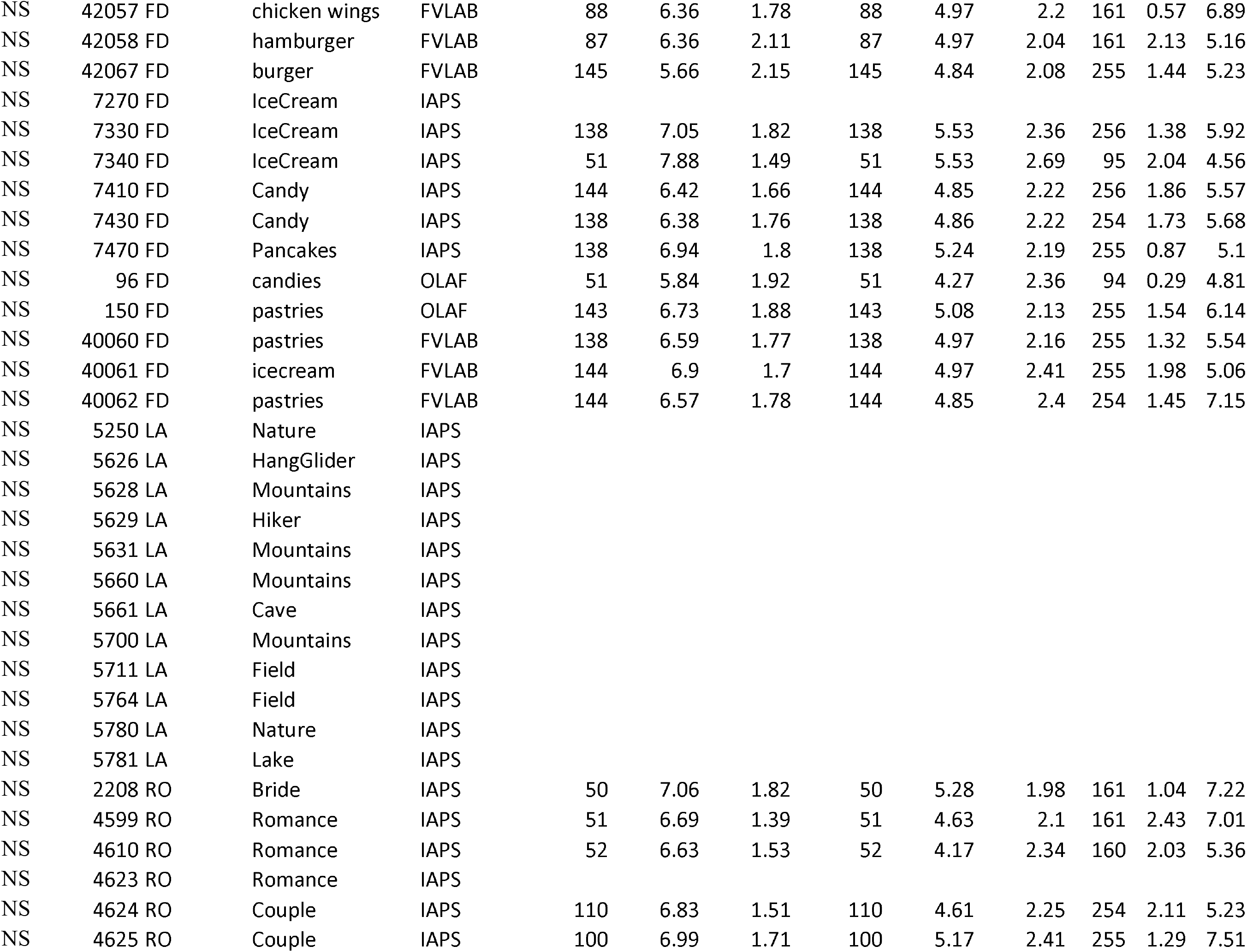
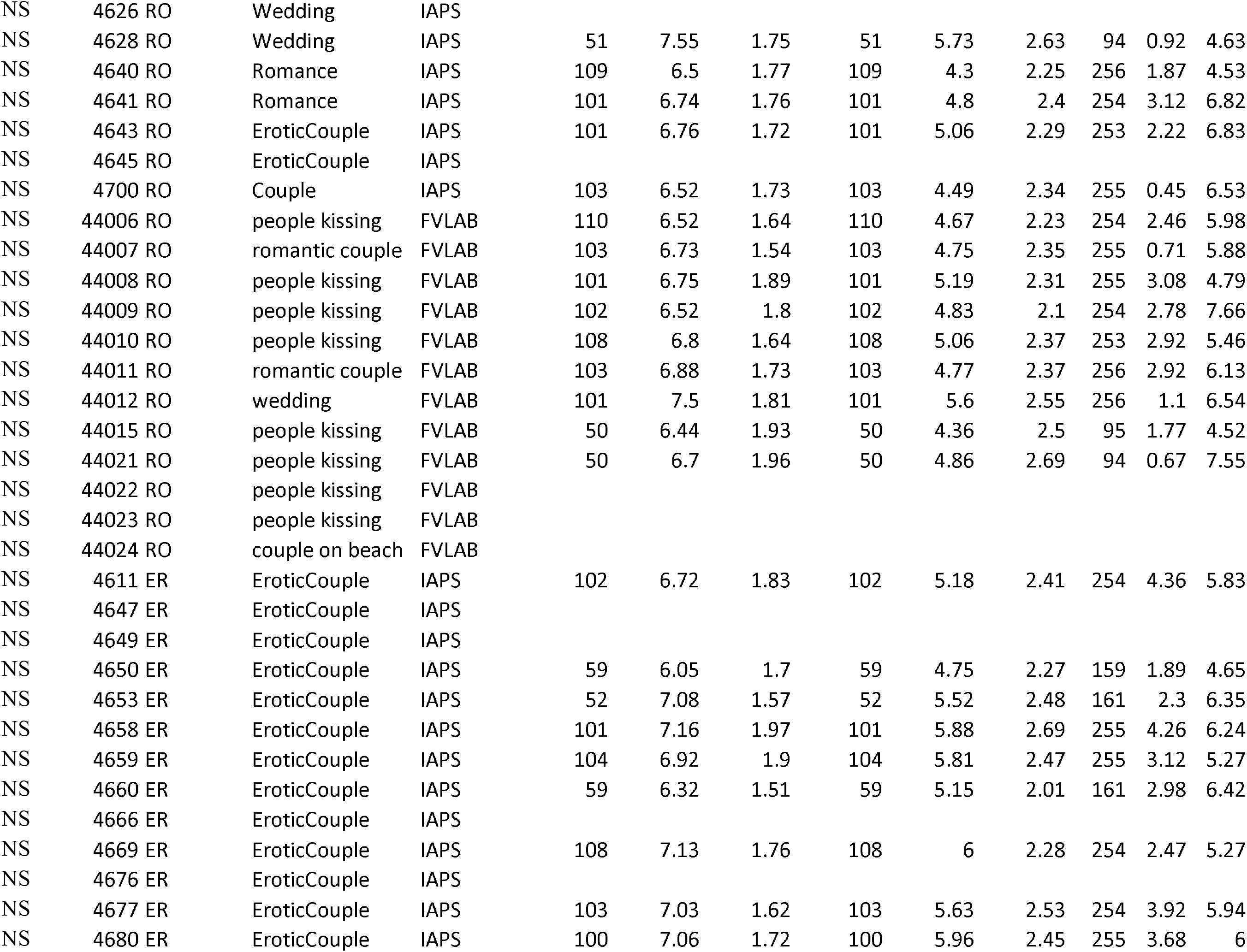
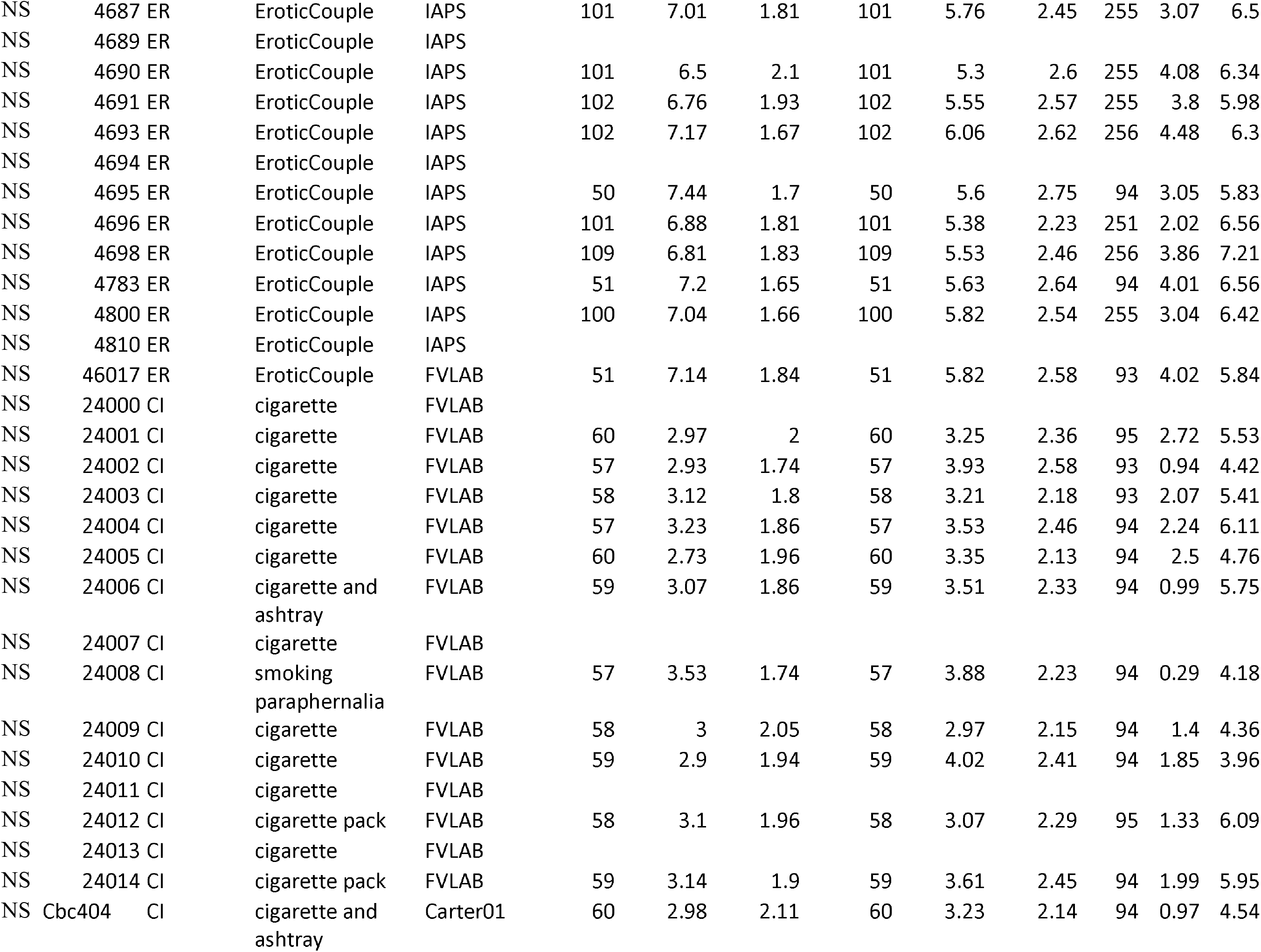
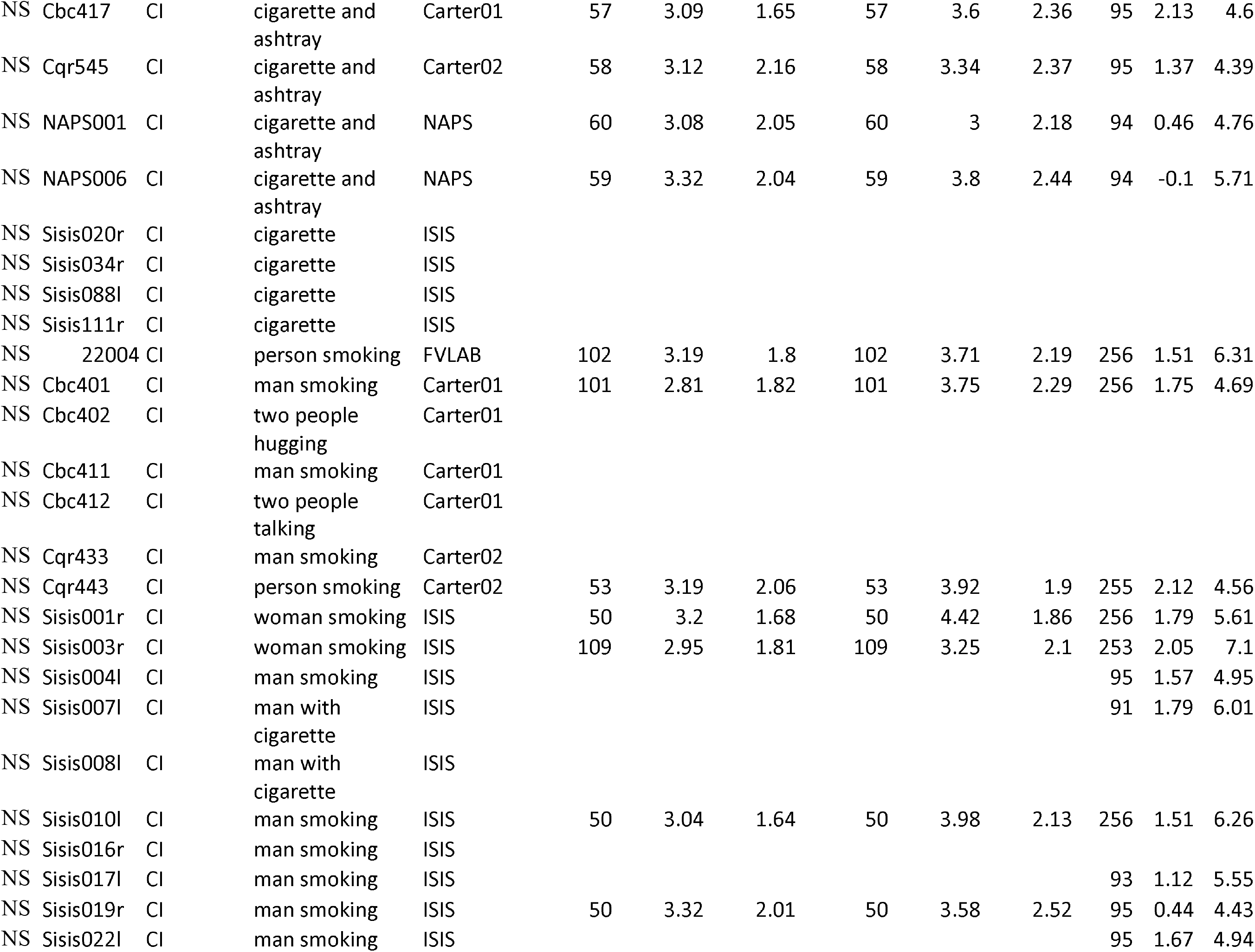
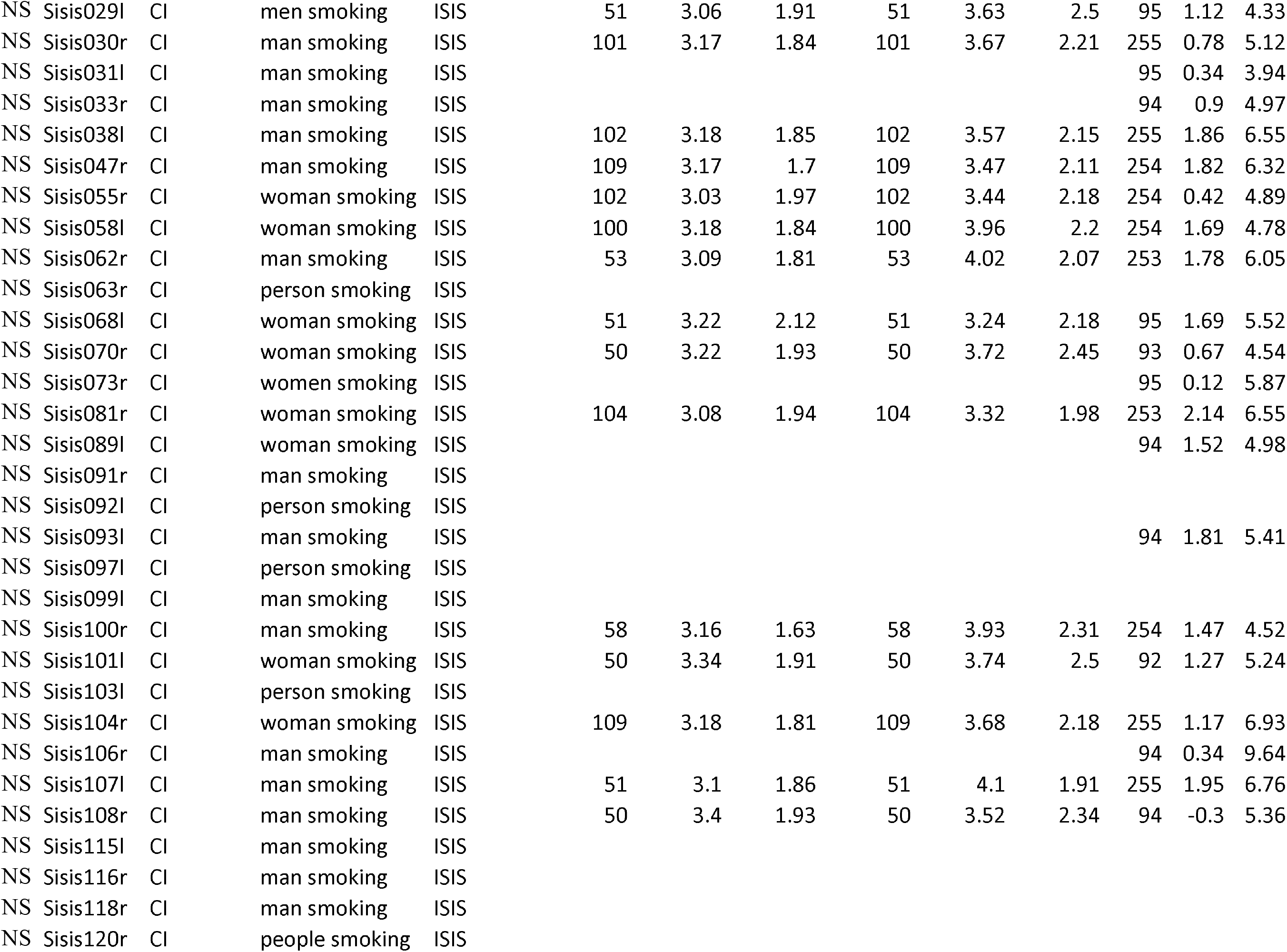
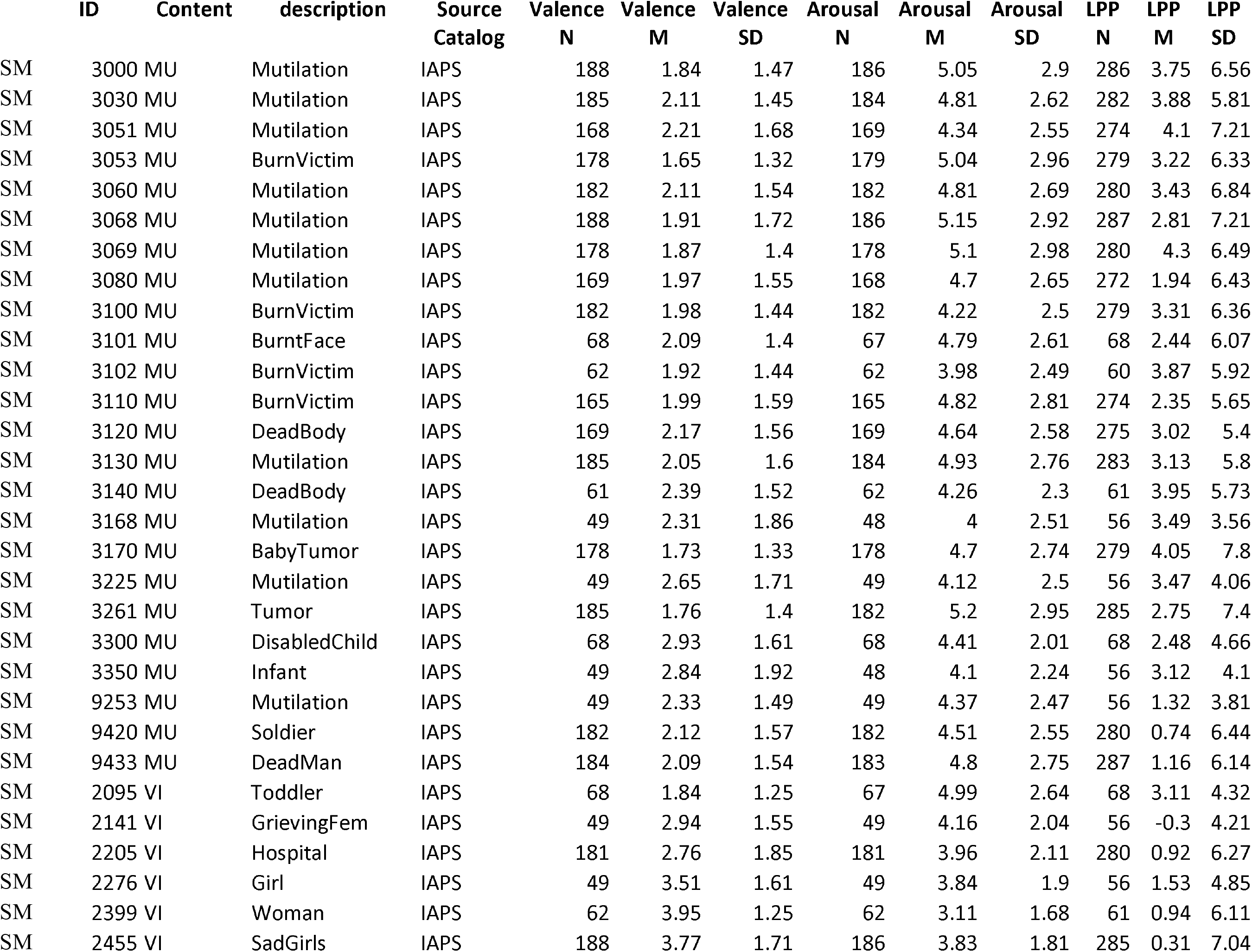
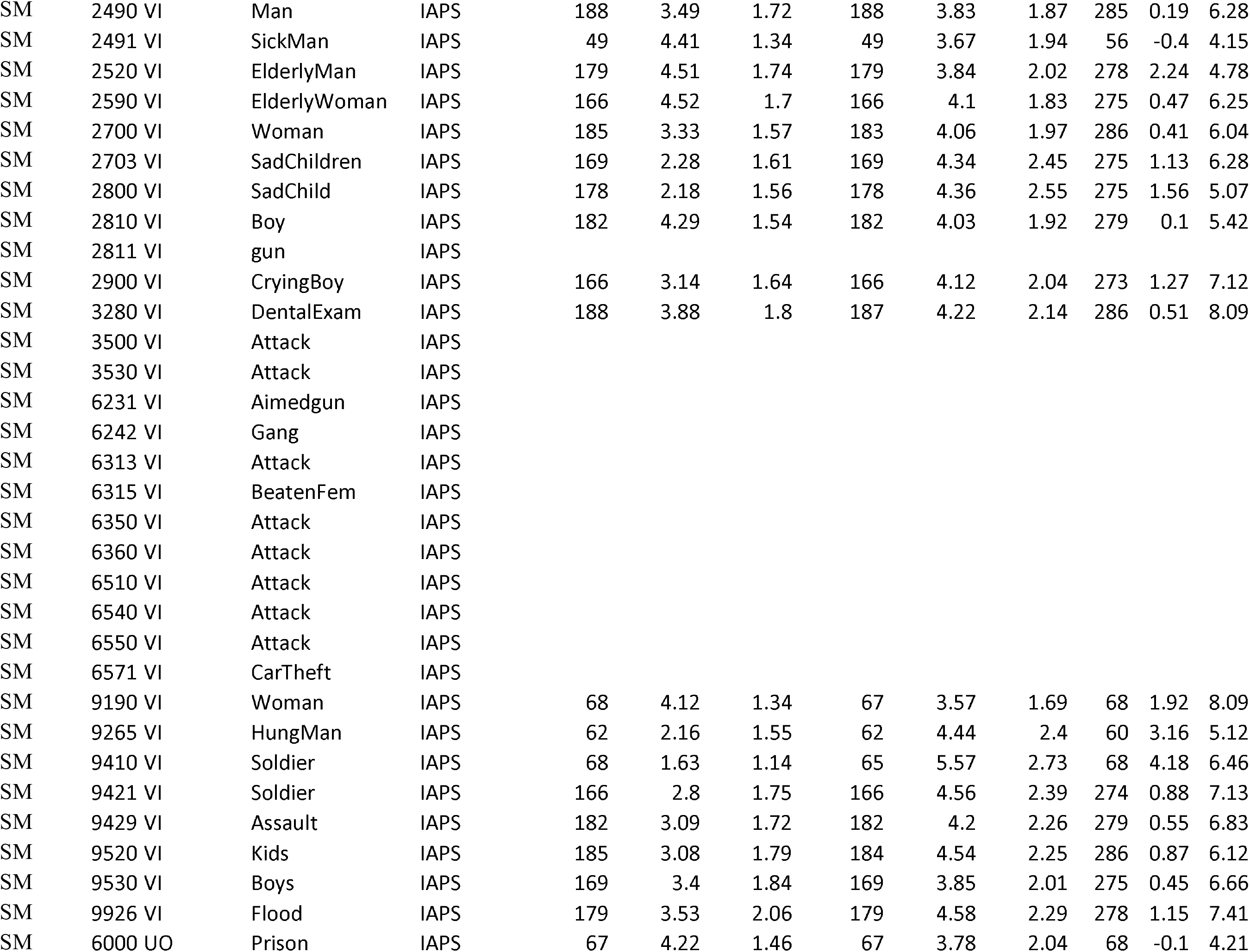
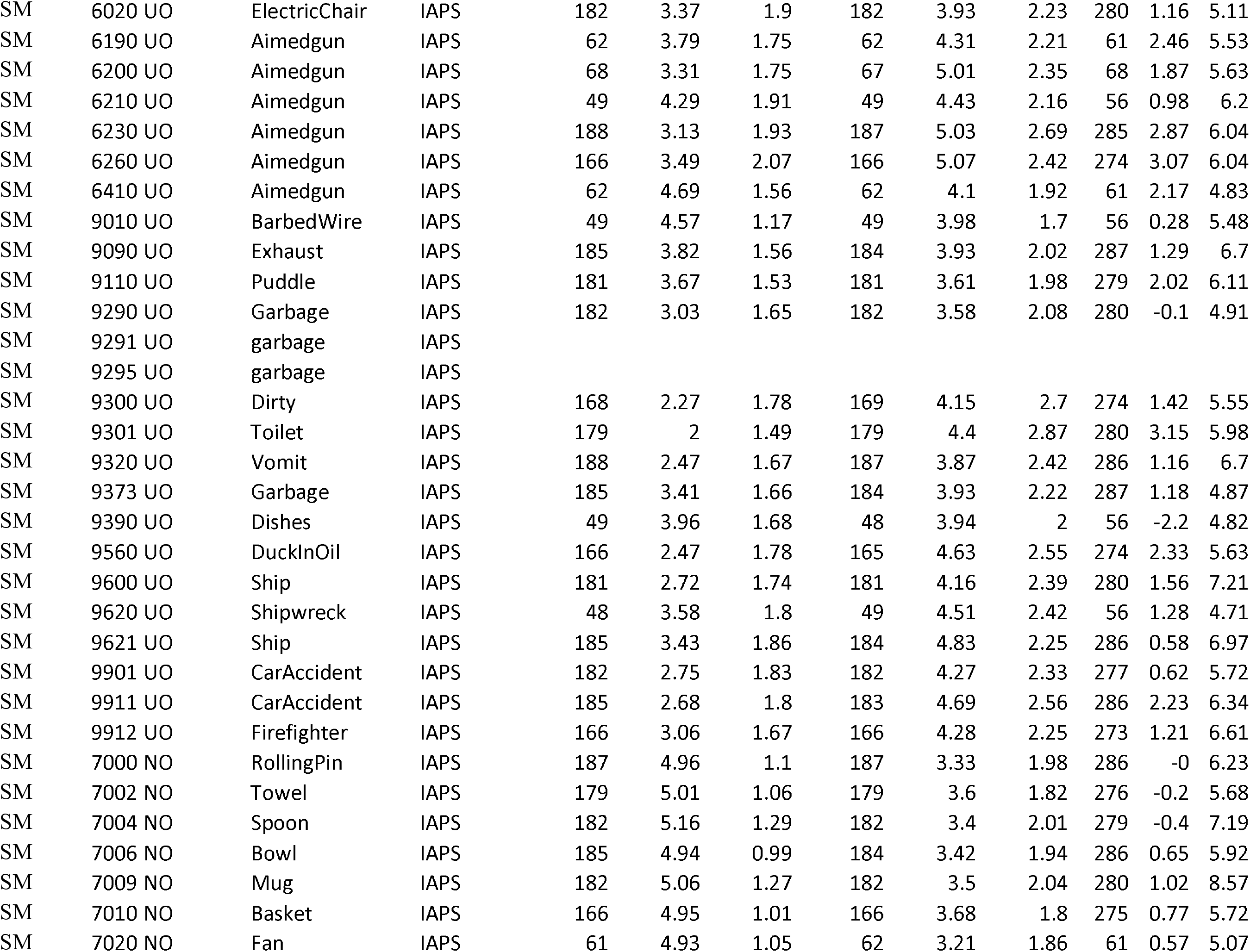
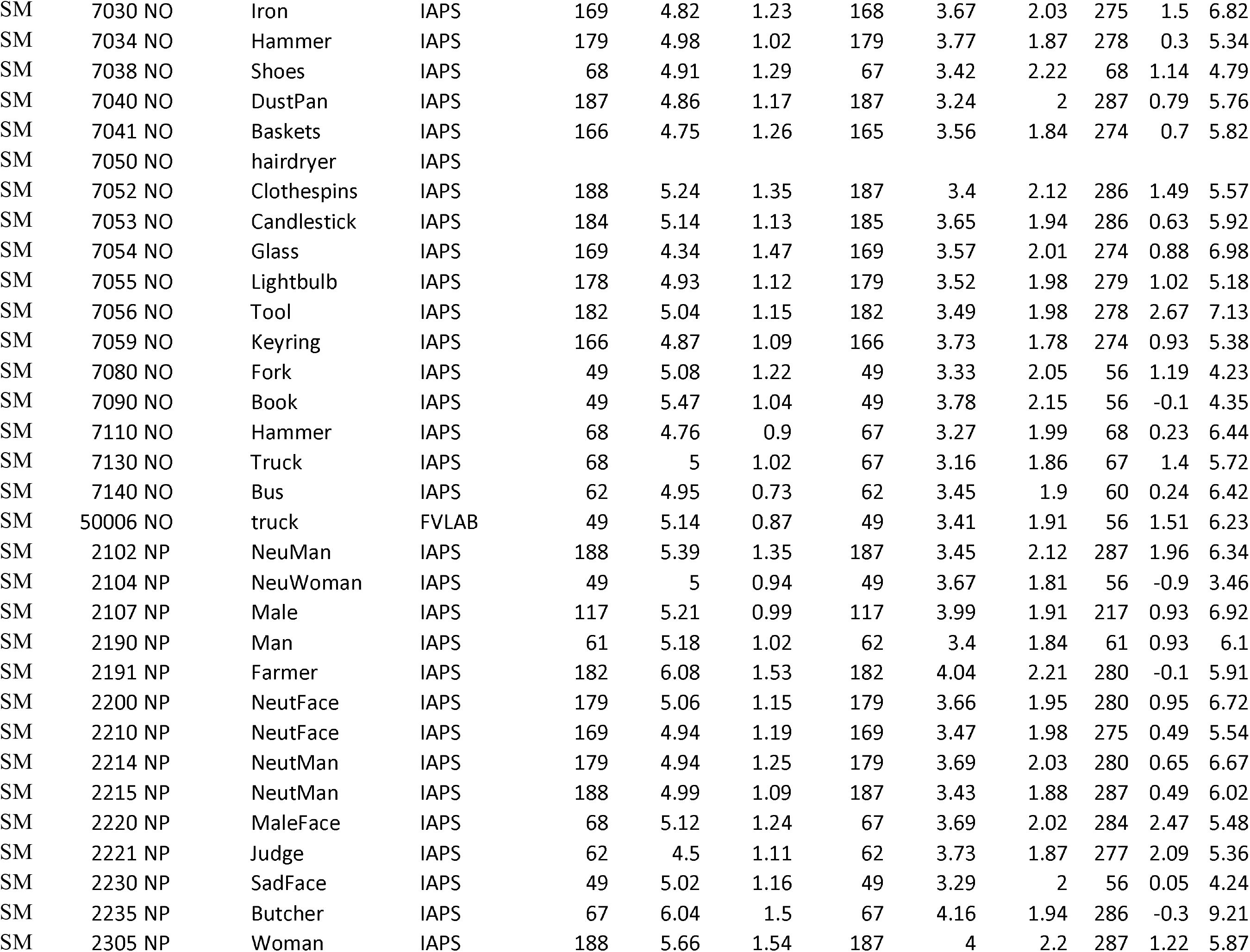
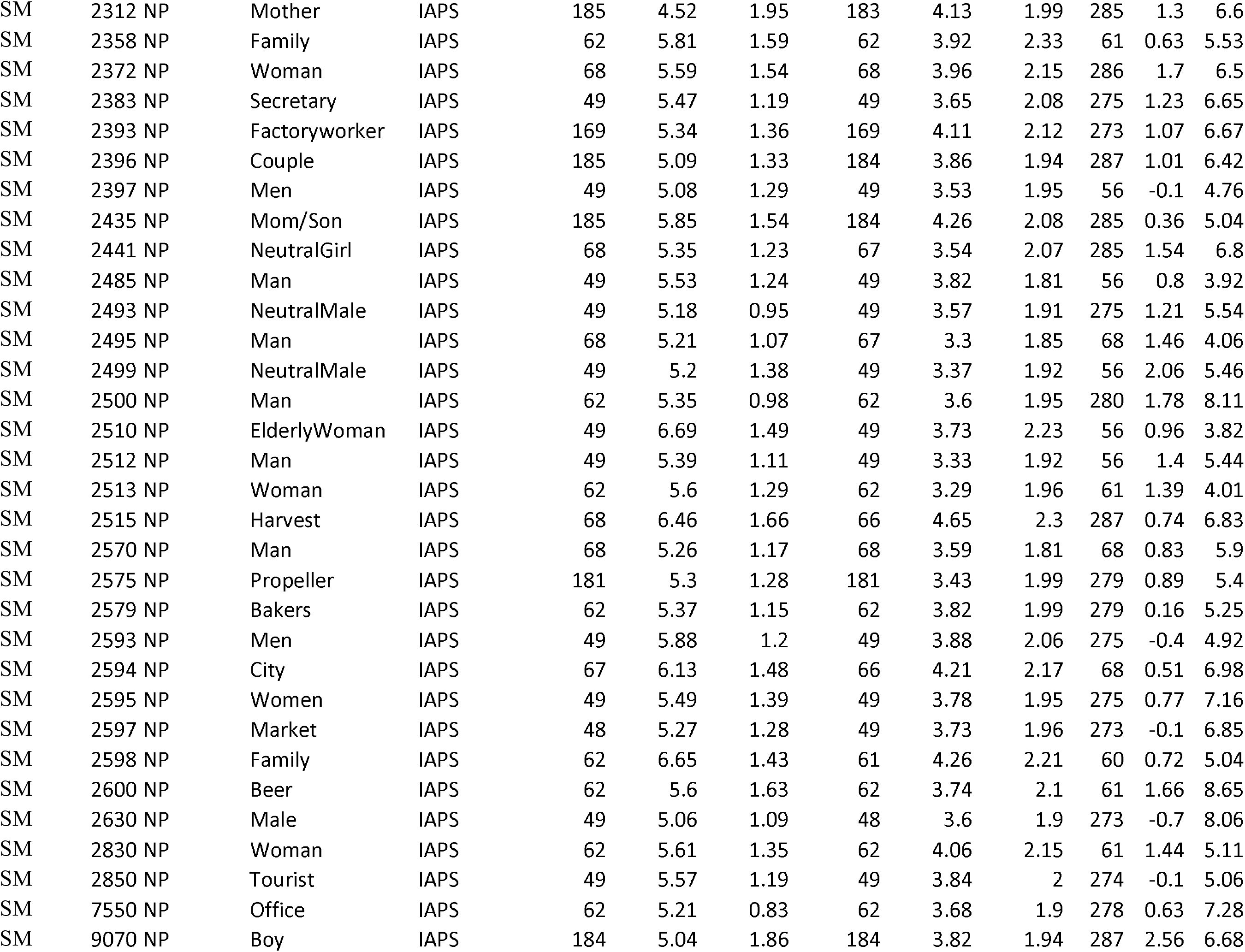
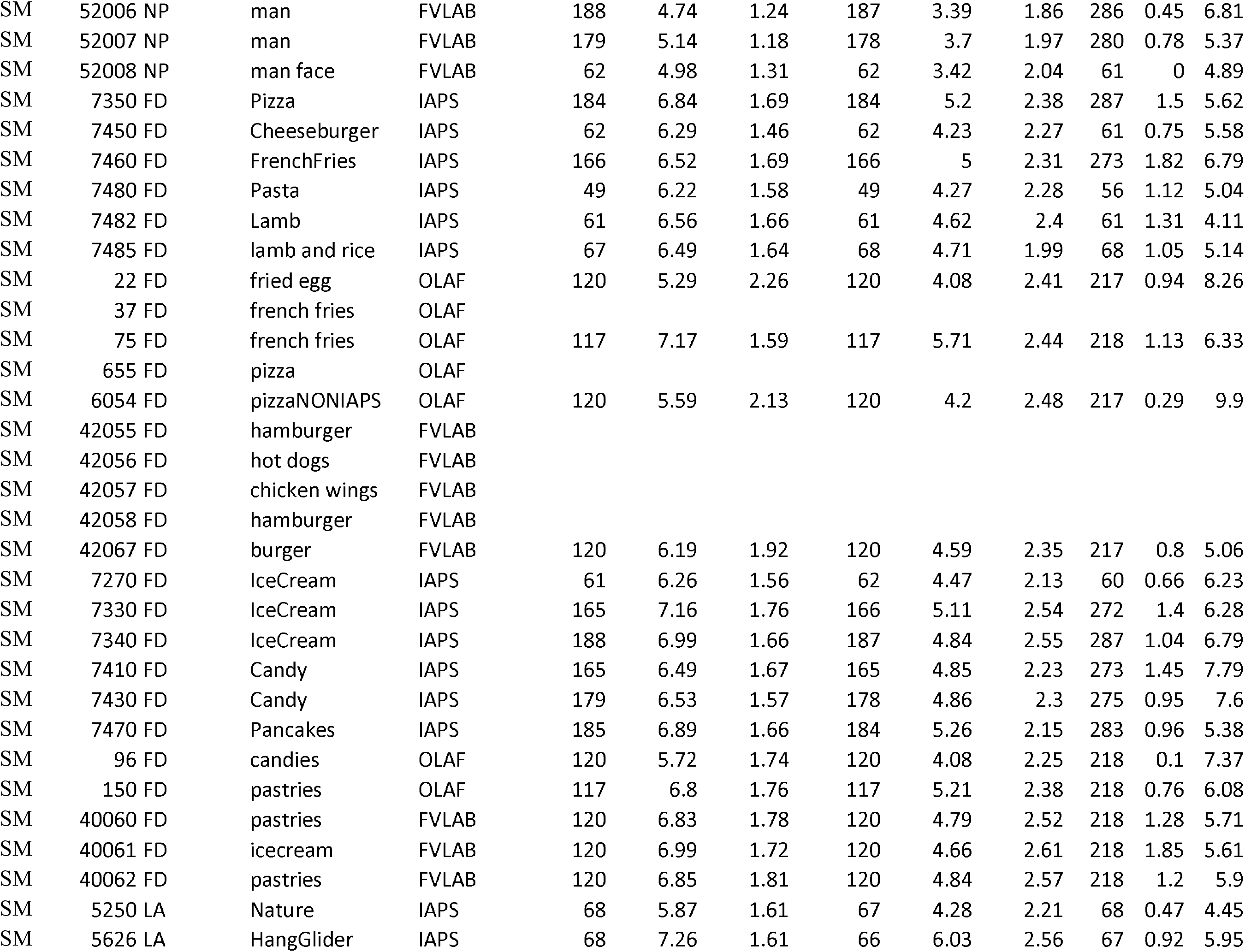
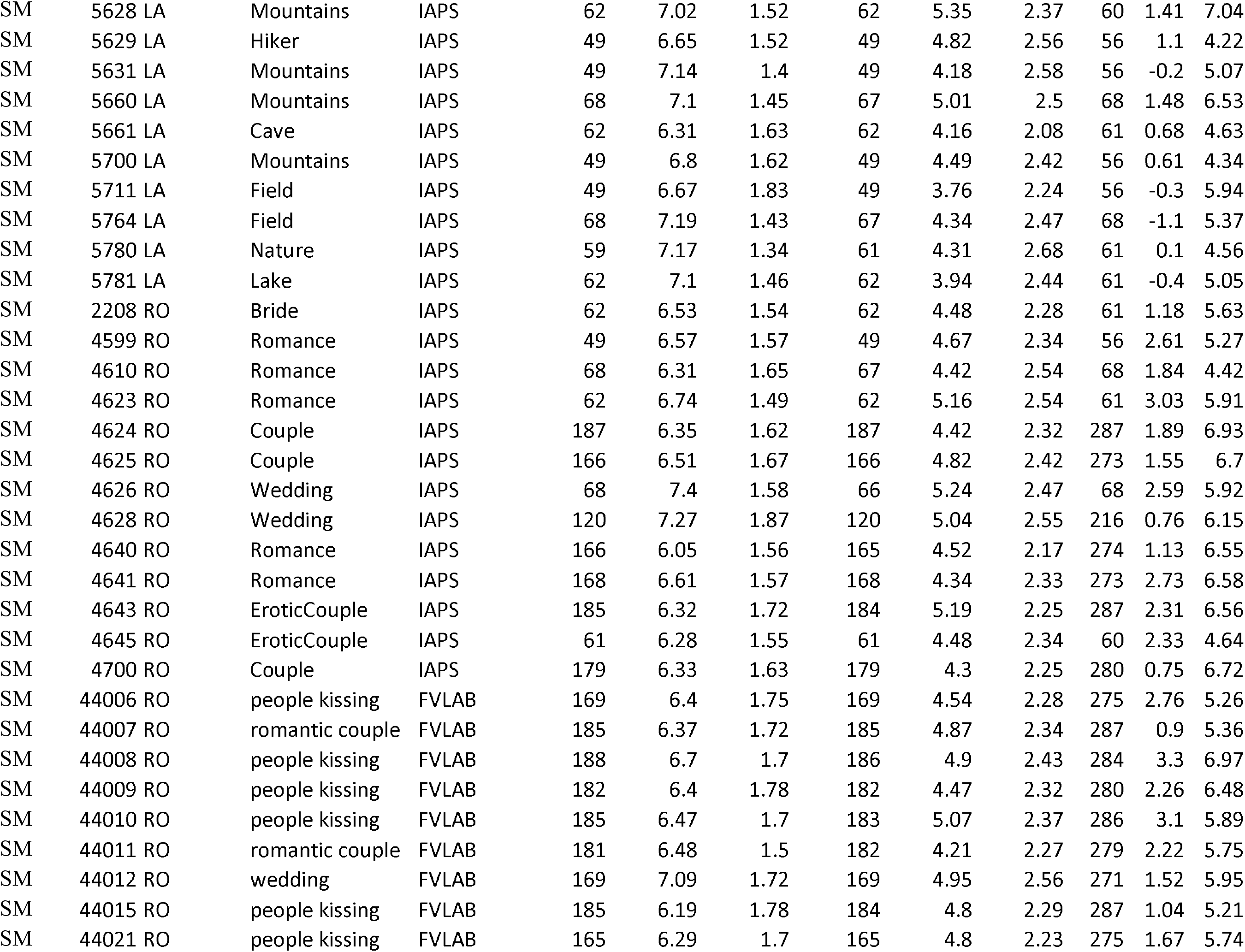
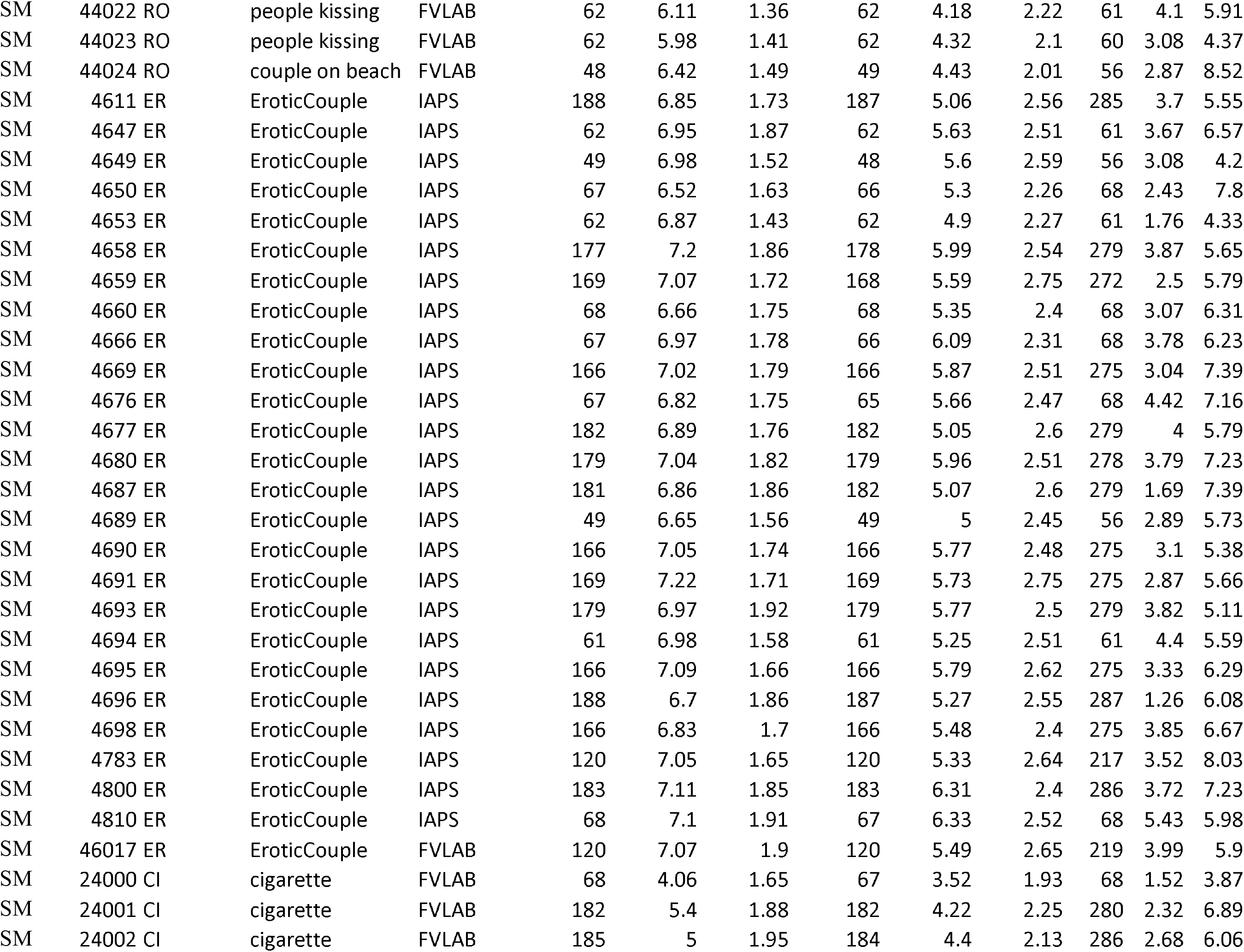
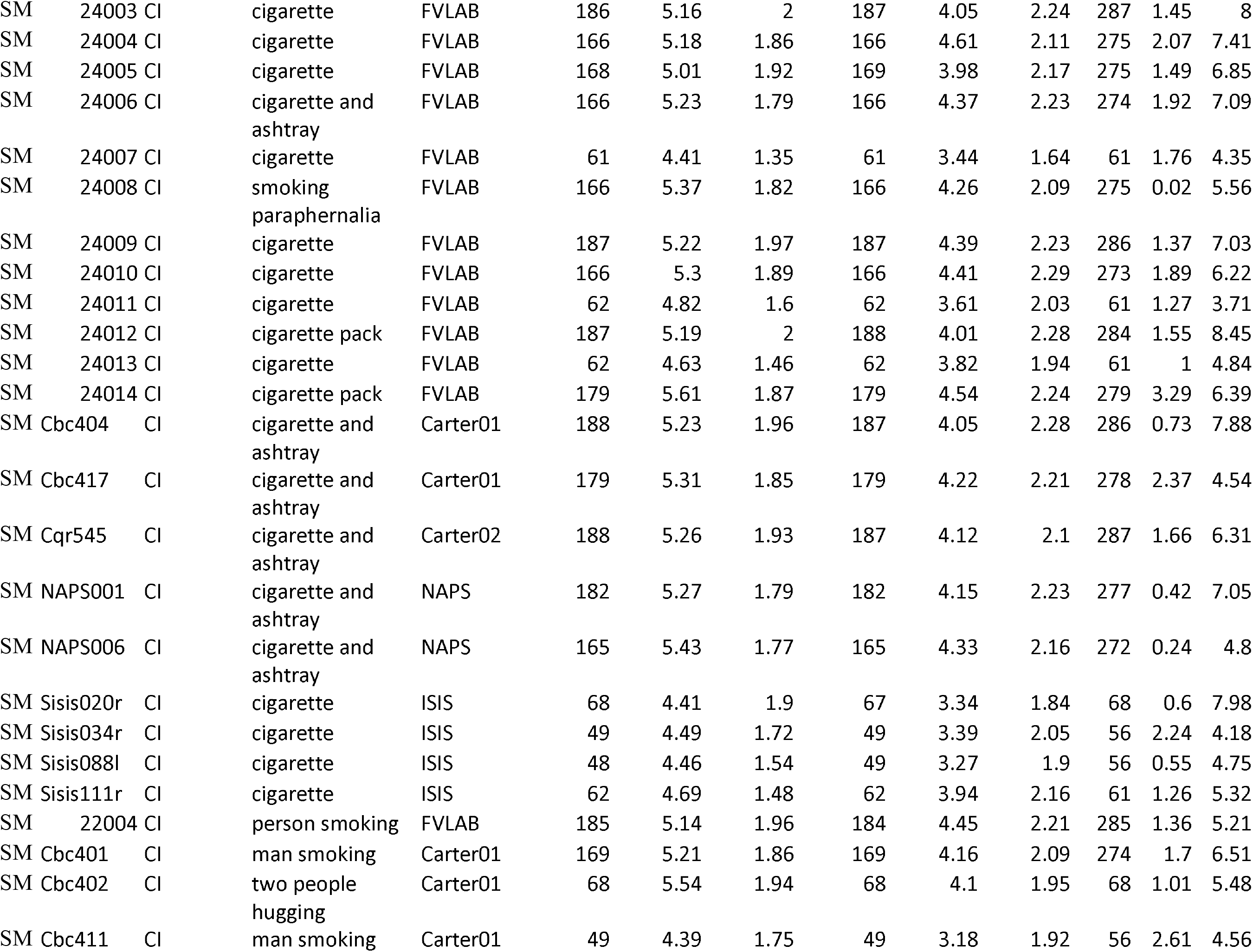
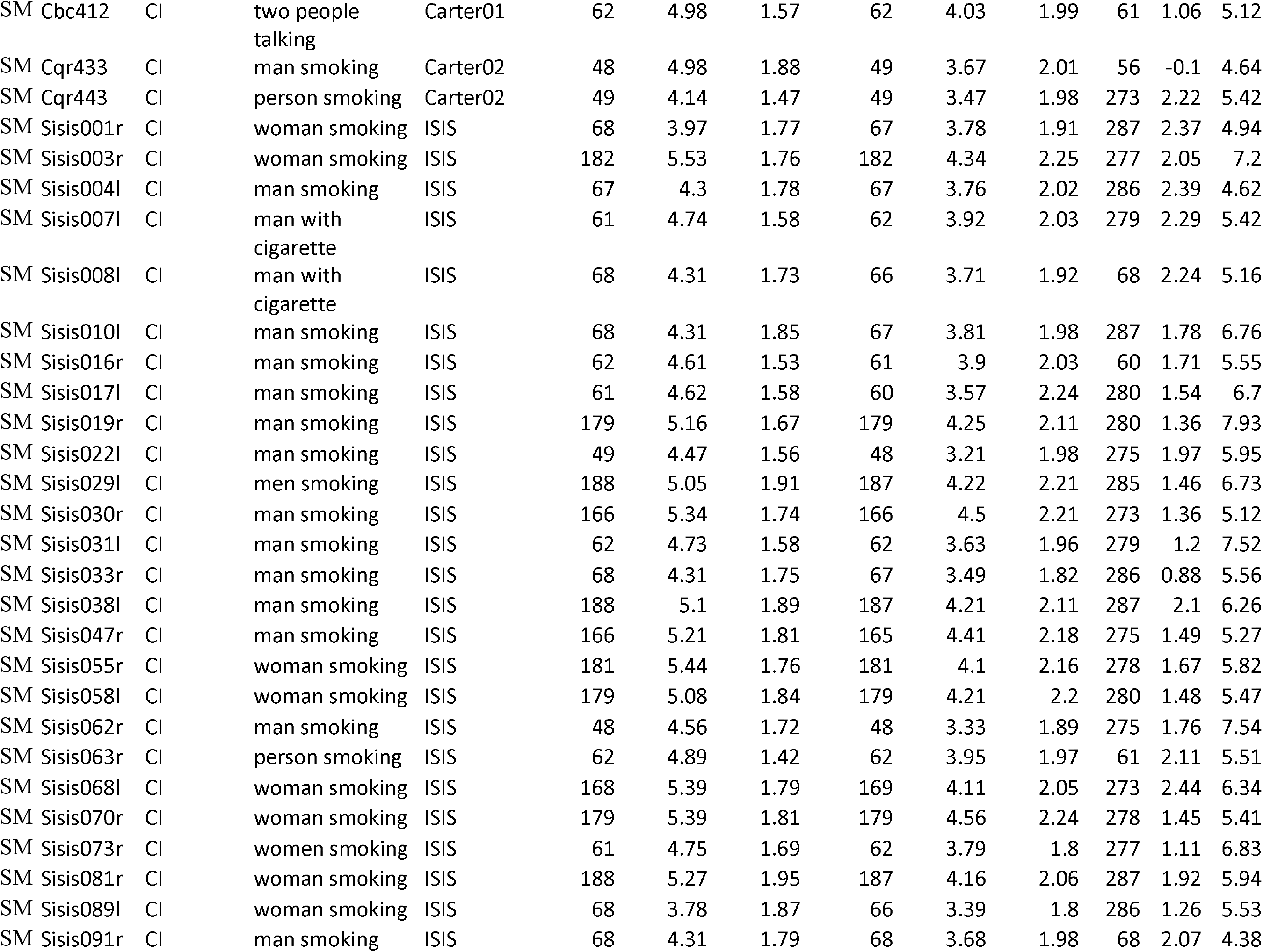

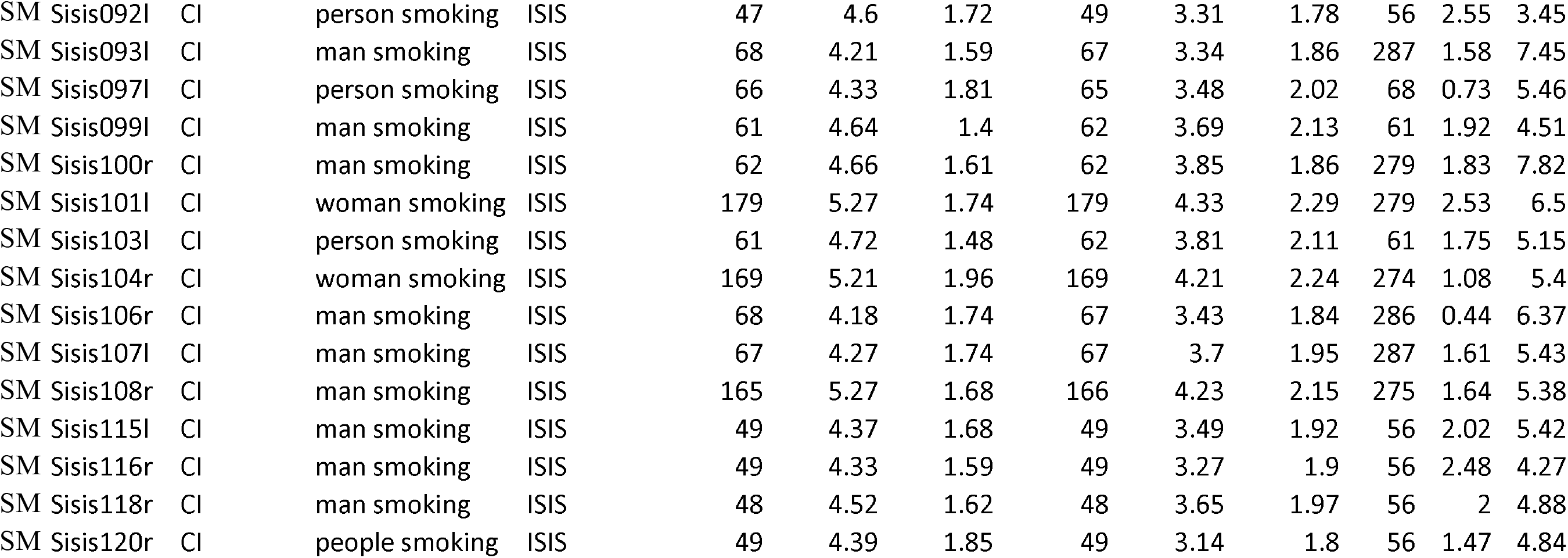

## Footnotes

1 One picture was rated by 15 participants. All other pictures were rated by at least 47 participants.

## REFERENCES

Blechert, J., Meule, A., Busch, N. A., & Ohla, K. (2014). Food-pics: An image database for experimental research on eating and appetite. Frontiers in Psychology, 5(JUN), 1–10. https://doi.org/10.3389/fpsyg.2014.00617

Bradley, M. M., Codispoti, M., Cuthbert, B. N., & Lang, P. J. (2001a). Emotion and motivation I: Defensive and appetitive reactions in picture processing. Emotion, 1(3), 276–298. https://doi.org/10.1037//1528-3542.1.3.276

Bradley, M. M., Codispoti, M., & Lang, P. J. (2006). A multi process account of startle modulation during affective perception. Psychophysiology, 43(5), 486–497. https://doi.org/10.1111/j.1469-8986.2006.00412.x

Bradley, M. M., Codispoti, M., Sabatinelli, D., & Lang, P. J. (2001b). Emotion and motivation II: Sex differences in picture processing. Emotion, 1(3), 300–319. https://doi.org/10.1037/1528-3542.1.3.300

Bradley, M. M., & Lang, P. J. (1994). Measuring emotion : the Self-Assessment Manikin and the semantic differential. J Behav Ther Exp Psychiatry, 25(I), 49–59.

Bradley, M. M., & Lang, P. J. (2007). The International Affective Picture System (IAPS) in the study of emotion and attention. In J. A. Coan & J. J. B. Allen (Eds.), Handbook of emotion elicitation and assessment (pp. 29–46). Oxford University Press.

Bradley, M. M, & Lang, P. J. (2018). Emotion in body & brain: Context-dependent action and reaction (pp. 280-282). In R. Davidson, A. Shackman, A. Fox, & R. Lapate (Eds.), The Nature of Emotion, 2nd Edition. Oxford University Press.

Bradley, M. M., Sapigao, R. G., & Lang, P. J. (2017). Sympathetic ANS modulation of pupil diameter in emotional scene perception: Effects of hedonic content, brightness, and contrast. Psychophysiology, 54(10), 1419–1435. https://doi.org/10.1111/psyp.12890

Carter, B. L., Robinson, J. D., Lam, C. Y., Wetter, D. W., Tsan, J. Y., Day, S. X., & Cinciripini, P. M. (2006). A psychometric evaluation of cigarette stimuli used in a cue reactivity study. Nicotine & Tobacco Research : Official Journal of the Society for Research on Nicotine and Tobacco, 8(3), 361–369. https://doi.org/10.1080/14622200600670215

Codispoti, M., De Cesarei, A., & Ferrari, V. (2012). Color and emotional perception. Psychophysiology, 49(1), 11–16. https://doi.org/10.1111/j.1469-8986.2011.01284.x

Codispoti, M., Ferrari, V., & Bradley, M. M. (2006). Repetitive picture processing: Autonomic and cortical correlates. Brain Research, 1068(1), 213–220. https://doi.org/10.1016/j.brainres.2005.11.009

Codispoti, M., Mazzetti, M., & Bradley, M. M. (2009). Unmasking emotion: Exposure duration and emotional engagement. Psychophysiology, 46(4), 731–738. https://doi.org/10.1111/j.1469-8986.2009.00804.x

Cofresí, R. U., Piasecki, T. M., Hajcak, G., & Bartholow, B. D. (2022). Internal consistency and test-retest reliability of the P3 event related potential (ERP) elicited by alcoholic and alcoholic beverage pictures. Psychophysiology, 59(2), e13967. https://doi.org/10.1111/psyp.13967

Cuthbert, B. N., Schupp, H. T., Bradley, M. M., Birbaumer, N., & Lang, P. J. (2000). Brain potentials in affective picture processing: Covariation with autonomic arousal and affective report. Biological Psychology, 52, 95–111. https://doi.org/ https://doi.org/10.1016/S0301-0511(99)00044-7

Deweese, M. M., Codispoti, M., Robinson, J. D., Cinciripini, P. M., & Versace, F. (2018). Cigarette cues capture attention of smokers and never-smokers, but for different reasons. Drug and Alcohol Dependence, 185, 50–57. https://doi.org/10.1016/j.drugalcdep.2017.12.010

De Cesarei, A., & Codispoti, M. (2011). Affective modulation of the LPP and α ERD during picture viewing. Psychophysiology, 48(10), 1397–1404. https://doi.org/10.1111/j.1469-8986.2011.01204.x

Ekhtiari, H., Kuplicki, R., Pruthi, A., & Paulus, M. (2020). Methamphetamine and Opioid Cue Database (MOCD): Development and validation. Drug and Alcohol Dependence, 209(November 2019), 107941. https://doi.org/10.1016/j.drugalcdep.2020.107941

Feldker, K., Heitmann, C. Y., Neumeister, P., Tupak, S. V., Schrammen, E., Moeck, R., Zwitserlood, P., Bruchmann, M., & Straube, T. (2017). Transdiagnostic brain responses to disorder-related threat across four psychiatric disorders. Psychological Medicine, 47(4), 730– 743. https://doi.org/10.1017/s0033291716002634

Ferrari, V., Bradley, M. M., Codispoti, M., & Lang, P. J. (2011). Repetitive exposure: Brain and reflex measures of emotion and attention. Psychophysiology, 48(4), 515–522. https://doi.org/10.1111/j.1469-8986.2010.01083.x

Frank, D. W., Cinciripini, P. M., Deweese, M. M., Karam-Hage, M. A., Kypriotakis, G., Lerman, C., Robinson, J. D., Tyndale, R. F., Vidrine, D. J., & Versace, F. (2020). Toward precision medicine for smoking cessation: Developing a neuroimaging-based classification algorithm to identify smokers at higher risk for relapse. Nicotine & Tobacco Research, 22(8), 1277– 1284. https://doi.org/10.1093/ntr/ntz211

Franken, I. H. A., Muris, P., Nijs, I., & Van Strien, J. W. (2008). Processing of pleasant information can be as fast and strong as unpleasant information: implications for the negativity bias. Netherlands Journal of Psychology, 64(4), 168–176.

Gilbert, D. G., & Rabinovich, N. E. (1999). The International Smoking Image Series (with Neutral Counterparts), V. 1.2. Department of Psychology, Southern Illinois University.

Henrich, J., Heine, S., & Norenzayan, A. (2010). The weirdest people in the world? Behavioral and Brain Sciences, 33(2-3), 61–83. https://doi.org/10.1017/S0140525X0999152X

Huffmeijer, R., Bakermans-Kranenburg, M.J., Alink, L.R.A., van IJzendoorn, M.H., 2014. Reliability of event-related potentials: the influence of number of trials and electrodes. Physiol. Behav. 130, 13–22. https://doi.org/10.1016/j.physbeh.2014.03.008

Ito, T. A., Larsen, J. T., Smith, N. K., & Cacioppo, J. T. (1998). Negative information weighs more heavily on the brain: The negativity bias in evaluative categorizations. Journal of Personality & Social Psychology, 75(4), 887–900. https://doi.org/10.1037//0022-3514.75.4.887

Kujawa, A., Proudfit, G. H., Kessel, E. M., Dyson, M., Olino, T., & Klein, D. N. (2015). Neural reactivity to monetary rewards and losses in childhood: Longitudinal and concurrent associations with observed and self-reported positive emotionality. Biological Psychology, 104, 41–47. https://doi.org/10.1016/j.biopsycho.2014.11.008

Lang, P. J. (1980). Behavioral treatment and bio-behavioral assessment: Computer applications. In J. B. Sidowski, J. H. Johnson, & T. A. Williams (Eds.), Technology in mental health care delivery systems (pp. 119–137). Norwood, NJ: Ablex.

Lang, P. J. (1988). What are the data of emotion? In V. Hamilton, G. H. Bower & N. Frijda (Eds.), Cognitive perspectives on emotion and motivation. Amsterdam: Martinus Nijhoff Publishers.

Lang, P. J. (2010). Emotion and motivation: Toward consensus definitions and a common research purpose. Emotion Review, 2(3), 229–233. https://doi.org/10.1177/1754073910361984

Lang, P. J., & Bradley, M. M. (2010). Emotion and the motivational brain. Biological Psychology, 84(3), 437–450. https://doi.org/10.1016/j.biopsycho.2009.10.007

Lang, P. J., Bradley, M. M., & Cuthbert, B. N. (2008). International affective picture system (IAPS): Affective ratings of pictures and instruction manual. Technical Report A-8. In Technical Report A-8. University of Florida. https://doi.org/10.1016/j.epsr.2006.03.016

Macatee, R. J., Carr, M., Afshar, K., & Preston, T. J. (2021). Development and validation of a cannabis cue stimulus set. Addictive Behaviors, 112(May 2020), 106643. https://doi.org/10.1016/j.addbeh.2020.106643

Manoliu, A., Haugg, A., Sladky, R., Hulka, L., Kirschner, M., Brühl, A. B., Seifritz, E., Quednow, B., Herdener, M., & Scharnowski, F. (2021). SmoCuDa: A Validated Smoking Cue Database to Reliably Induce Craving in Tobacco Use Disorder. European Addiction Research, 27(2), 107–114. https://doi.org/10.1159/000509758

Marchewka, A., Żurawski, Ł., Jednoróg, K., & Grabowska, A. (2014). The Nencki Affective Picture System (NAPS): Introduction to a novel, standardized, wide-range, high-quality, realistic picture database. Behavioral Research, 45, 596–610. https://doi.org/10.3758/s13428-013-0379-1

Moran, T.P., Jendrusina, A.A., Moser, J.S., 2013. The psychometric properties of the late positive potential during emotion processing and regulation. Brain Res. 1516, 66–75. https://doi.org/10.1016/j.brainres.2013.04.018

Miccoli, L., Delgado, R., Guerra, P., Versace, F., Rodríguez-Ruiz, S., Fernández-Santaella, M. C., & Rodrguez-Ruiz, S. (2016). Affective pictures and the open library of affective foods (OLAF): Tools to investigate emotions toward food in adults. PLoS ONE, 11(8), 1–13. https://doi.org/10.1371/journal.pone.0158991

Neumeister, P., Feldker, K., Heitmann, C. Y., Helmich, R., Gathmann, B., Becker, M. P. I., & Straube, T. (2016). Interpersonal violence in posttraumatic women: brain networks triggered by trauma-related pictures. Social Cognitive and Affective Neuroscience, 12(4), nsw165. https://doi.org/10.1093/scan/nsw165

Padmala, S., Sambuco, N., Codispoti, M., & Pessoa, L. (2018). Attentional Capture by Simultaneous Pleasant and Unpleasant Emotional Distractors. Emotion, 18(8), 1189–1194. https://doi.org/10.1037/emo0000401

Sabatinelli, D., Bradley, M. M., Fitzsimmons, J. R., & Lang, P. J. (2005). Parallel amygdala and inferotemporal activation reflect emotional intensity and fear relevance. NeuroImage, 24(4), 1265–1270. https://doi.org/10.1016/j.neuroimage.2004.12.015

Sambuco, N., Bradley, M. M., Herring, D. R., & Lang, P. J. (2020). Common circuit or paradigm shift? The functional brain in emotional scene perception and emotional imagery. Psychophysiology, 57(4). https://doi.org/10.1111/psyp.13522

Schupp, H. T., Cuthbert, B. N., Bradley, M. M., Cacioppo, J. T., Ito, T., & Lang, P. J. (2000). Affective picture processing: The late positive potential is modulated by motivational relevance. Psychophysiology, 37(2), 257–261. https://doi.org/10.1111/1469-8986.3720257

Schupp, H. T., & Kirmse, U. M. (2021). Case-by-case: Emotional stimulus significance and the modulation of the EPN and LPP. Psychophysiology, 58(4), 1–13. https://doi.org/10.1111/psyp.13766

Schupp, H. T., & Kirmse, U. (2022). Neural correlates of affective stimulus evaluation: a case-by-case analysis. Social Cognitive and Affective Neuroscience, 17(3), 300–310. https://doi.org/10.1093/scan/nsab095

Versace, F., Engelmann, J. M., Jackson, E. F., Costa, V. D., Robinson, J. D., Lam, C. Y., Minnix, J. A., Brown, V. L., Wetter, D. W., & Cinciripini, P. M. (2011). Do brain responses to emotional images and cigarette cues differ? An fMRI study in smokers. European Journal of Neuroscience, 34(12), 2054–2063. https://doi.org/10.1111/j.1460-9568.2011.07915.x

Versace, F., Frank, D. W., Stevens, E. M., Deweese, M. M., Guindani, M., & Schembre, S. M. (2019). The reality of “food porn”: Larger brain responses to food related cues than to erotic images predict cue induced eating. Psychophysiology, 56(4), e13309.https://doi.org/10.1111/psyp.13309

Versace, F., Kypriotakis, G., Basen-Engquist, K., & Schembre, S. S. M. (2016). Heterogeneity in brain reactivity to pleasant and food cues: evidence of sign-tracking in humans. Social Cognitive and Affective Neuroscience, 11(4), 604–611. https://doi.org/10.1093/scan/nsv143

Versace, F., Lam, C. Y., Engelmann, J. M., Robinson, J. D., Minnix, J. A., Brown, V. L., & Cinciripini, P. M. (2012). Beyond cue reactivity: Blunted brain responses to pleasant stimuli predict long-term smoking abstinence. Addiction Biology, 17(6), 991–1000. https://doi.org/10.1111/j.1369-1600.2011.00372.x

Taschereau-Dumouchel, V., Michel, M., Lau, H., Hofmann, S. G., & LeDoux, J. E. (2022). Putting the “mental” back in “mental disorders”: a perspective from research on fear and anxiety. Molecular Psychiatry, 1–9. https://doi.org/10.1038/s41380-021-01395-5

Weinberg, A., Correa, K. A., Stevens, E. S., & Shankman, S. A. (2021). The emotion elicited late positive potential is stable across five testing sessions. Psychophysiology, 58(11), e13904. https://doi.org/10.1111/psyp.13904

Weinberg, A., & Hajcak, G. (2010). Beyond Good and Evil: The Time-Course of Neural Activity Elicited by Specific Picture Content. Emotion, 10(6), 767–782. https://doi.org/10.1037/a0020242

Wessa, M., Kanske, P., Neumeister, P., Bode, K., Heissler, J., & Schonfelder, S. (2010). EmoPics: Subjektive und psychophysiologische Evaluation neuen Bildmaterials für die klinisch-biopsychologische Forschung. Zeitschrift Für Klinische Psychologie Und Psychotherapie, 39(11), 77.

